# Repurposing of a DNA segregation machinery into a cytoskeletal system controlling cell shape

**DOI:** 10.1101/2025.06.27.661541

**Authors:** Benjamin L. Springstein, Manjunath G. Javoor, Daniela Megrian, Roman Hajdu, Dustin M. Hanke, Bettina Zens, Gregor L. Weiss, Florian K. M. Schur, Martin Loose

**Author notes:** Corresponding authors: Benjamin L. Springstein and Martin Loose. These authors contributed equally to this work (Manjunath G. Javoor, Daniela Megrian).

## Abstract

Bacteria, like eukaryotes, use conserved cytoskeletal systems for intracellular organization. The plasmid-encoded ParMRC system forms actin-like filaments that segregate low-copy plasmids. In multicellular cyanobacteria like *Anabaena* sp., we found that a chromosomally-encoded ParMR has evolved into a novel cytoskeleton-termed CorMR-with a function in cell shape control rather than DNA segregation. Using live-cell imaging, *in vitro* reconstitution and Cryo-EM, we demonstrate that CorM forms dynamically unstable, antiparallel double-stranded filaments, which are recruited to the membrane by CorR via an amphipathic helix conserved in multicellular cyanobacteria. CorMR filaments are regulated by MinC, which excludes them from the poles and division plane. Comparative genomics reveal that the repurposing of ParMR and Min systems co-evolved with cyanobacterial multicellularity, highlighting the evolutionary plasticity of cytoskeletal systems in bacteria.

## Main Text

Bacteria employ different polymeric protein systems to control cell shape, growth and division (*1*). Protein filaments homologous to the eukaryotic cytoskeleton segregate plasmid DNA (ParM) (*2*), control cell division (FtsZ and FtsA) (*3–5*), or mediate cell elongation (MreB) (*6*), as also reviewed in (*1*). In *E. coli* and many other rod-shaped species, a spatiotemporal oscillation of the Min system generates an inhibitory gradient that restricts FtsZ polymerization to the midcell (*7*). While these discoveries provided profound insights into how bacterial cells are organized, prokaryotes exhibit immense diversity, and we are only beginning to grasp the full spectrum of their organizational strategies and corresponding evolutionary histories (*8*).

This is particularly true for Cyanobacteria, one of the most morphologically complex phyla described to date (*9*). Especially multicellular cyanobacteria have highly diverse cell morphologies, some of which are capable of cellular differentiation (*10*). Among them, cyanobacteria of the order *Nostocales*, including the model organism *Anabaena* sp. PCC 7120 (hereafter *Anabaena*), form long multicellular filaments (termed trichomes) and can differentiate heterocysts - specialized cells capable of nitrogen fixation (*11*). Multicellularity in cyanobacteria arises through incomplete cell division, resulting in a shared outer membrane, a continuous peptidoglycan (PG) layer and metabolically connected cytoplasms (*11*). Septal junctions - functional analogs to eukaryotic gap junctions - promote cell-cell communication through nanopores in the septal PG (*11–13*). The emergence of such complex, multicellular species likely requires distinct strategies to organize the intracellular space, to maintain cell morphology and to control cell division. However, so far, we know very little about the mechanisms that allow for the more complex intracellular organization of cyanobacteria as well as the evolutionary history of the biochemical systems involved.

For example, while cyanobacteria rely on FtsZ for cell division, they lack the membrane anchor FtsA (*14*, *15*), which is essential in Gram negative bacteria (*16*), and instead recruit FtsZ to the membrane by the cyanobacteria-specific protein ZipN (*17*). Despite the presence of thylakoid membranes, rod-shaped unicellular cyanobacteria like *Synechococcus elongatus* still rely on the oscillation of the Min system to control cell division as in *E. coli* (*18*). The spatial regulation of MinC’s inhibitory effect on Z-ring formation, however, depends on the additional, cyanobacteria-specific protein – Cdv3 – to recruit MinC to midcell (*18*). MreB is crucial for preserving the barrel-like shape of *Anabaena* cells, but unlike in other rod-shaped bacteria, it is not essential for their viability (*19–21*). MreB is also important for maintaining cell–cell communication between vegetative cells and heterocysts (*20*, *22*).

Cyanobacteria are also remarkably diverse in chromosome copy numbers, ranging from one to over 200 copies per cell (*23*), suggesting different strategies to distribute the genetic information to their daughters. In *Anabaena*, which contains ∼6-8 chromosome copies, a dedicated segregation mechanism seems likely, but none has been described to date (*21*). In the context of a recent comparative analysis (*19*), we discovered a chromosomal ParMR system in *Anabaena* and other multicellular cyanobacteria. As ParMR systems typically ensure maintenance of low copy plasmids during bacterial cell division (*24*), we hypothesized that the chromosomal systems in multicellular cyanobacteria may have evolved to control segregation of chromosomes. Typically, the ParMR system consists of the actin-like ATPase ParM, the DNA-binding adaptor protein ParR, and the centromere-like DNA element *parC* (*24*, *25*). The system gets initiated upon binding of ParR (generally encoded in an operon with *parM*) to short, centromere-like *parC* elements on the plasmid, thereby forming ParRC complexes. *E. coli* ParM forms parallel (i.e., polar), helical double-stranded filaments (a filament being composed of two protofilaments), which undergo cycles of polymerization and catastrophic depolymerization (*25–28*). Through this dynamic instability, ParM searches and captures ParRC complexes, which then stabilizes one filament end. Antiparallel alignment of two ParRC-bound, parallel ParM double-filaments gives rise to a cytoskeletal structure analogous to the mitotic spindle in eukaryotic cells, pushing newly replicated plasmids to opposite cell poles by incorporating new ParM subunits at the ParRC-bound filament ends (*24*, *25*, *27*). Whether the chromosomal ParMR systems of multicellular cyanobacteria segregate chromosomes has not been shown so far.

Here, we set out to understand the function of the chromosome-encoded ParMR system in *Anabaena*. To our surprise, we found that this system - and those of other multicellular cyanobacteria - have undergone extensive evolutionary diversification such that they now regulate cell shape and that their polymerization is under the spatiotemporal control of the Min system. This functional repurposing of existing biochemical systems – from DNA segregation to cell shape control – reveals an unexpected evolutionary innovation in bacterial intracellular organization coinciding with the emergence of multicellularity.

## Results

### A chromosomal ParMR system not involved in DNA segregation

While looking for proteins involved in division, growth and cell-cell communication in multicellular cyanobacteria (*19*), we discovered a chromosome-encoded ParMR system in *Anabaena* (which we first called chrParMR; accession numbers: WP_044522477.1 (chrParM) and WP_010999216.1 (chrParR); Fig. 1A). This finding was surprising, as ParMR systems have until recently (*29*) only been described on plasmids, with a single complete chromosomal ParMR system reported in *K. pneumoniae* (*30*) and few chromosomal ParR homologs in selected *Enterobacteriaceae* species (*31*) - though the function of these chromosomal systems remains unknown. *Anabaena* has about 8 chromosome copies and how they are maintained during cell division remains unknown (*21*, *23*). We therefore speculated that the chrParMR system may function as a chromosome segregation machinery.

**Fig. 1.**
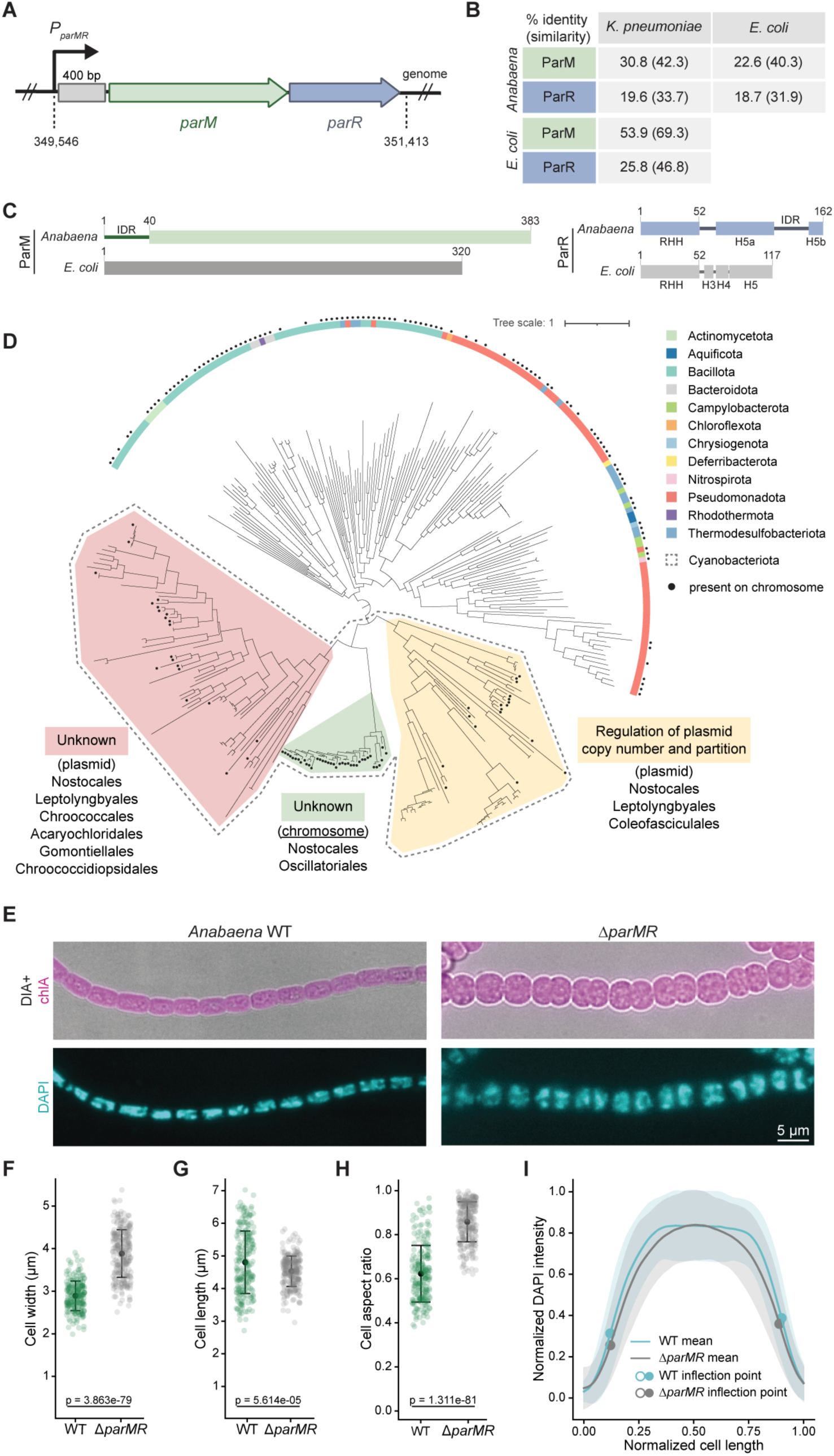
A ParMR system functioning in cell shape control. (**A**) Schematic representation of the chromosomal *parMR* operon from *Anabaena*. The promoter region is predicted to be about 400 bp long. Numbers below indicate genomic locus position. (**B**) Sequence identity and similarity (in brackets) in percent of chrParMR from *Anabaena* with chromosome-encoded KpParMR from *K. pneumoniae* and plasmid-encoded ParMR from *E. coli* R1 plasmid. (**C**) Schematic representation of the chromosome and plasmid-encoded ParM and ParR proteins from *Anabaena* and *E. coli*, respectively. chrParM contains a N-terminal 40 amino acid (aa) long intrinsically disordered region (IDR) in addition to globular domain observed in both, chrParM and EcParM. chrParR is longer than EcParR and lacks helix H3 and H4. Instead, helix H5 appears to be split into one large helix (H5a) and one small helix (H5b). Numbers indicate aa positions. (**D**) Maximum-likelihood phylogeny of bacterial ParM. All cyanobacterial sequences (outlined with a dashed line) form a monophyletic clade. Chromosome-encoded sequences are marked with a green background for Cyanobacteria, and with a black dot for other bacterial phyla. The scale bar represents the average number of substitutions per site. For the detailed tree, see Source Data. ParM proteins from the yellow clade structurally resemble previously characterized plasmid segregation systems, whereas the functions of ParM proteins from the green and pink clades remain unknown. (**E**) Micrographs of *Anabaena* WT and Δ*parMR* cells showing an overlay of Direct Illumination Acquisition (DIA) with chlorophyll a (chlA) autofluorescence images and DAPI staining from the same cells. (**F-H**) Scatter plots showing the mean cell property values ± standard deviation (SD) for (**F**) cell width (WT: 4.8 ± 0.96 µm; Δ*parMR*: 4.53 ± 0.47 µm), (**G**) cell length (WT: 2.89 ± 0.35 µm; Δ*parMR*: 3.88 ± 0.56 µm) and (**H**) cell aspect ratio (WT: 0.62 ± 0.13; Δ*parMR*: 0.86 ± 0.09) of *Anabaena* WT and Δ*parMR* cells. WT: n=222 cells; Δ*parMR*: n=259 cells. (**I**) Mean DAPI fluorescence intensity from individual cells plotted against the normalized cell length (0 = one cell pole, 1 = opposite cell pole). Values for WT are shown in cyan and values for Δ*parMR* are shown in grey. Shaded areas correspond to the SD around the mean. The intensity profiles show a peak near the center of the cell, indicating midcell enrichment of DAPI signal, with reduced signal toward the poles. WT: n=46 cells; Δ*parMR*: n=49 cells.

Compared to other plasmid-encoded ParMR such as the *E. coli* ParMR (EcParMR) from the R1 plasmid (*32*), the chrParMR of *Anabaena* does not have *parC*-like DNA repeats (*33–35*). In addition, the proteins are longer and share only limited sequence identity of 20-30% with the other ParMR proteins, contrasting the relatively high sequence identity of the plasmid-encoded EcParM with the chromosomal *K. pneumoniae* ParM (KpParM) (∼54 %) (Fig. 1B). Structure predictions indicated that chrParM contains an extensive N-terminal intrinsically disordered region (IDR), which is absent in EcParM (Fig. S1A) and KpParMR. Both chrParR and EcParR are ribbon-helix-helix (RHH) domain containing proteins (Fig 1C, S1B), which are typically dimers and bind DNA (*36*). chrParR is predicted to lack helices H3 and H4, which are characteristic for canonical ParR proteins, while helix H5 appears to be divided into two segments - a long H5a and a short H5b segment - connected by an IDR that is not present in EcParR (Fig 1C, S1B). Notably, in *E. coli*, helix H5 mediates ParMR interactions (*37*).

We first conducted a phylogenetic analysis to identify other chromosomal ParMR systems within cyanobacteria and other bacterial species. As the sequence conservation among ParR homologs (Fig. 1B) is low we performed this analysis looking for ParM homologs by sequence similarity and identifying ParR homologs based on their genomic context and 3D structure prediction. We limited our search to fully sequenced species with closed chromosomes and plasmids to unambiguously distinguish plasmid-encoded from chromosomal ParMR systems. The phylogeny reconstructed from the identified ParM protein sequences (Fig. 1D) shows that all cyanobacterial sequences are monophyletic, suggesting a common origin. However, we observed three distinct clades of ParMR systems in Cyanobacteria (Fig. 1D; Supplementary File 1), including two evolutionarily distant clades whose proteins are mostly plasmid-encoded (yellow and salmon). Of these only the ParMR proteins from the yellow clade structurally resembled previously described plasmid segregation systems, based on their 3D structure prediction and automatic annotation (*31*, *38*). One well-separated clade containing only chromosomal ParMR systems (green) is restricted to the orders *Nostocales* (including *Anabaena*) and *Oscillatoriales*. Within our representative sampling of 148 fully assembled cyanobacterial genomes, we found a chromosomal ParMR system in every *Nostocales* and *Oscillatoriales* genus analyzed, suggesting that its evolutionary emergence likely traces back to an ancestor of these two multicellular cyanobacterial orders. Notably, extended N-terminal IDRs were restricted to the green clade of chromosomal ParMs and are largely absent in the plasmid-encoded clades (Fig. S1C). ParM sequences identified in other bacterial phyla do not show monophyly within phyla, suggesting horizontal gene transfer between species. Surprisingly, we found chromosomal ParMR systems in a broad spectrum of bacterial species, challenging the notion of ParMR as a sole plasmid segregation system. Chromosomal ParM sequences appear scattered throughout the phylogenetic tree and are widely distributed. Indeed, when we searched for proteins identical to *E. coli* ParM from the R1 plasmid (WP_000959884.1) in all *E. coli* assemblies available in the NCBI RefSeq Genome database, we found 365 occurrences in plasmids and 12 in chromosomes, indicating that such exchanges occur even among different strains of the same species. When present in the chromosome, we identified a transposase gene (WP_001067858.1) in the same genomic context as *parM*, supporting the incorporation of the gene from a mobile element.

Together, these findings suggest that during bacterial evolution, the ParMR system has likely been exchanged between bacterial chromosomes and plasmids multiple times undergoing both vertical and horizontal transfers between species. In contrast, our results indicate that the evolutionary history of the ParMR systems in Cyanobacteria is more constrained, with two ancestral duplications and an inferred event of chromosomal integration that has been vertically inherited from an ancestor of *Nostocales* and *Oscillatoriales*.

To test the hypothesis that chrParMR is involved in chromosome segregation in *Anabaena*, we generated a Δ*parMR* mutant and assessed general DNA distribution and segregation using DAPI staining. Contrary to our initial hypothesis, we observed no obvious defects in cellular DNA distribution or any anuclear cells (Fig. 1E, I) as are commonly observed in strains deficient in other types of chromosome segregation systems (*39–41*). Instead, the mutant showed a striking change in cell morphology, losing the characteristic barrel-like shape of the wild type (WT) and forming enlarged and rounded cells instead (Fig. 1E). This phenotype was consistent across three independently obtained and verified Δ*parMR* mutant clones. Δ*parMR* cells displayed a near 1:1 aspect ratio, with width (3.88 ± 0.56 µm) approximately equal to length (4.53 ± 0.47 µm), contrasting WT cells (length: 4.8 ± 0.96 µm, width: 2.89 ± 0.35 µm) (Fig. 1F-H). Scanning electron microscopy (SEM) of Δ*parMR* cells (Fig. S1D) also showed possible defects in septum maturation or completion, indicating a potential role in cell-cell connectivity or communication. A slight reduction in fitness under standard growth conditions was also observed (Fig. S1E). Overall, the Δ*parMR* phenotype resembles a previously described *Anabaena ΔmreB* strain (*22*), suggesting a similar function in cell shape control. Complementation of the mutant with a replicative plasmid expressing chrParMR from the medium-to-strong copper-inducible P_petE_ promoter (*42*) produced cells with normal and aberrant shapes within a filament (Fig. S1F), potentially due to the high degree of cell-to-cell variation in plasmid copy number - and consequently in expression levels and the extent of complementation - a phenomenon well described in multicellular cyanobacteria (*43*, *44*). We could not recapitulate the native chr*parMR* expression levels through heterologous expression since the native chr*parMR* promoter (P_parMR_) is also employed by *E. coli* and strains carrying plasmids with P_parMR_-chr*parM*-chr*parR* could not be generated, likely due to the toxic effects of chrParMR expression (Fig. S1G). Based on these observations, we conclude that the *Anabaena* chrParMR system represents a unique variant of the ParMR family that plays a role in cell shape control, rather than in DNA segregation.

### Anabaena’s chrParM forms dynamically instable cortical filaments in vivo

To investigate the subcellular localization of chrParM in *Anabaena*, we used a CRISPR-Cpf1 (*45*, *46*) system to generate a strain in which the 5’ end of chr*parM* is translationally fused to mNeonGreen (mNG), resulting in sole expression of the tagged protein. We note that the mNG-chrParM fusion protein was not fully functional, as cells displayed an intermediate phenotype between WT and Δ*parMR* (Fig. 2A, S2A). In addition, ectopic expression of the mNG-chrParM-chrParR construct from the medium-to-strong *petE* promoter (P_petE_) (*42*) on a replicative plasmid in the Δ*parMR* background resulted in limited complementation, where cells with a detectable mNG-chrParM signal appeared slightly more WT-like (Fig. S2B, C). Attempts to generate a functional chromosomal translational C-terminal chr*parM-mNG* fusion were unsuccessful, likely due to the presence of the chr*parR* ribosomal binding site within the 3’ end of chr*parM*, consistent with the absence of reported C-terminal fluorescent EcParM fusions (*47*).

**Fig. 2.**
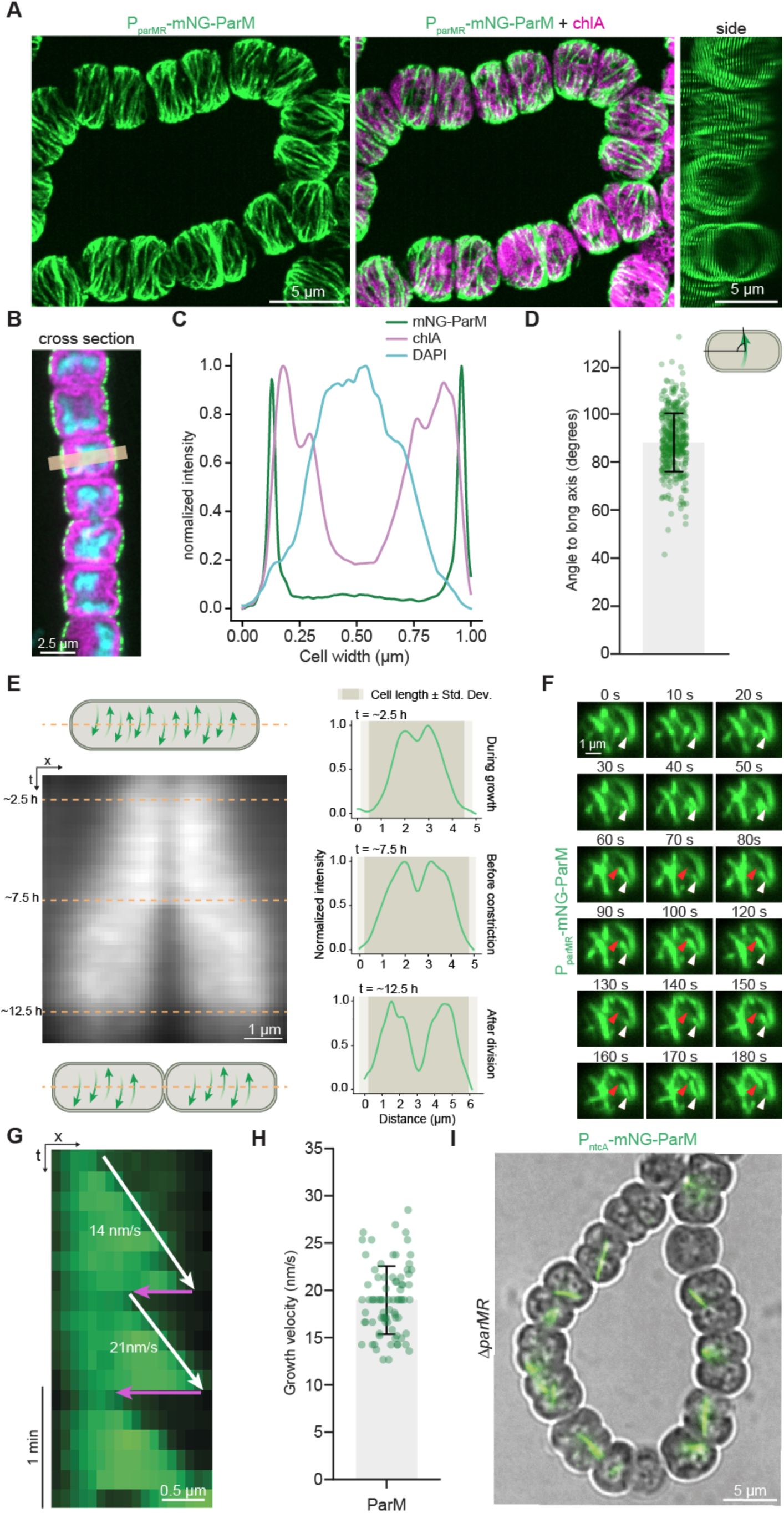
CorM forms cortical filaments transverse to the long cell axis. (**A**) Maximum intensity projection micrographs of *Anabaena* cells expressing mNG-ParM from its native locus as the sole copy within the cell. Left: mNG-ParM fluorescence signal, middle: overlap of mNG-ParM and chlorophyll A (chlA) autofluorescence (magenta), right: side view of the left image. (**B**) Cross section view of the same strain as in (**A**) showing the middle plane of the cells, highlighting that mNG-ParM and chlA signals do not overlap. Beige bar indicates region used to measure the intensity plot profile from mNG-ParM and chlA signal as shown in (**C**). (**D**) Angles of mNG-ParM filaments (n=386 filaments) in respect to the long cell axis. Cartoon indicates process of angle measurement. (**E**) Dynamic localization of mNG-ParM during the cell division cycle (see Supplementary Movie 1). The kymograph (center) shows the mean fluorescence intensity of mNG-ParM over time from 56 dividing cells. Top and bottom cell sketches indicate the chrParM patterns before and after cell division, respectively. Horizontal dashed lines indicate specific time points (∼2.5 h, ∼7.5 h, ∼12.5 h), corresponding to intensity profiles shown on the right. These profiles display the average distribution of mNG-ParM intensities at three key division stages: during growth, before constriction, and after division. Each line profile is the average signal from 3 micrographs taken within a 45-minute time window (images were taken every 15 min). Intensity profiles correspond the normalized mean fluorescence intensity signal from mNG-ParM, while shaded purple regions indicate the cell boundaries. The SD of the cell boundaries is shaded in light purple (n=56). (**F**) Time series from Supplementary Movie 2 displaying TIRF micrographs of individual mNG-ParM filaments over a period of 180 seconds. White triangle marks the initial position of a mNG-ParM filament, and the red triangle indicates its furthest observed position during the time series. (**G**) Kymograph of individual mNG-ParM filaments from TIRF microscopy. White arrows indicate growing filaments whereas magenta arrows indicate filament disassembly/shrinkage. (**H**) Scatter plot showing the growth velocities (mean 18.96 ± 3.61 nm/s) of mNG-ParM filaments as recorded by TIRF microscopy. Error bars represent the mean ± SD of 80 independent filaments measured. (**I**) Micrograph showing merged DIA and mNG-ParM fluorescence images of Δ*parMR* carrying a low copy number plasmid expressing mNG-ParM from a synthetic *ntcA* promoter integrated into the neutral *thrS2* site on the *Anabaena* chromosome (*48*) resulting in medium expression levels. Unlike in the WT, mNG-ParM loses its transverse localization in the absence of ParR.

Unlike EcParM, which forms cytoplasmic filament bundles (*2*), fluorescence imaging of live cells showed that chrParM assembled into filament arrays along the inner cell membrane, with no association to the intracellular thylakoid membranes or the cellular DNA (Fig. 2A-C). These filaments aligned perpendicular to the cell’s long axis, with a mean angle of 88.3 ± 12.15° (Fig. 2D). Although the chromosomal *mNG-*chr*parM* was not fully functional, multiple lines of evidence support the cortical localization of chrParM: (i) alternative fluorescent fusions expressed from the neutral *thrS2* locus on the chromosome (*48*) or from replicative plasmids retain WT-like morphology and show consistent cortical localization (Fig. S2D-F) and (ii) immunofluorescence microscopy of WT cells using an anti-chrParM antibody confirmed a similar pattern (Fig. S2G, H). Time lapse microscopy of dividing cells revealed a cell cycle-dependent distribution of chrParM filaments. During cell elongation chrParM is more and more excluded from the cell poles and the future division sites. Right after division chrParM is completely absent from the new cell poles (Fig. 2E; Supplementary Movie 1), suggesting active mechanisms limiting its cellular distribution. Using total internal reflection fluorescence (TIRF) microscopy, we then imaged the polymerization dynamics of chrParM in living cell. Consistent with previously described plasmid-encoded ParMs (*47*, *49*), chrParM exhibits dynamic instability *in vivo* (Fig. 2F, G; Supplementary Movie 2), with a filament growth rate of 18.96 ± 3.61 nm/s for mNG-chrParM (Fig. 2H), approximately half that of EcParM (*47*). Filaments also underwent rapid shrinkage at a rate of at least 52.06 ± 23.54 nm/s, about one-fifth of the shrinkage rate observed in *E. coli* (*47*). Expressing an mNG fusion of a chrParM variant lacking the N-terminal IDR (i.e., the first 40 amino acids; ^Δ1-40^chrParM) as the sole gene copy resulted in more severe morphological defects than the strain expressing mNG-tagged full-length chrParM (Fig. S2A), indicating that the N-terminal IDR is essential for maintaining proper cell shape in *Anabaena*. While mNG-^Δ1-40^chrParM still predominantly localizes perpendicular to the long cell axis, we observed a generally less ordered mesh-like filament arrangement with occasional localizations to the division site (Fig. S3A). Interestingly, although dynamic instability was retained (Fig. S3B), the N-terminal deletion significantly increased the filament growth rate to 36.14 ± 8.04 nm/s (Fig. S3C). Thus, chrParM maintained the polymerizing properties of EcParM, including dynamic instability, but exhibits a vastly different cellular localization pattern with dynamic properties dependent on the N-terminal IDR absent in plasmid-encoded ParMs.

To assess the role of chrParR for chrParM localization, we ectopically expressed mNG-chrParM from a replicative plasmid in Δ*parMR*. Here, in the absence of chrParR, we observed mNG-chrParM filaments appearing in the cytoplasm of the cells without a clear localization pattern (Fig. 2I), suggesting that chrParR is essential for the organization of chrParM into a cortical array. Genomic replacement of chrParR with a C-terminally mNG-tagged variant (chrParR-mNG) as the sole gene copy confirmed its localization to the cell envelope (Fig. S3D). The accumulation of chrParR-mNG signal at the cell poles is likely an artifact resulting from the close proximity of two neighboring cell membranes at these sites, where signal from adjacent cells merges due to the resolution limits of fluorescence microscopy. However, the swollen and rounded phenotype of this strain indicates that the fusion protein is not fully functional, suggesting that an unmodified C-terminus is essential for chrParM binding similar to what was previously reported for EcParR (*31*, *37*). When heterologously expressed alone in *E. coli*, chrParM formed intracellular filaments leading to cell elongation (Fig. S4A). This phenotype was observed for both full-length chrParM and ^Δ1-40^chrParM (Fig. S4A). In contrast, chrParR C-terminally fused to mScarlet-I resulted in distinct accumulations at the cell poles and throughout the cell (Fig. S4B). Upon extended expression periods, these sites of accumulations were frequently associated with severe local distortions of cell shape (Fig. S4C). Short-term co-expression of both proteins in *E. coli* led to recruitment of chrParM to the sites of chrParR accumulation (Fig. S4D), which, upon extended expression periods, caused pronounced morphological defects and ultimately cell death (Fig. S1G). These heterologous expression results support a direct chrParM-chrParR interaction.

In summary, *Anabaena*’s chrParM and chrParR proteins form an evolutionarily distinct ParMR system, whose deletion results in a pronounced cell shape defect. Using fluorescent versions of the protein, we found that this actin-related protein assembles into a cortical array of membrane-associated filaments. Together, these findings strongly suggest that chrParMR is not involved in DNA segregation. According to their intracellular localization and cellular function, we therefore propose to rename chrParM and chrParR as CorM and CorR, respectively, to reflect their *<u>cor</u>tical* cytoskeletal organization.

### In vitro characterization of the CorMR system

To biochemically characterize CorM, we purified the protein via heterologous expression in *E. coli*. CorM retained ATPase activity but in contrast to EcParM, showed negligible GTPase activity (Fig. S5A), and was unable to polymerize in the presence of GTP and only weakly polymerized in the presence of AMP-PNP, a non-hydrolyzable ATP analog (Fig. S5B-D). ATP hydrolysis by CorM increased with protein concentration (Fig. S5E), while high-speed pelleting assays did not indicate a clear critical concentration for polymerization under the tested conditions (Fig. S5F). Interestingly, deletion of the N-terminal IDR (^Δ1–40^CorM) led to elevated ATPase activity (Fig. S5G), indicating that the IDR modulates polymerization kinetics.

We next confirmed interaction between CorM and CorR (Fig. S5H) in high-speed co-pelleting assays (Fig. S5I, J), as well as through bacterial two-hybrid (B2H) screens (Fig. S5K). Using CorR mutants, we confirmed that this interaction is mediated by CorR’s helix H5b (Fig. S5K), consistent with the *E. coli* ParMR system, In plasmid-based ParMR systems, DNA segregation depends on ParR binding to *parC* repeats (see for example (*28*, *31*, *50*)). To determine whether CorR also binds to DNA, we performed electrophoresis mobility shift assays (EMSA) using the promoter region of *corMR* (P_corMR_) and *E. coli parC* as bait. CorR failed to bind to the P_corMR_ DNA region even at high concentrations (Fig. 3A) and could also not bind to the *E. coli parC* element (Fig. S6A), whereas EcParR produced a clear mobility shift of *parC* (Fig. 3A).

**Fig. 3.**
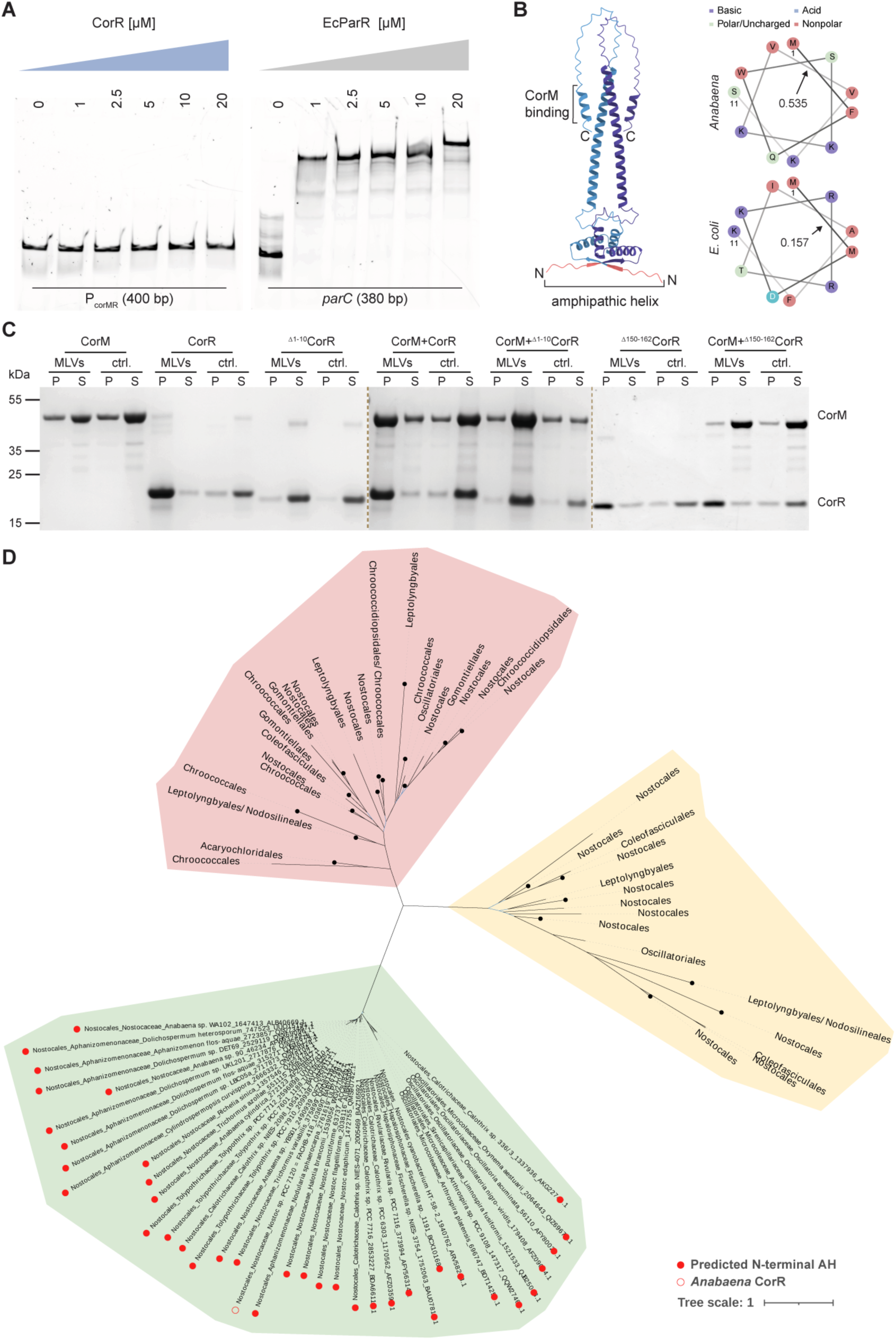
**CorR binds to lipid membranes and recruits CorM**. (**A**) EMSA assay with increasing concentrations of CorR or *E. coli* ParR using the promoter region of *corMR* (P_corMR_, 400 bp) or the promoter region of Ec*parMR* (termed *parC*; 380 bp), respectively, as bait DNA (150 ng/lane). Protein concentrations in µM are given above the individual lanes. The gel shows a representative example from six independent experiments, displaying Alexa488 fluorescence signals of PCR-amplified DNA regions obtained using fluorescently labeled primers. (**B**) Left shows an AlphaFold3 prediction of a CorR dimer with C-terminal CorM interaction site and N-terminal amphipathic helix (AH) indicated. The AH within the first 10 aa was identified using Amphipaseek (*51*). Notably, AlphaFold3 does not predict the N-terminal AH as a helical structure but rather as an unstructured region. This may be because AHs often exist as nascent helices that only adopt their proper fold upon contact with the membrane. Right shows the helical wheel projection of the first 11 aa of CorR and EcParR. Arrows indicate the hydrophobic moment of the predicted helix indicating the amphipathicity of the respective proteins. The high hydrophobic moment of CorR indicates the potential to bind to lipid membranes at the interface between the lipid bilayer and the cytosolic phase. (**C**) Representative Coomassie-stained SDS-PAGE gel of a pelleting assay with indicated protein combinations with or without MLVs. 30 % of the total pellet [P] and 12.5 % of the total supernatant [S] samples were separated on the gel. The respective quantifications are shown in Fig. S6B. CorR is enriched in the pellet fraction in the presence of MLVs. CorM alone is not efficiently pelleted at 21,000 x *g* but gets enriched in the pellet when co-incubated with CorR, indicating that CorR recruits CorM to the MLV-containing pellet. CorR lacking the N-terminal AH (^Δ1–10^CorR) fails to bind MLVs and is thus not enriched in the MLV-containing pellet fraction. In contrast, CorR^Δ150–162^ retains the N-terminal AH and can be recruited to the pellet via MLV interactions but cannot recruit CorM to the pellet due to the absence of the C-terminal peptide required for CorM binding. Dotted lines separate individual gels stitched together for better visualization. (**D**) Maximum-likelihood phylogeny based on the concatenation of CorM/ParM and CorR/ParR sequences from Cyanobacteria. The presence of a predicted AH in CorR/ParR cyanobacterial homologs is indicated with a red dot. *Anabaena* CorR is indicated with a red circle. The scale bar represents the average number of substitutions per site. For the detailed tree, see Source Data.

Sequence analysis using Amphipaseek (*51*) revealed an N-terminal amphipathic helix (AH) in CorR’s first 10 amino acids, located just upstream of the (RHH)_2_ domain (Fig. 3B). We confirmed membrane binding of CorR in vesicle co-sedimentation assays, where the protein was only found in the pellet in the presence of multilaminar vesicles (MLVs) (Fig. 3C, S6B). Furthermore, CorM co-sedimented with MLVs mostly only in the presence of CorR, confirming that CorR recruits CorM to lipid membranes. Although deletion of CorM’s N-terminal IDR did not impair its recruitment to the pellet by CorR, ^Δ1-40^CorM exhibited increased pelleting compared to the full-length CorM, suggesting that the N-terminal IDR contributes to filament length control or bundling (Fig. S6C, D). In contrast, deletion of CorR’s N-terminal AH (^Δ1-10^CorR) abolished CorR’s ability to bind MLVs and to recruit CorM (Fig. 3C, S6B). Meanwhile, a CorR variant lacking the C-terminal α-helix (amino acid residues 150-162; CorR^Δ150-162^) retained lipid-binding capacity but failed to recruit CorM (Fig. 3C, S6B), consistent with our B2H results (Fig. S5K). Together with our observation of cytoplasmic filaments of ectopically expressed mNG-CorM in Δ*corMR* (Fig. 2I), and the observation that Δ*corR* and Δ*corMR* mutants display the same phenotype (Fig. S7), these results identify CorR as the essential membrane anchor of CorM.

A Δ*corM* mutant could not be obtained, as attempts to construct a plasmid for homologous recombination replacing P_corMR_-*corM*-*corR* with P_corMR_-*corR* failed. This likely reflects CorR’s toxicity in *E. coli*, where the P_corMR_ promoter is active and would drive direct CorR expression (see Fig. S1G, S4C). To dissect the contribution of individual domains, we generated three additional mutants besides the original Δ*corMR* double mutant: ^Δ1-10^*corR* (lacking the AH required for membrane binding), *corR*^Δ150-162^ (lacking the C-terminal CorM-binding peptide), and ^Δ1-40^*corM* (lacking the N-terminal IDR of CorM involved in filament dynamics and bundling). All three mutants showed phenotypes essentially indistinguishable from Δ*corMR*, although ^Δ1-40^CorM showed some phenotypic variability (Fig. S7), confirming the essential nature of these domains for proper CorMR system function. Finally, we were wondering if other bacterial ParRs have acquired an N-terminal AH suggesting membrane-instead of DNA-binding. However, by analyzing the protein sequence of all cyanobacterial ParRs, we found that only CorR homologs from *Nostocales* and *Oscillatoriales* cyanobacteria (Fig 3D) contain an AH, with strong sequence conservation in *Nostocales* but not in *Oscillatoriales* species (Fig. S6E, S8). This suggests that the acquisition of *corR* in the chromosome and the emergence of this membrane binding motif have a common origin. Together, these findings provide the first evidence for a ParR homolog that has lost its DNA-binding capacity and instead evolved the ability to bind membranes, mediated by a newly acquired amphipathic helix at the N-terminus of the (RHH)_2_ domain.

### CorM assembles into antiparallel, double-stranded helical filaments

Given the unexpected behavior of the CorMR system, we aimed to gain structural insights into the molecular architecture of CorM filaments and their interaction with CorR. To this end, we performed single particle analysis cryo-electron microscopy (cryo-EM) of filaments consisting of solely CorM, ^Δ1-40^CorM or CorM assembled in the presence of CorR (CorMR) (Fig. 4A, S9.1-9.4).

**Fig. 4.**
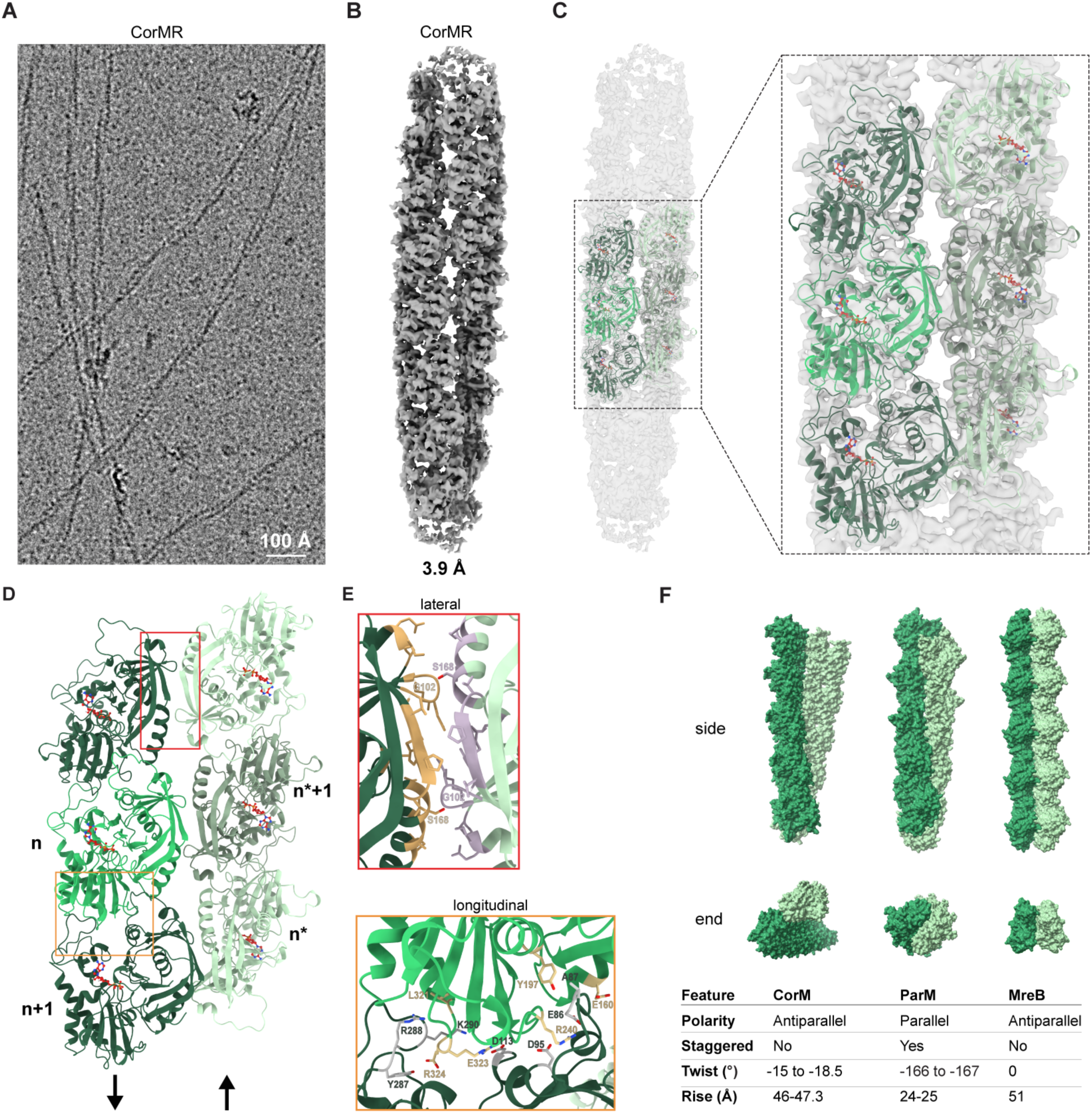
Cryo-EM structure of *in vitro* reconstituted CorM filaments. (**A**) Cryo-EM micrograph showing CorMR filaments formed in vitro at a protein concentration of 5 µM for each CorM and CorR. (**B**) Cryo-EM density map of *in vitro* reconstituted wild type CorM filament in the presence of CorR (CorMR) at 3.9 Å. The details of the cryo-EM processing pipeline for all the three obtained structures are provided in Fig. S9.1-S9.3. (**C-E**) Refined model of the CorM filament built by fitting the AlphaFold-predicted CorM monomer into CorM+CorR cryo-EM density map. ADP molecules are shown. (**D**) The two antiparallel protofilaments are annotated as n and n*. Rectangles outline the enlarged lateral and longitudinal interaction interfaces in (**E**). Both of the interface interactions were determined using PDBePISA (Proteins, Interfaces, Structure and Assemblies) (*52*). (**F**) Comparison of surface representations of CorM (this study), with previously published *E. coli* ParM (PDB entry 5AEY) and *C. crescentus* MreB (PDB entry 4CZJ) models, respectively. Individual protofilaments are colored in dark or light green to illustrate their relative arrangement. Similarities and differences in polarity, subunit arrangement and helical parameters are given below the models.

We included ^Δ1-40^CorM to investigate if the unusually long N-terminal part of CorM might play a structural role in filament assembly. For all three samples, we obtained electron microscopy density maps at resolutions that allowed unambiguous assignment of individual CorM subunits (Fig. 4B, C, S9.1-9.4), revealing in each case that CorM filaments consist of two antiparallel, unstaggered protofilaments forming a left-handed helix. Our structures also show that both deletion of the N-terminal amino acids as well as the presence of CorR in the *in vitro* assembly do not visibly alter filament architecture (Fig. S9.1-9.4). However, we observed that CorM alone forms shorter filaments in our *in vitro* preparation (Fig. S9.3) and that these filaments had a slightly differing rise and twist (46.5 Å/-18.5°) as compared to the other two structures (CorMR: 46.427 Å/-15.75°; ^Δ1-40^CorM 47.294 Å/-15.344°) (Fig. S9.1-9.2). Since the CorMR sample reached the highest resolution (3.9 Å as compared to 4.4 Å for ^Δ1-40^CorM and 7.1 Å for CorM alone), we used this electron microscopy density map to build a molecular model of the CorM filament (Fig. 4C, D, S9.4). Upon closer inspection of the maps fitted with the molecular model, we found that the 40 N-terminal amino acids do not adopt a structured conformation in either of the CorM or CorMR samples. Similarly, within the CorMR filament structure all visible density was occupied by CorM alone, which suggests that CorR only binds to the filament ends, as has been shown for other ParR systems (*25*). Given the non-staggered nature of the filament assembly, the lateral association of the two CorM protofilaments (designated n and n* in Fig. 4D) is established by interactions between a subunit in one strand with only one subunit in the other strand. Analysis of our model using PDBePISA (*52*) shows only weak hydrogen bond interactions across a small lateral interface (interface area 452 Å^2^) (Fig. 4E). Hence, it seems CorM forms weaker lateral interactions compared to EcParM, where multiple lateral interactions between the protofilaments are determined by salt bridges (*27*). This potentially has implications in filament stability or dynamic behavior of CorM. Further cryo-EM processing to obtain a higher resolution structure and a structure of filament ends will be required to further characterize the lateral interactions as well as how CorR interacts with CorM.

In contrast, the longitudinal interactions between CorM subunits along the filament are extensive (interface area 1466 Å^2^) and involve multiple salt bridges and hydrogen bonds (Fig. 4E), similar to EcParM filaments (*27*). Interestingly, EcParM filaments have been described to form parallel, polar and staggered helices, which is drastically different from the antiparallel, bipolar and non-staggered assembly of CorM (Fig. 4F). Instead, the filament architecture of CorM more closely resembles the antiparallel, bipolar filament doublets formed by MreB (Fig. 4F), although MreB does not adopt a twisted conformation and does not show dynamic instability. By using cryogenic electron tomography (cryo-ET) of focused ion beam (FIB)-milled *Anabaena* cells (WT and Δ*cse* mutant (*53*)), we identified cytoplasmic filaments positioned close to the inner membrane, with appearance and dimensions closely matching those of the *in vitro*-assembled CorM filaments (Fig. S10, Supplementary Movie 4, 5). Due to limitations in cryo-ET sample preparation and milling orientation, we predominantly observed cross-sections of these filaments in our tomograms. Nevertheless, by examining more than 100 high-quality tomograms, we identified a CorM filament oriented perpendicular to the tomogram’s z-axis at the edge of a septum (Fig. S10D, E; Supplementary Movie 5). Together, these observations support the native cortical localization of CorM filaments and reveal a novel filament architecture within the ParM family, underscoring the uniqueness of the CorMR system relative to plasmid-encoded, DNA-segregating ParMR systems.

### Reconstitution of CorM dynamics in vitro on supported lipid bilayers

We next asked whether CorM dynamics could be reconstituted on membranes *in vitro*. To this end, we used supported lipid bilayers (SLBs), as previously described for the cytoskeletal system formed by FtsZ and FtsA (*4*) (Fig. 5A). Indeed, we successfully recapitulated the *in vivo* behavior of CorM and CorR *in vitro* (Fig. 5B), demonstrating that these two proteins are sufficient to drive filament formation and dynamics.

**Fig. 5.**
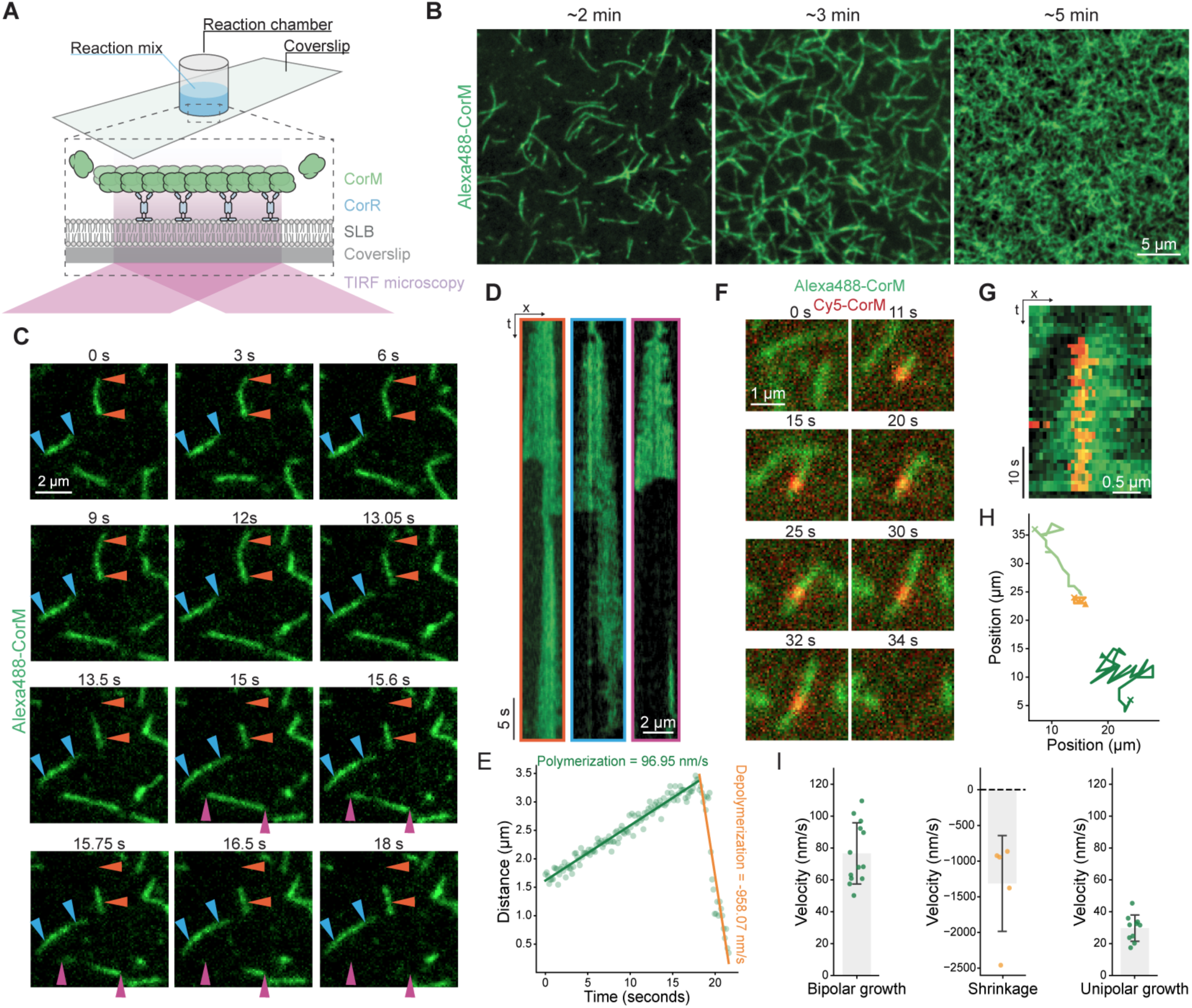
*In vitro* reconstitution of CorMR polymerization on supported membranes. (**A**) Illustration of the experimental assay to reconstitute CorM polymerization with CorR on supported lipid bilayers (SLBs). CorR binds to the SLB through its AH and recruits CorM filaments to the membrane. CorM filaments grow from both sites through bidirectional addition of monomers to an existing filament. (**B**) TIRF micrographs of purified and Alexa488-labeled CorM (0.25 µM) together with unlabeled CorM (0.75 µM) recruited to the SLB through interaction with CorR (0.25 µM). After induction of polymerization, CorM initially forms individual filaments (∼2 min) that then eventually form a denser network of filaments (∼3 min) and ultimately merge into a dense, seemingly highly interconnected meshwork of thin filaments (∼5 min). (**C**) Time series from Supplementary Movie 8 displaying TIRF micrographs of individual Alexa488-labeled CorM filaments showing dynamic instability. Blue triangles mark the initial position of an individual CorM filament that continues to grow throughout the whole time series. Orange triangles mark the initial position of an individual CorM filament that grows initially and then rapidly collapses before regrowing. The purple triangles indicate the furthest observed position of another filament which ultimately undergoes rapid and complete depolymerization during the time series. (**D**) Kymographs of the respective filaments marked with blue, orange and purple triangles in (C) showing growth, catastrophe and rescue events. The filament indicated with blue triangles only grows during the 120 sec time series shown in (C) and eventually also undergoes depolymerization. The filament indicated with purple triangles grows bidirectionally and then completely collapses. (**E**) CorM filament undergoing an initial polymerization phase followed by rapid depolymerization. Growth is represented by the increase in distance between the two poles of the filament over time. The filament polymerizes steadily at a rate of 96.95 nm/s (green line) followed by a rapid depolymerization phase with a rate of-958.07 nm/s (orange line). (**F**) Time series from Supplementary Movie 9 displaying TIRF micrographs of an individual Alexa488-labeled CorM filament supplemented with Cy5-CorM over a period of 32 seconds. Alexa488-CorM: 0.5 µM; Cy5-CorM: 4.48 pM; CorR: 0.25 µM. The Cy5-CorM monomer within the Alexa488-CorM filament remains static while Alexa488-CorM monomers are added to both filament poles. (**G**) Kymograph of CorM filament depicted in (F), confirming the immobile nature of the Cy5-CorM monomer and the bipolar extension of the CorM filament. (**H**) Start (triangle) and end (cross) positions of the two ends of the filament ends (light and dark green) shown in (F) as well as the position of the Cy5-CorM monomer (orange) during the time lapse. (**I**) Growth and shrinkage properties of CorM filaments from *in vitro* time-lapse TIRF microscopy. Scatter plots show the mean ± SD filament velocities of Alexa488-CorM during bipolar growth (76.63 ± 19.23 nm/s; n=13), unipolar growth (29.69 ± 8.28 nm/s; n=10), and shrinkage (-1313.42 ± 671.54 nm/s; n=5), as determined by single-filament tracking from TIRF microscopy. Bipolar growth and shrinkage were measured from single and dual color time-lapse imaging series using Alexa488-CorM or Alexa488-CorM with low levels of Cy5-CorM, while unipolar growth was extracted from dual color time-lapse imaging series only.

When Alexa488-labeled CorM was added to SLBs in the presence of CorR, it initially assembled into individual filaments that tended to swivel on the membrane surface. Over time, these filaments formed stable and increasingly dense networks, eventually giving rise to a highly complex filamentous mesh (Fig. 5B). We saw the same filament assembly phenotype over a range of CorM concentrations (from 1-2.5 µM). Fluorescence recovery after photobleaching (FRAP) experiments confirmed the dynamic nature of CorM filaments *in vitro* (Fig. S11A; Supplementary Movie 6), and filament turnover rates increased with rising concentrations of CorR (half recovery times (T_1/2_) for different CorR concentrations: 0.125 µM: 10.64 ± 0.87 s; 0.25 µM: 6.97 ± 0.24 s; 0.75 µM: 4.94 ± 0.14 s) (Fig. S11B).

The ^Δ1–40^CorM mutant retained dynamic behavior, although FRAP recovery was significantly slower than for the WT protein (T_1/2_ WT: 10.64 ± 0.87 s; T_1/2_ ^Δ1–40^CorM: 24.65 ± 1.22 s) (Fig. S11A, C, D; Supplementary Movie 7). Notably, filament architecture differed substantially: while initial filament formation resembled that of WT CorM, ^Δ1–40^CorM ultimately formed a much more ordered and aligned network (Fig. S11E), again suggesting that the IDR modulates filament architecture over time.

Consistent with observations in live cells, CorM filaments displayed dynamic instability *in vitro*, undergoing repeated cycles of growth and sudden shrinkage events (Fig. 5C-E; Supplementary Movie 8). Growth was observed at both filament ends (Fig. 5F-I; Supplementary Movie 9), with a mean growth velocity of 76.63 ± 19.23 nm/s (Fig 5I), comparable to EcParM (*49*). Dual-color imaging, where individual monomers were labeled with a different fluorophore than the rest of the filament, revealed that while filaments grew at both ends, individual monomers within the filament remained static (Fig. 5F-H). Intriguingly, when we quantified bidirectional growth, we saw that one filament end grew slightly faster than the other (faster end: 35.85 ± 5.53 nm/s; slower end: 23.53 ± 5.36 nm/s), on average resulting in a net unipolar growth rate of 29.69 ± 8.28 nm/s (Fig. 5I). As the two ends are structurally identical, we assume that the observed kinetic difference is likely due to only one filament end being engaged by CorR at a given time. End-binding of CorR would also be consistent with previous reports on the ParMR system (*37*) and would also align with the observed swiveling motion of CorM filaments on membrane surfaces. Due to its small size and the essential roles of its N-and C-termini, we were unable to generate a fluorescently labeled version of CorR to verify this assumption. Shrinkage occurred at a much higher rate, with mean disassembly velocities of-1313.42 ± 671.54 nm/s (Fig. 5I), approximately eight times faster than previously reported for EcParM (*49*).

### Control of CorMR filaments by the Min system

Plasmid-encoded ParMR systems typically function independently of host regulation and no other regulators or interactors have been described to date. In contrast, the distinct filament pattern of CorM *in vivo* (Fig. 2A) and the pronounced phenotype observed upon CorMR deletion (Fig 1E) suggest active spatial regulation of CorM polymerization and possible interactions with proteins involved in cell wall remodeling.

To identify putative binding partners for CorM, we performed an *in silico* screen using AlphaFold-Multimer (*54*, *55*) of CorM against the entire *Anabaena* proteome (Fig. 6A; Supplementary File 2). This approach uses structural predictions of pairwise protein-protein interactions and ranks them according to their interface-predicted template modeling score (ipTM score). Consistent with our experimental data, the top-ranking predictions included CorM-CorR (rank 8) and CorM-CorM (rank 12) interactions, both with ipTM scores above 0.65. Notably, MinC - a well-characterized inhibitor of FtsZ polymerization and an element of the Min system (*56*) - ranked second in the list, with an ipTM score of 0.77 (Fig. 6A, Fig. S12A). This unexpected prediction raised the possibility that, in multicellular cyanobacteria, MinC may regulate not only the Z-ring formation but also the CorMR system.

**Fig. 6.**
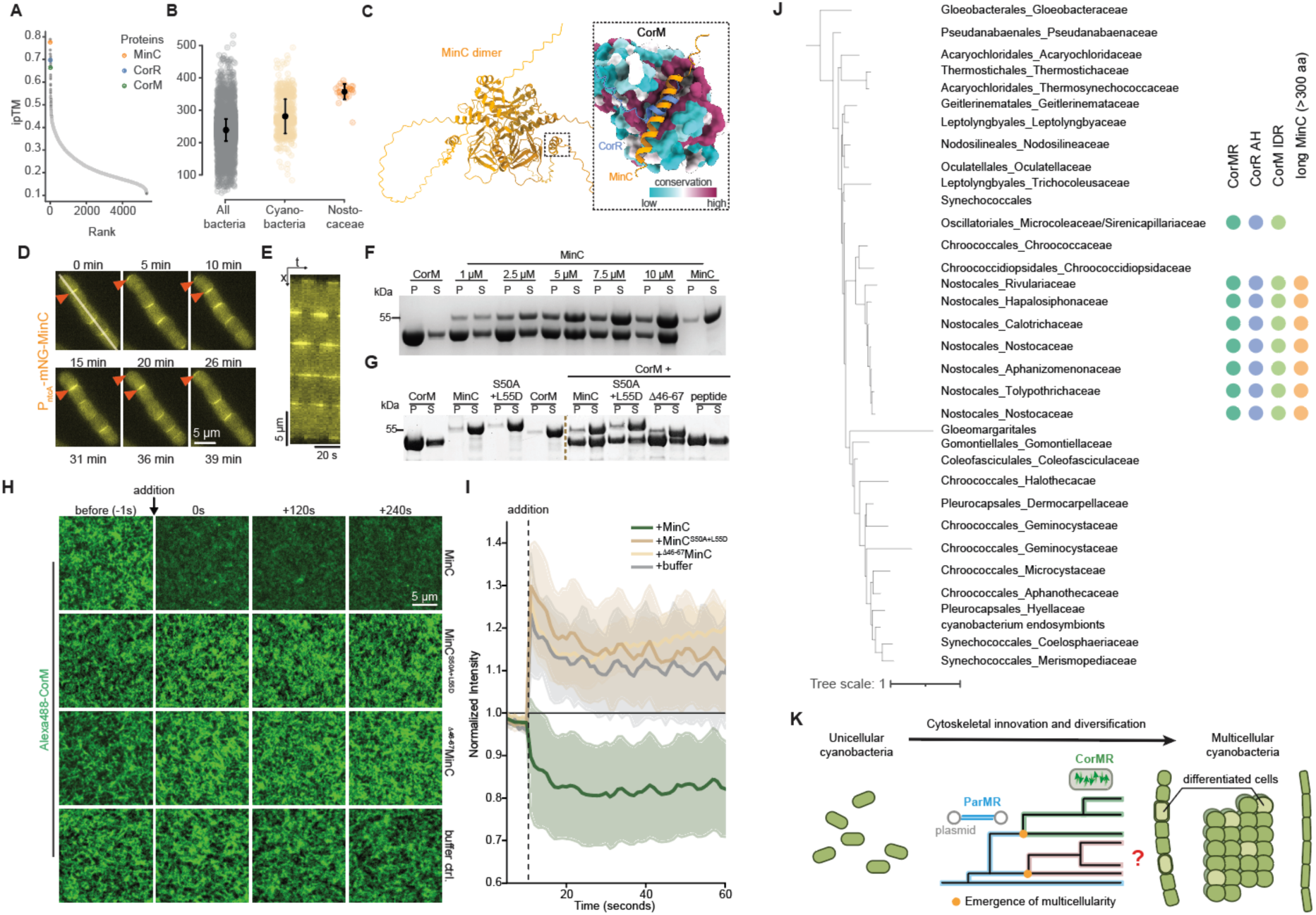
Control of CorMR filaments by the Min system. (**A**) *in silico* pulldown to identify interactions of CorM (bait) against the whole *Anabaena* proteome (5326 open reading frames included in the analysis). Accuracy of the predicted interaction is given as interface predicted template modelling (ipTM) score as described by AlphaFold2 (*77*). ipTM scores for CorM, CorR and MinC are highlighted in the plot. (**B**) Scatter plots of MinC sequence lengths (in aa) from all bacteria, cyanobacteria and *Nostocales* cyanobacteria. (**C**) (top) AlphaFold3 prediction of an *Anabaena* MinC dimer. (bottom) A zoom-in into a well-conserved pocket of CorM predicted to bind to the C-terminal α-helix of CorR (blue) and the N-terminal α-helix of MinC (orange), suggesting a competition of CorR and MinC for binding to CorM. Conservation is based on a multiple sequence alignment from all identified cyanobacterial chromosomal CorMs (green clade). Coloring by sequence conservation is based on the entropy-based measure AL2CO from ChimeraX (*78*) using the default coloring scheme (cyan: poor conservation; white: intermediate conservation; red: highly conserved; grey: no conservation). (**D**) *in vivo* time lapse microscopy of *Anabaena* cells expressing mNG-MinC from a synthetic *ntcA* promoter integrated into the neutral *thrS2* site on the *Anabaena* chromosome. Orange arrows indicate the positions of mNG-MinC fluorescence maxima in the uppermost cell over time, highlighting its pole-to-pole oscillatory behavior throughout the time-lapse analysis. Grey bar spanning the cell filament indicates region used to generate the kymograph show in (**E**). (**F**) Representative Coomassie-stained SDS-PAGE gel from a high-speed ultracentrifugation-based pelleting assay of 5 µM CorM with or without indicated MinC concentrations in the presence of 2.5 mM ATP (i.e., under polymerizing conditions). 40 % of the total pellet [P] and 11.1 % of the total supernatant [S] samples were separated on the gel. CorM runs at around 45 kDa while MinC runs at ∼55 kDa. The respective quantifications are shown in Fig. S14C. (**G**) Representative Coomassie-stained SDS-PAGE gel from a high-speed ultracentrifugation-based pelleting assay of 5 µM CorM with or without 5 µM MinC WT, MinC^S50A+L55D^, ^Δ46-67^MinC or a synthesized peptide mimicking MinC’s N-terminal α-helix in the presence of 2.5 mM ATP (i.e., under polymerizing conditions). 40 % of the total pellet [P] and 11.1 % of the total supernatant [S] samples were separated on the gel. The respective quantifications are shown in Fig. S14E. (**H**) Snapshots from TIRF microscopy time lapse series extracted from Supplementary Movies 11-14 of pre-formed CorMR filaments (0.75 µM Alexa488-CorM, 0.25 µM CorM and 0.25 µM CorR) on SLBs supplemented with 1 µM MinC WT, MinC^S50A+L55D^, ^Δ46-67^MinC or buffer. Only MinC WT can depolymerize CorMR filaments on the SLBs. We note that CorMR filaments re-assemble several minutes after addition of MinC WT, indicating that CorM depolymerization by MinC is a transient effect. (**I**) Normalized fluorescence intensity over time from the time lapse series shown in (H), with the initial value set arbitrarily to 1. A decrease in fluorescence signal is observed only upon addition of MinC WT, while both MinC mutants and the buffer control samples cause an increase in fluorescence signal, likely due to mixing after addition, which replenishes the local concentration of CorM monomers near the membrane. (**J**) Cyanobacterial phylogenetic species tree with annotations indicating the presence of a chromosomal CorMR system, an amphipathic helix (AH) in CorR homologs, an N-terminal intrinsically disordered region (IDR) in CorM homologs, and a long MinC protein (>300 aa). Monophyletic families were collapsed into single branches. Presence of the respective feature within a family is assumed if at least 50 % of the species within the clade display the feature. The complete tree is available in the Supplementary Figure 8 and Source Data, and detailed information is provided in Supplementary File 1 and 3. (**K**) A model for the evolutionary diversification and cytoskeletal innovation of an existing plasmid segregation system (ParMR), which was functionally repurposed and evolved into CorMR, thereby contributing to the emergence of complex multicellular (cyano)bacterial species.

To experimentally validate these interactions, we first used B2H assays, which confirmed that CorM interacts with MinC and revealed that CorR also interacts with MinC (Fig. S12B). However, neither CorM nor CorR interacted with MinD or MinE (Fig. S12D). Additionally, CorR was found to interact with Cdv3 (Fig. S12C), a cyanobacteria-specific component of the Min system (*14*, *18*). MinC interacted with Cdv3 and – as expected from prior findings in *E. coli,* with MinD and MinE (Fig. S12E).

Interestingly, we found that MinC proteins from multicellular cyanobacteria - particularly of *Nostocales* - are substantially longer on average than their counterparts in unicellular cyanobacteria or other bacteria (Fig. 6B, S13; Supplementary File 3). Multiple sequence alignments showed that this length increase is always due to MinC’s N-terminal extension, which is unique to *Nostocales* cyanobacteria (Fig. S13).

Structural predictions suggest that this region is intrinsically disordered and includes a short α-helix (Fig. 6C, S12A). We superimposed the predicted structures of CorM-CorR and CorM-MinC (compare Fig. S5H and S12A) and found that of MinC’s N-terminal α-helix binds to the same pocket on CorM as the C-terminal α-helix of CorR (Fig 6C). This suggests that CorR and MinC compete for CorM binding. Notably, this binding pocket of CorM is well conserved among chromosomal CorM homologs (Fig. 6C) while highly variable in plasmid-encoded ParMs (Fig. S14A), highlighting its likely relevance in regulating CorM’s function.

*In vivo*, we found that MinC oscillates from pole to pole (Fig. 6D, E; Supplementary Movie 10), as previously described for unicellular bacteria such as *E. coli* (*57*, *58*) and the rod-shaped cyanobacterium *S. elongatus* (*18*). This provides the first evidence that robust Min system oscillation can also occur in multicellular bacteria. As described in *S. elongatus*, MinC forms local maxima at the cell poles and at the sites of active constriction (Fig. S14B). This localization pattern is mutually exclusive with CorM, which is excluded from these sites, suggesting spatial antagonism.

To test for the functional relevance of the CorM and MinC interaction, we performed *in vitro* pelleting assays with CorM under polymerizing conditions while titrating MinC. We found that MinC potently depolymerized CorM in a concentration dependent manner (Fig. 6F, S14C). This effect was abolished when we mutated conserved residues in the N-terminal α-helix of MinC (S50A+L55D; Fig. S14D) or deleted the entire helix (residues 46-67) (Fig. 6G, S14E). No effect was observed upon addition of a synthetic peptide mimicking MinC’s N-terminal α-helix, suggesting that additional elements of MinC are needed to promote CorM disassembly (Fig. 6G). Depolymerization of CorM filaments was also observed *in vitro* on SLBs upon addition of WT

MinC but not with the mutant variants or buffer control (Fig. 6H, I; Supplementary Movies 11-14). Moreover, MinC addition reduced CorM ATPase activity (Fig. S14F), suggesting a sequestering mechanism of filament depolymerization as described for inhibitory proteins of actin (*59–61*), tubulin (*62*) or FtsZ (*63*) polymerization and providing a mechanistic link between MinC and filament disassembly. Finally, using the B2H system, we confirmed that the N-terminal helix is essential for MinC’s ability to interact with CorM (Fig. S14G).

Taken together, these findings reveal a novel regulatory role for the Min system in multicellular cyanobacteria. Beyond its established role in regulating Z-ring positioning, MinC modulates the CorMR system by directly competing with CorR for CorM binding, thereby spatially restricting CorM polymerization and contributing to cell shape control.

## Discussion

This study uncovers the functional repurposing of a protein system - from plasmid segregation to cell shape control. In multicellular cyanobacteria, the canonical plasmid segregating ParMR system has undergone substantial evolutionary adaptation to form the cortical CorMR cytoskeleton on the cell membrane (Fig. 6J). This adaptation seems to be specific to *Nostocales* and *Oscillatoriales* cyanobacteria, most of which retain plasmid-encoded ParMR systems in addition to the chromosomal CorMR system.

The evolutionary history of ParMR appears to be complex, shaped by multiple horizontal transfers, gene duplications, and transfer between plasmids and chromosomes. These results challenge the notion that ParMR systems solely act as plasmid segregation systems, and detailed, lineage-specific analyses will be necessary to determine whether ParMR systems have acquired distinct functions independently in other groups. In Cyanobacteria, however, the evolutionary trajectory seems more straightforward, involving two ancestral duplications and some lineage-specific chromosomal acquisition, with a single chromosomal incorporation that has been transmitted vertically. Besides CorMR, two broadly distributed plasmid-associated ParMR clades are observed: one is known to mediate plasmid copy number regulation and partitioning, while the function of the other remains unknown. Observations from both bacterial and cyanobacterial phylogenetic relationships raise the question of whether the ParMR system is more functionally plastic than previously recognized and might have been repurposed multiple times during evolution.

The transition from plasmid segregation to cell shape control is accompanied by several structural and functional innovations. Most prominently, the associated adaptor protein, CorR, has acquired an N-terminal amphipathic helix that enables membrane association while classical ParR proteins bind DNA. Our cryo-EM structure also reveals that CorM subunits form bipolar filaments, in contrast to ParM which forms polar double-stranded filaments (*25*). We also find that the canonical cell division regulator MinC has acquired a novel regulatory role. In *Nostocales*, MinC has an extended N-terminal intrinsically disordered region containing short α-helical motif, which binds a conserved pocket in CorM. Here, MinC competes with CorR binding on the one hand and also with CorM polymerization, potentially through a sequestration mechanism similar to what was reported for actin (*59–61*), tubulin (*62*) or FtsZ (*63*). In contrast to MinC’s effect on FtsZ, which does not affect FtsZ’s GTPase activity, we observed a reduction in CorM’s ATPase activity in the presence of MinC, suggesting that additional mechanisms beyond mere sequestration may be involved. As a result, MinC is able to efficiently depolymerize and remove CorM filaments from supported membranes *in vitro* as well as from the cell poles *in vivo*. This dual functionality of a Min gradient - regulating both FtsZ and CorM - is, to our knowledge, unprecedented and suggests an expanded role for the Min system in the cytoskeletal organization of multicellular cyanobacteria (Fig. 6J).

The cell shape phenotype of Δ*corMR* and Δ*mreB* (*22*) observed in *Anabaena* suggests a role for CorMR as a spatial coordinator of PG biogenesis, while the precise molecular interactions between CorMR and the PG synthesis machinery are currently under investigation and will be addressed in subsequent studies. This function is also reminiscent of plant root cortical microtubules (prcMTs), which guide cellulose synthase complexes to shape plant cells (*64–66*). Both systems operate through transversely oriented polymers and likely act in concert with cell wall synthases to control cell shape. Although CorM is an actin homolog and prcMTs are built from tubulin, the mechanistic parallels suggest an important role of dynamic instability to orient a cortical cytoskeleton perpendicular to the cell axis, a potential case of convergent evolution of their functions.

Interestingly, while Δ*corMR* and Δ*mreB*, Δ*mreC* or Δ*mreD* mutants exhibit similar morphological phenotypes - a swollen or rounded and enlarged cell (*20*, *22*) - neither system is essential for viability in *Anabaena*, even under nitrogen starvation conditions that typically fragment cell filaments. This observation contrasts with the essential role of MreBCD in rod-shaped unicellular cyanobacteria and in most other bacteria (*67–70*), highlighting the robustness and redundancy of morphogenetic systems in multicellular cyanobacteria.

CorM and MreB form antiparallel, double-stranded filaments, where individual protofilaments have opposite orientation (*71*). In contrast to MreB filaments, whose motion relies on PG synthesis (*6*, *72*), we find that CorMR filaments exhibit dynamic instability both *in vivo* and *in vitro*, similar to its ancestor ParM. Unlike canonical polymeric systems composed of actin, tubulin, ParM, or FtsZ, which are intrinsically polar and stabilized through nucleotide caps at their structurally distinct plus ends (*27*, *49*, *73*, *74*), CorM filaments are antiparallel, and thus both filament ends are structurally and kinetically identical. Accordingly, the classic nucleotide cap model for dynamic instabilities of microtubules (*75*) cannot be applied here and additional mechanistic studies are necessary to reveal how this is achieved in a bipolar, antiparallel filament.

Despite the symmetry of CorM filaments, we observed slightly different growth velocities at the two ends *in vitro* (Fig. 5I) and apparent uni-directional growth *in vivo* (Fig. 2F, G). We believe that these asymmetric growth kinetics arises from the interaction with CorR. Based on the previous observations of a clash between the C-terminal peptide of ParR with a ParM monomer within the ParM filament (*37*) and the stabilization of the ATP cap at the filament end during growth (*49*), we believe that CorR competes with the addition of another CorM monomer during polymerization. Such a competition between CorR and CorM at one of the two ends could not only explain asymmetric growth kinetics, but also the characteristic swiveling of membrane-bound CorM filaments and accelerated CorM turnover at higher CorR concentrations (Fig. S11B).

The CorMR system exemplifies evolutionary innovation, having been repurposed from a plasmid segregation module into a membrane cytoskeleton that controls cell shape (Fig. 6K). Our structural data for CorM filaments adds to the literature showing that bacterial actin homologs display both great sequence and quaternary diversity, in contrast to the highly conserved filaments observed in eukaryotes (*76*). In addition, our findings establish the chromosomal CorMR as a novel membrane-associated actin system in bacteria that assembles into bipolar, dynamically instable filaments. Our findings highlight the plasticity of bacterial cytoskeletal systems and demonstrate how pre-existing modules can be co-opted and functionally diversified to serve novel cellular roles, offering insights into the evolution of cellular and multicellular complexity.

## Acknowledgments

We thank all members of the Martin Loose lab at ISTA for helpful discussions and Marko Kojic for critical reading of the manuscript. This research was supported by the Scientific Service Units (SSU) of ISTA through resources provided by the Imaging & Optics Facility (IOF), the Scientific Computing (SciComp) and the Electron Microscopy Facility (EMF), as well as the Lab Support Facility (LSF). We are grateful for Antonia Herrero (Sevilla University) for sharing her extensive BACTH plasmid library and other plasmids as well as cyanobacterial strains. Likewise, we would like to thank Tal Dagan and Fabian Nies (both Kiel University) for sharing cyanobacterial strains and plasmids and for valuable discussions. We would further like to express our gratitude to Nicolas Sapay and Alexis Michon for providing the Amphipaseek code, which enabled us to perform our large-scale amphipathic helix screen of cyanobacterial CorR proteins. Finally, we also want to thank Jesse Hansen for advice in cryo-EM data processing.

Views and opinions expressed are those of the authors only and do not necessarily reflect those of the European Union or the European Research Council. Neither the European Union nor the granting authority can be held responsible for them.

## Funding

European Union’s Horizon 2020 research and innovation program under the Marie Skłodowska-Curie Grant Agreement No.101034413 (BLS)

European Research Council (*ERC*) of the European Unition grant ActinID, No. 101076260 (FKMS).

Swiss National Science Foundation Starting Grant (TMSGI3_226208) and the Jean-Jacques et Letitia Lopez-Loreta Foundation (both GLW).

## Author contributions

Conceptualization: BLS, ML

Methodology: BLS, MGJ, DM, RH, BZ, GLW, DMH

Investigation: BLS, MGJ, DM, RH, DMH

Visualization: BLS, MGJ, DM, RH, FKMS, ML

Funding acquisition: BLS, FKMS, ML

Project administration: BLS, ML

Supervision: ML

Writing – original and revised draft: BLS and ML, with contributions from all co-authors

## Competing interests

Authors declare that they have no competing interests.

## Data and materials availability

The electron microscopy density maps of the different CorM filament structures have been deposited in the Electron Microscopy Data Bank (EMDB) under accession codes: EMD-54060 (CorM in the presence of CorR), EMD-54061 (CorM) and EMD-54062 (Δ^1-40^CorM). The model of the CorM filament refined into EMD-54060 has been deposited in the Protein Data Bank under accession code: PDB 9RMI (Extended PDB ID pdb_00009RMI). Cryo-electron tomograms have been deposited in the EMDB under accession codes: EMD-55454 and EMD-55455.

Temporarily, all EM-associated data is accessible from: https://seafile.ist.ac.at/d/fb1870908a4c40718475/

Source Data for phylogenetic analysis will be available in a Mendeley repository under accession code 10.17632/d6hmm7bjnt.1 upon publication, but is temporarily accessible from: https://data.mendeley.com/preview/d6hmm7bjnt?a=9155e04d-88b1-4719-8ec6-0a300aa479f6

All other data are either directly enclosed to this manuscript or available from the authors upon reasonable request.

## Materials and Methods

### Bacterial strains and growth conditions

*Escherichia coli* strain DH5α was used for routine cloning procedures and chemically competent *E. coli* were transformed with plasmid DNA by the standard heat shock procedure. BL21 (DE3) (NEB, USA) or C41 (DE3) (Merck, USA) were used for heterologous recombinant protein expression. Strain BTH101 was used for bacterial adenylate cyclase two-hybrid system (BACTH) analysis. All *E. coli* strains listed in Supplementary Table 1 were grown in standard LB or TB medium containing the appropriate antibiotics at standard concentrations. *Anabaena* was a gift from Tal Dagan (Kiel University, Germany) and cells were routinely grown photoautotrophically in BG11 medium at constant illumination with an intensity of 1200 Lux (using a Osram T8 L 15W/840 LUMILUX Cool White G13 light bulb) or at RT with ambient light exposure for extended culturing. When appropriate, liquid growth medium or agar plates (containing 1.5 % agarose) were supplemented with 50 µg/ml neomycin or 5% sucrose.

### Plasmid construction

All plasmids generated in this study (see Supplementary Table 1) were constructed using an in-house Gibson assembly mix or the Q5^®^ Site-Directed Mutagenesis Kit (NEB, USA). Primers employed for plasmid construction are listed in Supplementary Table 2. In each case, plasmid sequence integrity was verified by Sanger sequencing by Microsynth Austria GmbH (Austria). In addition, we are grateful for Antonia Herrero, Fabian Nies and Tal Dagan for a wealth of plasmids, particularly for the BACTH plasmid library from Antonia Herrero.

pBAD33.1-Km: The nptII cassette was amplified from pRL25C using primers (oBS218+oBS219) and integrated into PCR-amplified pBAD33.1 (using primers oBS216+oBS217). pBS1: *corM* was amplified from *Anabaena* genomic DNA using primers oBS8+oBS9 and then integrated into HindIII+BamHI-digested pKNT25 by Gibson assembly. pBS2: *corM* was amplified from *Anabaena* genomic DNA using primers oBS10+oBS11 and then integrated into BamHI+EcoRI-digested pKT25 by Gibson assembly. pBS3: *corM* was amplified from *Anabaena* genomic DNA using primers oBS8+oBS9 and then integrated into HindIII+BamHI-digested pUT18 by Gibson assembly. pBS4: *corM* was amplified from *Anabaena* genomic DNA using primers oBS12+oBS13 and then integrated into BamHI+EcoRI-digested pUT18C by Gibson assembly. pBS9: corR was amplified from *Anabaena* genomic DNA using primers oBS18+oBS19 and then integrated into HindIII+BamHI-digested pKNT25 by Gibson assembly. pBS10: *corR* was amplified from *Anabaena* genomic DNA using primers oBS20+oBS21 and then integrated into BamHI+EcoRI-digested pKT25 by Gibson assembly. pBS11: *corR* was amplified from *Anabaena* genomic DNA using primers oBS18+oBS19 and then integrated into HindIII+BamHI-digested pUT18 by Gibson assembly. pBS12: *corR* was amplified from *Anabaena* genomic DNA using primers oBS22+oBS23 and then integrated into BamHI+EcoRI-digested pUT18C by Gibson assembly. pBS37: *corM* was was amplified from *Anabaena* genomic DNA using primers oBS61+oBS62 and ligated into PCR-amplified pTB146 (using primers oBS59+oBS60) by Gibson assembly. pBS41: *corR* was amplified from *Anabaena* genomic DNA using primers oBS69+oBS70 and integrated into PCR-amplified pTB146 (using primers oBS59+oBS60) by Gibson assembly. pBS47: *parR* from the *E. coli* R1 plasmid was amplified from a gBlock (Twist Biosciences, USA) using primers oBS81+oBS82 and ligated into PCR-amplified pTB146 (using primers oBS59+oBS60) by Gibson assembly. pBS71: *corM* with 3x 5’ glycine codons for sortagging was amplified from *Anabaena* genomic DNA using primers oBS62+oBS126 and ligated into PCR-amplified pTB146 (using primers oBS59+oBS60) by Gibson assembly. pBS75: *parM* from the *E. coli* R1 plasmid was amplified from a gBlock (Twist Biosciences, USA) using primers oBS80+oBS130 and ligated into PCR-amplified pTB146 (using primers oBS59+oBS60) by Gibson assembly. pBS80 was constructed using Gibson assembly to ligate three PCR-amplified fragments. The vector was amplified from plasmid pTB146 using primers oBS59+oBS150, *mNeonGreen* (*mNG*) was amplified from a gBlock (Twist Biosciences, USA) using primers oBS151+oBS119 and *corM* was amplified from *Anabaena* genomic DNA using primers oBS120+oBS62. pBS81 was constructed using Gibson assembly to ligate three PCR-amplified fragments. The vector was amplified from plasmid pTB146 using primers oBS59+oBS150, *mNG* was amplified from a gBlock (Twist Biosciences, USA) using primers oBS154+oBS155 and *corM* was amplified from *Anabaena* genomic DNA using primers oBS152+oBS153. pBS103 was generated by annealing of primers oBS199+oBS200 and ligating the annealed primers into AarI-digested pCpf1b using T4 ligase (NEB, USA). pBS105 was generated by annealing of primers oBS203+oBS204 and ligating the annealed primers into AarI-digested pCpf1b using T4 ligase (NEB, USA). pBS108 was constructed by Gibson assembly, ligating two PCR-amplified fragments - amplified from *Anabaena* genomic DNA using primers oBS205+oBS206 and oBS207+oBS208 - into BamHI-digested pBS105. For pBS112, a synthetically optimized P_ntcA_ was amplified from pCSL135 (*79*) using primers oBS228+oBS229 and *mNG-corM* was amplified from pBS80 using primers oBS118+oBS221 and ligated into ClaI+EcoRI-digested pRL25C by Gibson assembly. For pBS113, P_ntcA_ was amplified from pCSL135 using primers oBS229+oBS230 and *corM-mNG* was amplified from pBS81 using primers oBS241+oBS223 and ligated into ClaI+EcoRI-digested pRL25C by Gibson assembly. pBS134 was constructed using Gibson assembly to ligate three PCR-amplified fragments. The vector was amplified from plasmid pTB146 using primers oBS59+oBS150, *mNG* was amplified from a gBlock (Twist Biosciences, USA) using primers oBS151+oBS119 and *parM* from the *E. coli* R1 plasmid was amplified from a gBlock (Twist Biosciences, USA) using primers oBS282+oBS283. pBS135 was constructed using Gibson assembly to ligate three PCR-amplified fragments. The vector was amplified from plasmid pTB146 using primers oBS59+oBS150, *parM* from the *E. coli* R1 plasmid was amplified from a gBlock (Twist Biosciences, USA) using primers oBS284+oBS285 and *mNG* was amplified from a gBlock (Twist Biosciences, USA) using primers oBS154+oBS155. pBS136 was generated by annealing of primers oBS286+oBS287 and ligating the annealed primers into AarI-digested pCpf1b using T4 ligase (NEB, USA). pBS138 was constructed by Gibson assembly, ligating two PCR-amplified fragments of the *thrS2* neutral locus of *Anabaena* - amplified from *Anabaena* genomic DNA using primers oBS290+oBS291 and oBS292+oBS293 - into BamHI-digested pBS136. pBS140 was constructed by Gibson assembly-based ligation of a PCR product containing P_ntcA_*-mNG-corM*, amplified from pBS112 with primers oBS294+oBS295, into BamHI-digested pBS138. pBS142 was constructed by Gibson assembly-based ligation of a PCR product containing P_ntcA_*-corM-mNG*, amplified from pBS113 with primers oBS294+oBS296, into BamHI-digested pBS138. pBS154 was generated by annealing of primers oBS401+oBS402 and ligating the annealed primers into AarI-digested pCpf1b using T4 ligase (NEB, USA). pBS159 was generated by Gibson assembly-based ligation of two PCR products into PCR-amplified (using primers oBS143+oBS144); one amplifying *corR* from *Anabaena* genomic DNA using primers oBS145+oBS147 and the 2^nd^ amplifying *mScarlet-I* from a gBlock (Twist Biosciences) using primers oBS148+oBS149. pBS166 was generated by Gibson assembly-based ligation of three PCR products into BamHI-digested pBS138; one amplifying P_ntcA_ using primers oBS294+oBS297 and pCSL135 as a template, the 2^nd^ one amplifying *corR* using primers oBS338+oBS345 and *Anabaena* genomic DNA as a template and the 3^rd^ amplifying SCFP3 using primers oBS346+oBS347 and a gBlock (Twist Biosciences, USA) as a template. pBS184 was generated by Gibson assembly-based ligation of three PCR products into BamHI-digested pBS138, one amplifying P_ND_ using primers oBS294+oBS297 and pCSL135 as a template, the 2^nd^ one amplifying *corR* using primers oBS338+oBS376 and *Anabaena* genomic DNA as a template and the 3^rd^ amplifying *mNG* using primers oBS377+oBS296 and a gBlock (Twist Biosciences, USA) as a template. pBS190 was generated by annealing of primers oBS385+oBS386 and ligating the annealed primers into AarI-digested pCpf1b using T4 ligase (NEB, USA). pBS191 was generated by Gibson assembly-based ligation of three PCR products into BamHI-digested pBS190. The first product amplified 1000 bp upstream of *corM* using primers oBS205+oBS117 and *Anabaena* genomic DNA as a template. The 2^nd^ product was amplified sequentially by first amplifying *mNG* using primers oBS118+oBS119 and a gBock as a template and then using this product as a new template for the following PCR using primers oBS118+oBS378. The 3^rd^ PCR amplified parts of the *corM* open reading frame (ORF) using primers oBS388+oBS389 and *Anabaena* genomic DNA as a template. pBS192 was generated by site-directed mutagenesis using primers oBS144+oBS390 and pBS159 as a template. pBS196 was generated by Gibson assembly-based ligation of three PCR products into BamHI-digested pBS154, one amplifying a ∼1000 bp fragment that encompassed *corR* and parts of *corM* using primers oBS403+oBS404 and *Anabaena* genomic DNA as a template, the 2^nd^ one amplifying *mNG* using primers oBS405+oBS406 and pBS184 as a template and the 3^rd^ amplifying 1000 bp downstream of *corR* using primers oBS407+oBS208 and *Anabaena* genomic DNA as a template. pBS201: *minD* was amplified from *Anabaena* genomic DNA using primers oBS415+oBS416 and then ligated into HindIII+BamHI-digested pKNT25. pBS202: *minD* was amplified from *Anabaena* genomic DNA using primers oBS417+oBS418 and then ligated into BamHI+EcoRI-digested pKT25. pBS203: *minD* was amplified from *Anabaena* genomic DNA using primers oBS415+oBS416 and then ligated into HindIII+BamHI-digested pUT18. pBS204: *minD* was amplified from *Anabaena* genomic DNA using primers oBS419+oBS420 and then ligated into BamHI+EcoRI-digested pUT18C. pBS205: *minE* was amplified from *Anabaena* genomic DNA using primers oBS421+oBS422 and then ligated into HindIII+BamHI-digested pKNT25. pBS206: *minE* was amplified from *Anabaena* genomic DNA using primers oBS423+oBS424 and then ligated into BamHI+EcoRI-digested pKT25. pBS208: *minD* was amplified from *Anabaena* genomic DNA using primers oBS425+oBS426 and then ligated into BamHI+EcoRI-digested pUT18C. pBS213: *cdv3* was amplified from *Anabaena* genomic DNA using primers oBS433+oBS434 and then ligated into HindIII+BamHI-digested pKNT25. pBS214: *cdv3* was amplified from *Anabaena* genomic DNA using primers oBS435+oBS436 and then ligated into BamHI+EcoRI-digested pKT25. pBS229 was generated by site-directed mutagenesis using primers oBS119+oBS462 and pBS80 as a template. pBS230 was generated by site-directed mutagenesis using primers oBS150+oBS463 and pBS80 as a template. pBS231 was generated by site-directed mutagenesis using primers oBS60+oBS464 and pBS41 as a template. pBS233 generated by site-directed mutagenesis using primers oBS60+oBS475 and pBS71 as a template. pBS313 was generated by Gibson assembly-based ligation of three PCR products into BamHI-digested pBS138: 1) P_ntcA_-*corM* (corresponding to amino acid 1-209) was amplified with primers oBS294+oBS484 and pBS140 as template; 2) *mNG* was amplified with primers oBS377+oBS119 and pBS140 as template; 3) *corM* (corresponding to amino acids 210-283) was amplified with primers oBS485+oBS486 and pBS140 as template. pBS316: *minC* was amplified from *Anabaena* genomic DNA using primers oBS648+oBS649 and ligated into PCR-amplified pTB146 (using primers oBS59+oBS60) by Gibson assembly. pBS320 was generated by annealing of primers oBS630+oBS631 and ligating the annealed primers into AarI-digested pCpf1b using T4 ligase (NEB, USA). pBS321 was generated by Gibson assembly-based ligation of two PCR products into BamHI-digested pBS320, one amplifying a ∼1000 bp fragment upstream of *corR* using primers oBS653+oBS654 and *Anabaena* genomic DNA as a template, the 2^nd^ one amplifying a ∼1000 bp fragment encompassing *corR* without the first 10 aa (i.e., the amphipathic helix) using primers oBS390+oBS208 and *Anabaena* genomic DNA as a template. pBS324 was generated by Gibson assembly-based ligation of two PCR products into BamHI-digested pBS190, one amplifying a ∼1000 bp fragment upstream of *corM* using primers oBS205+oBS249 and *Anabaena* genomic DNA as a template, the 2^nd^ one amplifying a ∼1000 bp fragment encompassing *corM* without the first 40 aa (i.e., the N-terminal IDR) using primers oBS664+oBS527 and *Anabaena* genomic DNA as a template. pBS326 was generated by Gibson assembly-based ligation of three PCR products into BamHI-digested pBS138, one amplifying P_ND_ using primers oBS294+oBS229 and pCSL135 as a template, the 2^nd^ one amplifying *mNG* using primers oBS118+oBS119 and a gBlock as a template and the 3^rd^ amplifying *minC* using primers oBS508+oBS669 and *Anabaena* genomic DNA as a template. pBS338: ^Δ150-162^*corR* was amplified from *Anabaena* genomic DNA using primers oBS20+oBS700 and then ligated into BamHI+EcoRI-digested pKT25. pBS339: ^Δ150-162^*corR* was amplified from *Anabaena* genomic DNA using primers oBS22+oBS701 and then ligated into BamHI+EcoRI-digested pUT18C. pBS346 was generated by site-directed mutagenesis using primers oBS59+oBS711 and pBS41 as a template. pBS348 was generated by Gibson assembly-based ligation of two PCR products into BamHI-digested pBS320, one amplifying a ∼1000 bp fragment upstream of *corR* using primers oBS632+oBS633 and *Anabaena* genomic DNA as a template, the 2^nd^ one amplifying a ∼1000 bp fragment downstream of *corR* using primers oBS350+oBS208 and *Anabaena* genomic DNA as a template. pBS353 was generated by site-directed mutagenesis using primers oBS720+oBS721 and pBS316 as a template. pBS354 was generated by site-directed mutagenesis using primers oBS722+oBS723 and pBS316 as a template. pBS356 was generated by Gibson assembly-based ligation of two PCR products into BamHI-digested pBS154, one amplifying a ∼1000 bp fragment upstream of *corR*’s C-terminal peptide (i.e., omitting amino acid 150-162) using primers oBS403+oBS724 and *Anabaena* genomic DNA as a template, the 2^nd^ one amplifying a ∼1000 bp fragment downstream of *corR* using primers oBS725+oBS208 and *Anabaena* genomic DNA as a template. pBS386 was generated by site-directed mutagenesis using primers oBS720+oBS721 and pCSAV269 (a gift from Antonia Herrero, Sevilla University) as a template. pBS387 was generated by site-directed mutagenesis using primers oBS722+oBS723 and pCSAV269 as a template. pBS396 was generated by Gibson assembly-based ligation of two PCR products into BamHI-digested pBS103. The first product amplified 1000 bp upstream of *corM* and *mNG* using primers oBS205+oBS119 and pBS191 as a template. The 2^nd^ product amplified 882 bp downstream of codon 40 of *corM* using primers oBS651+oBS389 and pBS191. pBS415 was generated by site-directed mutagenesis using primers oBS736+oBS844 and pCpf1b as a template. pBS416 was generated by Gibson assembly-based ligation of three PCR products into EcoRI+BamHI-digested pBS415: 1) P_petE_ was amplified from *Anabaena* genomic DNA using primers oBS811+oBS845; 2) *corMR* was amplified from *Anabaena* genomic DNA using primers oBS834+oBS835; 3) the *rrnB* T2 terminator was amplified from pAM5411 using primers oBS806+oBS846. pBS417 was generated by Gibson assembly-based ligation of three PCR products into EcoRI+BamHI-digested pBS415: 1) P_petE_ was amplified from *Anabaena* genomic DNA using primers oBS805+oBS845; 2) *mNG* with a GSAGSAAGSGEFGS linker was amplified from pBS140 using primers oBS118+oBS689) *corMR*-*rrnB* T2 terminator were amplified from pBS416 using primers oBS836+oBS846.

### Strain construction

Transformation into *Anabaena* WT or mutant strains was achieved using a triparental mating, essentially as described before in (*45*), but with the following modification: instead of *E. coli* HB101, we used *E. coli* DH5α. Over-night grown *E. coli* and *Anabaena* were mixed and plated onto nitrocellulose filters (0.45 µm pore size) placed on BG11 plates supplemented with 5% LB pre-mixed powder. Cells were incubated for 6 - 8 hours at 30 °C under low light and then transferred to BG11 plates containing 50 µg/ml neomycin (BG11+Nm) and exposed to full light. Colonies routinely appeared after 10 - 12 days. These were picked and re-streaked on BG11+Nm plates to isolate single colonies and eliminate residual *E. coli* for several rounds. For heterologous, plasmid-based expression of fluorescently labeled proteins, newly emerging colonies were screened by fluorescence microscopy and propagated on BG11+Nm plates instead. For chromosomal insertions, replacements or deletions, the following strategies were applied:

Strain BS140 was generated by Cpf1-based heterologous insertion of the P_ntcA_-*mNG-corM* construct into the *thrS2* open reading frame (encoding threonine-tRNA ligase; NCBI Ref. Seq.: WP_010998854.1), a chromosomal neutral site (*20*, *48*). Double homologous recombination using ∼1000 bp flanking regions cloned into plasmid pBS140 replaced the *thrS2* segment encoding residues Q188–F228 with P_ntcA_-*mNG-corM*.

Strain BS142 was generated by Cpf1-based heterologous insertion of the P_ntcA_-*corM-mNG* construct into the *thrS2* open reading frame. Double homologous recombination using ∼1000 bp flanking regions cloned into plasmid pBS142 replaced the *thrS2* segment encoding residues Q188–F228 with P_ntcA_*-corM-mNG*.

Strain BS313 was generated by Cpf1-based heterologous insertion of the P_ntcA_-*corM-mNG*^SW^ construct into the *thrS2* open reading frame. Double homologous recombination using ∼1000 bp flanking regions cloned into plasmid pBS313 replaced the *thrS2* segment encoding residues Q188–F228 with a *mNG* sandwich fusion to *corM* under the control of P_ntcA_.

Strain BS326 was generated by Cpf1-based heterologous insertion of the P_ntcA_-*mNG-minC* construct into the *thrS2* open reading frame. Double homologous recombination using ∼1000 bp flanking regions cloned into plasmid pBS326 replaced the *thrS2* segment encoding residues Q188–F228 with P_ntcA_-*mNG-minC*. Complete segregation was verified by colony PCR using construct-specific primers.

For colony PCR, colonies were either picked with a pipette tip, briefly spotted onto a fresh BG11+Nm plate, and then resuspended directly in the PCR reaction mixture, or resuspended in 10 µl sterile PBS, of which 1 µl was used for each 10 µl PCR reaction.

For the Δ*corMR* (strain BS108), plasmid pBS108 was conjugated into *Anabaena* WT. Arising colonies were re-streaked twice onto fresh BG11+Nm plates. Deletion of *corM-corR* was confirmed by colony PCR using site-and gene-specific primers, followed by Sanger sequencing of the encompassing regions. The same procedure was applied for the verification of ^Δ1-10^*corR* (strain BS321), ^Δ1-40^*corM* (strain BS324), *ΔcorR* (strain BS348) and *corR*^Δ150-162^ (strain BS356) mutants, using plasmids pBS321, pBS324, pBS348 or pBS356, respectively for conjugation.

Strain BS191, expressing *mNG*-CorM as the sole copy of CorM, was generated by conjugating plasmid pBS191 into *Anabaena* WT. Resulting colonies were picked, re-streaked on BG11+Nm plates, screened for *mNG*-CorM fluorescence by microscopy, and verified by Sanger sequencing of a PCR-amplified fragment encompassing the P_corMR_-*mNG-corM-corR* region from a single colony. Strain BS396 was obtained in the same manner as BS191, using plasmid pBS396 instead of pBS191. Sanger sequencing confirmed the presence of P_corMR_-*mNG*-^Δ1–40^*corM-corR*.

In all cases, curing of pCpf1b was performed by sucrose counter-selection. For this, strains were grown in BG11 medium, sonicated in a VWR Ultrasonic Cleaner USC-TH for 10 min at maximum intensity, and then plated in 1:10 serial dilutions on BG11 plates supplemented with 5% sucrose. Arising colonies were screened by streaking on BG11 and BG11+Nm plates; those that failed to grow on BG11+Nm (indicating loss of pCpf1) were selected for further experiments.

### BACTH analysis

For protein-protein interaction analysis, the BACTH system (Euromedex, France) was employed. Genes were cloned into the expression vectors pKNT25, pKT25, pUT18, and pUT18C using GIBSON assembly, resulting in N-or C-terminal translational fusions to the T25 or T18 subunit of the *E. coli* adenylate cyclase. *E. coli* BTH101 cells were co-transformed with about 5 ng of the respective plasmid combinations and then plated on LB agar containing 200 µg/ml X-gal, 0.5 mM IPTG, 100 µg/ml ampicillin (Amp), and 50 µg/ml Kanamycin (Km). As a positive control, plasmids pKT25-zip and pUT18C-zip and as a negative control empty plasmid pUT18 and a pKNT25 plasmid containing a test gene were also co-transformed into BTH101 cells. The plates were incubated at RT for 2-3 days. To then measure beta-galactosidase as a surrogate measure to quantify protein-protein interaction strengths, a modified protocol from Karimova et al. (*80*) was employed. For each combination, three independent colonies were picked and grown in 300 µl LB supplemented with 0.5 mM IPTG, Amp, and Km in a 2 ml 96-well deep well plate for 2-3 days at RT and 1000 rpm in an orbital microplate shaker.

Afterwards, 50 µl of cells were diluted with 50 µl 2x M63 medium and OD_595_ values were recorded in a 96-well plate. Cells were then lysed by adding 10 µl of PopCulture^®^ Reagent (Merck, USA) supplemented with 10µl/ml Lysonase™ Bioprocessing Reagent (Merck, USA). Lysis was conducted for at least 30 min at RT and rigorous shaking with 650 rpm in the 96-well plate on an orbital microplate shaker (We find that even hour-long lysis did not decrease experimental readout). Subsequently, 20 µl of lysate was added to 105 µl of PM2 assay medium supplemented with 100 mM β-mercaptoethanol in a new 96-well plate and incubated at RT with mild shaking for 20 min before the reaction was stopped by adding 50 µl of 1 M Na_2_CO_3_.

Finally, OD_405_ values were recorded, and beta-galactosidase activity was calculated as described in the BACTH System Kit (Euromedex).

### Microscopy

Experiments were performed using a Nikon Eclipse Ti2 microscope equipped with iLas2 (GATACA) 360° / Ring / Azimuthal TIRF module, a TI2-N-ND-P Perfect Focus System and either a CFI Apochromat TIRF 60x oil immersion objective (NA 1.49, WD 0.13 mm) or a CFI Plan Apo Lambda D 100x oil immersion objective (NA 1.45, WD 0.13 mm).

Fluorophores and fluorescent proteins were excited using the Omicron LightHUB Ultra laser system for TIRF imaging and Fluorescence Recovery After Photobleaching (FRAP), equipped with 405nm, 488nm, 561nm and 640nm lasers. Emitted fluorescence signals were filtered using a Nikon 525/50 ET bandpass and a 635 nm long pass filter for dual-color imaging. For the dual-color experiments, a Cairn TwinCam camera splitter equipped with a TwinCam Cube dichroic mirror for green/red,far-red was used in addition.

For epifluorescence microscopy, LED-FITC-A Filter Cube EX 474/27, EM 525/45 and LED-Cy5-A Filter Cube EX 628/40, BA 692/40 filter cubes were used, and Direct Illumination Acquisition (DIA) images were obtained with a CoolLED pE300 White MB light source. The microscope was operated using the NIS elements AR (v.5.42.04) software and images were recorded using one or two Andor iXon Life 888 Back Illuminated EMCCD cameras.

High-resolution spinning disk confocal microscopy was conducted using a Nikon Ti2 CSU W1-01 SoRa+NIR microscope, equipped with a Nipkow disk-based Super-resolution via optical Re-assignment (SoRa) module and a Nikon CFI Plan Apochromat 100× silicone immersion objective (NA 1.35, WD 0.3 mm). Excitation was provided by an Omicron LightHUB UltraN laser system with 405 nm, 488 nm, and 561 nm lasers, guided through a confocal quad-band dichroic mirror (405/488/561/640 nm). Emission signals were collected through a quad-band emission filter (440/521/607/700 nm) and imaged using a Teledyne Photometrics BSI scientific CMOS camera.

### E. coli live cell imaging

Plasmids encoding CorM and CorR variants under control of P_T7_ (CorM) or P_BAD_ (CorR) were transformed into *E. coli* C41 (DE3) and plated on LB agar with ampicillin (100 µg/ml), kanamycin (50 µg/ml), or both. Single colonies were inoculated into selective LB and grown overnight at 37 °C, 220 rpm, under inducing (0.25 mM IPTG, 0.02% L-arabinose, or both) or non-inducing conditions. Overnight-induced cultures were diluted 1:100, immobilized on BG11 agar pads (2% agarose), and imaged. Uninduced cultures were diluted 1:100, induced for 3 h at 37 °C and 220 rpm, and imaged as above.

### DAPI staining

For DAPI staining, cells grown on BG11 agar plates were re-suspended in PBS supplemented with 10 µg/ml DAPI (final), incubated for 1-2 min and then spotted and immobilized on BG11 2% agar pads prior to imaging.

### Time lapse microscopy of mNG-CorM in Anabaena cells

For time-resolved tracking of mNG-CorM filaments in *Anabaena* cells of strain BS191, cells were grown on BG11 plates, re-suspended in sterile BG11 and spotted on BG11 pads containing 1.5 % low melting agarose (TopVision Low Melting Point Agarose, Thermo Fisher Scientific) supplemented with 10 mM NaHCO_3_ in a Grene Frame (65 µl; Thermo Fisher Scientific) to avoid evaporation. Cells were visualized in a Nikon Eclipse Ti2 microscope equipped with iLas2 system and mNG-CorM filaments were recorded in epifluorescence mode (50 ms exposure, 10 % blue LED light intensity). Cells were grown at RT and illuminated using a custom-made plant light equipped with three LEDs emitting blue (450 nm), red (655 nm), and far-red (725 nm) light. The light intensities at the sample position were 285.20 mW/m² (blue), 41.25 mW/m² (red), and 38.65 mW/m² (far-red). Images were acquired every 15 minutes with the perfect focus system enabled to maintain the correct focal plane.

### Protein purification

#### Purification of CorM

CorM with three N-terminal Gly (GGG-CorM) to make it amenable for sortagging was purified as a N-terminal His_6_-SUMO fusion protein expressed from pBS71 in *E. coli* C41 (DE3) cells.

For this, cells were grown at 37 °C, 220 rpm in Terrific Broth (TB) supplemented with Amp and expression was induced at an OD_600_ 0.5-0.7 with 0.5 mM isopropyl-β-thiogalactopyranoside (IPTG) for 4 h at 37 °C. Cells were harvested by centrifugation (8,000 x *g,* 20 min at 4 °C) and cell pellets were scooped out and stored in 50 ml reaction tubes at-70 °C. Cell pellets were resuspended in HEPES lysis buffer (HLB; 50 mM HEPES, 150 mM KCl, 5 mM MgCl_2_, 1 mM DTT, 25 mM imidazole and 10 % glycerol) supplemented with 1x EDTA-free protease inhibitor cocktail (Roche, Switzerland). Cells were lysed by sonication using Q700 Sonicator equipped with a 1/2 in horn tip (amplitude 40, 1 sec on + 1 sec off with 10 min on time) and subsequently incubated with 10 µl DNase I (2500 U/ml; Thermo Fisher Scientific, USA) per every 1 L of initial bacterial culture volume for 15 min. The lysate was then centrifuged for 30 min at 60,000 x *g* at 4 °C and the supernatant transferred to HisPur™ Ni-NTA Resin (1 ml per 1 L of initial bacterial culture volume; Thermo Fisher Scientific, USA) pre-equilibrated with HLB in incubated with mild agitation for 1-2 h at 4 °C. The mixture was loaded onto a gravity flow column, the flow through was discarded and the resin was washed with 3x 20 ml HLB. Proteins were eluted with HEB (HLB with 250 mM imidazole) in 1 ml steps. Eluates were briefly tested for protein content using a quick and dirty Bradford (Bio-Rad Protein Assay Dye Reagent; BioRad, USA) assay by adding 2 µl of each elution fraction to 200 µl Bradford assay solution. Protein fractions that resulted in a discernible blue color shift were pooled and dialyzed over night at 4 °C against HLB in Pur-A-Lyzer™ Maxi 6000 dialysis tubes (Sigma Aldrich, USA) supplemented with His-tagged SUMO protease (Ulp1) (ratio of 1:100). The next day, reverse affinity was performed to remove His-SUMO and His-Ulp1. For this, the solution was subjected to Ni-NTA (half the volume as used for the initial purification) pre-equilibrated in HLB, incubated for 1 h at 4 °C and mild rotation. The resin was applied to a gravity flow column and the flow through was collected followed by 3x washing with 1x column volume (CV) of HLB. The flow through and the wash fractions were pooled, concentrated using a Vivaspin 20 10KMWCO centrifugal concentrator (Sartorius, Germany) and subjected to size-exclusion chromatography (SEC) on an Äkta pure system using a HiLoad 16/600 Superdex 75 PG column (Cytiva, USA) equilibrated with HEPES storage buffer (HSB, HLB without imidazole). Protein-containing fractions were pooled, concentrated using another Vivaspin 20 10KMWCO centrifugal concentrator and then finally flash-frozen in liquid nitrogen and then stored at-70 °C.

#### Purification of Δ^1-40^CorM

^Δ1-40^CorM with three N-terminal Gly (GGG-^Δ1-40^CorM) to make it amenable for sortagging was purified as a N-terminal His_6_-SUMO fusion protein expressed from pBS233 in *E. coli* C41 (DE3) cells for 3 h at 37 °C with 0.5 mM IPTG. Cell pellets were re-suspended in HLB and lysed as described for CorM. Cell-free lysates were then subjected to Ni-NTA purification using the Äkta pure system equipped with a HisTrap^TM^ HP 5 ml column and proteins were eluted in HEB. Pooled fractions were then subjected to Ulp1 cleavage and dialyzed overnight at 4 °C against HLB. After subsequent reverse affinity, the flow through and two wash fractions were pooled, concentrated using a Vivaspin 20 10KMWCO centrifugal concentrator and subjected to another round of reverse affinity due to remaining contamination with His6-SUMO. The flow through and two wash fractions were then pooled, and protein purity was confirmed by SDS polyacrylamide gel electrophoresis (SDS-PAGE) followed by flash-freezing in liquid nitrogen and storage at-70 °C.

#### Purification of EcParM from *E. coli* R1 plasmid

ParM from *E. coli* R1 plasmid was expressed from pBS75 with three N-terminal Gly (GGG-ParM) in C41 (DE3) for 3 h at 37 °C with 0.5 mM IPTG. Lysis and subsequent Ni-NTA-based purification were done as described for CorM. After the reverse affinity, ParM was concentrated using a Vivaspin 20 10KMWCO centrifugal concentrator and subjected to buffer exchange to HSB using a PD-10 desalting column. ParM was again concentrated using a Vivaspin 20 10KMWCO centrifugal concentrator and flash-frozen in liquid nitrogen and then stored at-70°C.

#### Purification of CorR

CorR was expressed as a N-terminal His_6_-SUMO fusion protein from pBS41 in BL21 (DE3) for 3 h at 37 °C with 0.5 mM IPTG or in C41 (DE3) for 18 h at 18 °C with 0.5 mM IPTG. Lysis and subsequent Ni-NTA-based purification were done as described for GGG-CorM. After the reverse affinity, flow through and wash fractions were pooled, concentrated using a Vivaspin 20 10KMWCO centrifugal concentrator and subjected to anion exchange chromatography (AIEX) on Äkta pure system using HiTrap 1ml Q HP anion exchange chromatography column (Cytiva, USA) equilibrated with no salt buffer (50 mM HEPES pH 7.4, 5 mM MgCl_2_, 10 % glycerol). For this, CorR was diluted 1/5 in no salt buffer and subjected to a linear gradient of no salt and high salt buffer (50 mM HEPES pH 7.4, 1 M KCl, 5 mM MgCl_2_, 10 % glycerol). CorR elutes as numerous, not well-defined peaks, likely representing different oligomeric states of CorR. CorR-containing fractions were pooled, concentrated using a Vivaspin 20 5KMWCO centrifugal concentrator, flash-frozen in liquid nitrogen and stored at-70 °C.

#### Purification of ^Δ1-10^CorR

^Δ1-10^CorR was expressed as a N-terminal His_6_-SUMO fusion protein from pBS231 in BL21 (DE3) for 3 h at 37 °C with 0.5 mM IPTG. Lysis and subsequent purification steps were done as described for CorR.

#### Purification of ^Δ150-162^CorR

^Δ150-162^CorR was expressed as a N-terminal His_6_-SUMO fusion protein from pBS346 in BL21 (DE3) for 16 h at 18 °C with 0.5 mM IPTG. Lysis and subsequent purification steps were done as described for CorR.

#### Purification of ParR

ParR from *E. coli* R1 plasmid was expressed as a N-terminal His_6_-SUMO fusion protein from pBS47 in BL21 (DE3) for 3 h at 37 °C with 0.5 mM IPTG. After overnight cleavage with Ulp1, ParR was subjected to SEC using HiLoad 16/600 Superdex 75 PG equilibrated with HSB. Protein containing fractions were pooled, supplemented with 10 mM imidazole and subjected to reverse affinity. The flow through and wash fractions were pooled, concentrated using a Vivaspin 20 5KMWCO centrifugal concentrator, flash frozen in liquid nitrogen and stored at-70°C.

#### Purification of MinC

MinC with three N-terminal Gly (GGG-MinC) to make it amenable for sortagging was purified as an N-terminal His_6_-SUMO fusion protein expressed from pBS316 in *E. coli* BL21 (DE3) or C41 (DE3) cells for 3 h at 37 °C with 0.5 mM IPTG. Lysis and subsequent Ni-NTA-based purification were done as described for CorM. After the reverse affinity, MinC was concentrated using a Vivaspin 20 10KMWCO centrifugal concentrator and was then subjected to SEC using HiLoad 16/600 Superdex 75 PG column equilibrated with HSB. One major peak was detected and pooled and proteins subjected to AIEX as described for CorR with a gradient from no salt to high salt buffer. A single peak was identified, pooled, flash-frozen in liquid nitrogen and stored at-70 °C.

#### Purification of MinC^S50A+L55D^

MinC^S50A+L55D^ with three N-terminal Gly (GGG-MinC^S50A+L55D^) to make it amenable for sortagging was purified as a N-terminal His_6_-SUMO fusion protein expressed from pBS353 in *E. coli* BL21 (DE3) cells for 4 h at 37 °C with 0.5 mM IPTG. Lysis and subsequent Ni-NTA-based purification were done as described for CorM. After the reverse affinity, MinC^S50A+L55D^ was concentrated using a Vivaspin 20 10KMWCO centrifugal concentrator and was then subjected to AIEX as described for CorR. Due to remaining contamination with His6-SUMO, another reverse affinity step was included, the flow through and wash steps were collected, pooled and then dialyzed overnight at 4 °C against HSB in Pur-A-Lyzer™ Maxi 6000 dialysis tubes. Afterwards, MinC^S50A+L55D^ was concentrated using a Vivaspin 20 10KMWCO centrifugal concentrator, flash-frozen in liquid nitrogen and stored at-70 °C.

#### Purification of ^Δ46-67^MinC

^Δ46-67^MinC with three N-terminal Gly (GGG-^Δ46-67^MinC) to make it amenable for sortagging was purified as a N-terminal His_6_-SUMO fusion protein expressed from pBS354 in *E. coli* BL21 (DE3) cells for 4 h at 37 °C with 0.5 mM IPTG. ^Δ46-67^MinC was purified as described for _MinCS50A+L55D._

#### Peptide labelling

For labelling of peptide for subsequent sortagging reactions, we obtained dried CLPETGG peptides from Biomatik (USA). 1.5 mg of the peptide was dissolved in 50 mM HEPES pH 7.4 to a final concentration of 100 mM. Alexa Fluor™ 488 (Alexa488; Thermo Fisher Scientific) C5 Maleimide and Sulfo-Cyanine5 maleimide (Lumiprobe, Germany) were dissolved in DMSO to obtain a 150 mM solution. For extended storage, dissolved dyes were stored at-70 °C. A roughly 2x molar excess of TCEP was added to the reconstituted peptide (i.e., 10 µl of peptide and 5 µl of 0.4 M TCEP) to obtain a 66 mM peptide solution. Afterwards, a roughly 3x molar excess of dye was added to the peptide+TCEP mix (i.e., 20 µl of dye was added leading to a peptide concentration of roughly 28 mM and a dye concentration of roughly 86 mM). The reaction was then incubated over night at RT protected from light. The next day, 2 µl of 2-mercaptoethanol was added to quench unreacted maleimide dye. The 25 mM peptide solution was then flash-frozen in liquid nitrogen and stored at-70 °C.

#### Protein labelling - Sortagging

For Sortase-based N-terminal fluorescence labelling, we routinely prepare 1 or 2 ml reactions of 50 µM GGG-CorM, 0.5 mM labelled peptides (Alexa488-LPETGG or Cy5-LPETGG) and 10 µM Sortase 7M in HSB, which are then incubated overnight at 4 °C in the dark with gentle rotation. The next day, 3 µl of 2-mercaptoethanol was added to the reaction to quench all unreacted dye and incubated for 15 min at 4 °C. The reactions were supplemented with 25 mM imidazole (final) and 300 µl pre-equilibrated Ni-NTA slurry was added. After incubation for 60 min at 4 °C with mild rotation, the solution was applied to a gravity flow column and the flow through was collected. The Ni-NTA was washed 2x with 1x column volume HDB and pooled with the flow through. To remove unbound labelled peptide and to exchange the buffer to HSB, Cy5-CorM was then subjected to a PD-10 desalting column (Cytiva, USA) and eluted protein was concentrated using a Vivaspin 2 10KMWCO centrifugal concentrator (Sartorius, Germany). For Alexa488-CorM, unbound labelled peptide and imidazole traces were removed by size-exclusion chromatography on a Superdex 75 Increase 10/300 GL (Cytiva, USA) using HSB. Protein-containing peaks were then again concentrated using a Vivaspin 2 10KMWCO centrifugal concentrator and in both cases protein concentration was measured using NanoDrop One and labelling efficiency was calculated based on the ratio of the concentration of fluorescence dye (equals labelled protein) to the protein concentration. In both cases, labelling efficiency was about 30 %. Finally, labelled proteins were flash-frozen in liquid nitrogen and stored at-70 °C.

#### Antibody synthesis

Polyclonal rabbit antibodies raised against purified CorM protein in PBS (137 mM NaCl, 1.8 mM KH₂PO₄, 10 mM Na₂HPO₄, and 2.7 mM KCl; pH 7.4) were obtained from GenScript (Rijswijk, Netherlands)

#### Western blot analysis

To verify the antibody specificity, western blot analysis of *Anabaena* WT and *ΔcorMR* cell lysates was performed. Liquid cultures of *Anabaena* WT or *ΔcorMR* were grown for one or two weeks in BG11 and 1 ml culture was pelleted by centrifugation (5 min, 21,100 x *g*, RT) and re-suspended in 400 µl PopCulture (Merck, USA) supplemented with 1x cOmplete™, EDTA-free Protease Inhibitor Cocktail (MilliporeSigma, MA, USA). Cells were transferred to MN Bead Tubes Typ B beads (MACHEREY-NAGEL, Germany) and lysed by vortexing for 3x 1 min at maximum speed, chilling the cells on ice in between. 15 µl of cell lysates were then mixed with 3 µl of 6X reducing Laemmli SDS sample buffer (Thermo Fisher Scientific, USA). Samples were boiled for 10 min at 95 °C and then separated by SDS polyacrylamide gel electrophoresis using SurePAGE™ gradient gels (Bis-Tris, 4-20 %; GenScript, Netherlands). Upon band separation, proteins were transferred to a 0.2 µm nitrocellulose membrane (Trans-Blot Turbo Mini 0.2 µm Nitrocellulose Transfer Packs; BioRad, USA) using a Trans-Blot Turbo Transfer System (BioRad, USA) and then blocked in blocking buffer (TBST: 50 mM Tris-HCl, 150 mM NaCl, pH 7.4, 0.1 % Tween-20 supplemented with 5% non-fat dry milk) for 30 min at RT. Afterwards, the blot was incubated with rabbit anti-CorM primary antibody (GenScript, Netherlands, 1µg/ml diluted in blocking buffer) for 1 hour at RT. The blot was washed 4x with TBST and then incubated with goat anti-Rabbit IgG (H&L HRP conjugated; Agrisera, Sweden) secondary antibody (1:10,000 diluted in blocking buffer) for 1 hour at RT followed by 4x washing with TBST and visualization of protein bands using Clarity™ Western ECL Substrate (BioRad, USA) in a ChemiDoc MP imaging system (BioRad, USA).

#### Immunofluorescence staining

For immunofluorescence analysis of CorM localization, *Anabaena* WT cells were grown in BG11 liquid culture for about 2 weeks and 2x 1 ml culture was pelleted by centrifugation (2 min, 2000 x *g*, RT) and once washed with 1 ml TBS (50 mM Tris-HCl, 150 mM NaCl, pH 7.4) by centrifugation (2 min, 2000 x *g*, RT). The supernatant was removed, and cells were re-suspended in 1 ml ice-cold 100 % MeOH and incubated at-20 °C for 20 min. Afterwards, 40 µl of cells in MeOH were spotted into *in situ* Gene Frame® II hybridization frames (65 µl variants; Thermo Fisher Scientific, USA) attached onto Polysine microscope slides (Menzel, USA) and air-dried. Cells were washed for 5 min by adding 200 µl TBS to into the Gene Frame, followed by aspiration of the TBS solution and incubation for 30 min at 30 °C with 200 µl of 25 µg/ml lysozyme solution (25 µg/ml lysozyme in 50 mM Tris-HCl (pH 8.0), 25 mM NaCl, 2 mM EDTA). We note that using higher lysozyme concentrations (i.e., 1 mg/ml) had no discernible effect. The lysozyme solution was removed, and frames were washed 3 x with 200 µl TBS for 5 min at RT. After aspiration of the TBS, cells were blocked with 200 µl blocking buffer (TBS supplemented with 0.05% Tween-20 and 3 % BSA) for 30 min at RT. After removing the blocking solution, the cells were incubated overnight at 4 °C with 200 µl of either blocking buffer (no primary antibody control control) or a 1:100 dilution of the anti-CorM primary antibody (GenScript, Netherlands) in blocking buffer, sealed to prevent evaporation. The next day, the primary antibody solution was removed, and cells were washed 3x with 200 µl TBST (TBS with 0.05 % Tween-20) for 5 min at RT. The TBST solution was removed, and cells were incubated in 200 µl goat anti-Rabbit IgG (H+L) Highly Cross-Adsorbed secondary antibody Alexa Fluor Plus 488 (1.5 µg/ml; Thermo Fisher Scientific, USA) in blocking buffer for 3 hours at RT in the dark. Secondary antibody solution was removed, and cells were washed 3x with 200 µl TBST for 5 min at RT. Finally, the TBST solution was removed, and the cells were allowed to air-dry. The Gene Frame was then detached from the Polysine slides, and the cells were mounted in a self-prepared mounting medium (20 mM Tris-HCl pH 8.0, 0.5% N-propyl gallate, 90% glycerol) and sealed with VALAP (a 1:1:1 mixture of Vaseline, lanolin, and paraffin) prior to imaging.

#### Piranha treatment of coverslips for TIRF microscopy

TIRF-suitable glass coverslips (18X18 mm NR 1.5 high precision cover glasses; Marienfield, Germany) in a custom-made Teflon holder were subjected to piranha solution (mixture of 50 ml 30 % H_2_O_2_ and 150 ml H_2_SO_4_) for 2 h at RT until the solution cooled down to RT. Cover slips were then washed 10x with 500 ml MiliQ H_2_O, sonicated in a sonicator bath (VWR Ultrasonic Cleaner USC - TH) for 20 min and washed again 4x with 500 ml MiliQ H_2_O and finally stored at RT in 500 ml MiliQ H_2_O for no longer than 2 weeks. For TIRF microscopy, coverslips were dried by compressed air and reaction chambers made of 0.5 ml reaction tubes (cut in half to create a chamber without the conical end) were glued onto the coverslips with a UV-reactive glue (Norland Optical Adhesive 63) for 2 min under UV light exposure.

#### Preparation of multilaminar vesicles and small unilaminar vesicles

For pelleting assays and TIRF microscopy experiments we used DOPC (1,2-dioleoyl-sn-glycero-3-phosphocholine) and DOPG (1,2-dioleoyl-sn-glycero-3-phospho-(1′-rac-glycerol)) *E. coli* polar lipids (Avanti, USA) dissolved in chloroform (stock concentration 2.5 mg) at a ratio of 67:33 mol %. Lipids were mixed in a glass vial and then dried in a stream of nitrogen gas.

Residual chloroform was removed from the lipids by vacuum desiccation for at least 3 h or overnight. Afterwards, lipids were resuspended by vortexing in HEPES reaction buffer (HRB; 50 mM HEPES [pH 7.4], 150 mM KCl, 5 mM MgCl_2_) to obtain a 5 mM large multilaminar vesicles (MLVs) solution. To prepare small MLVs, this solution was then incubated at 37 °C for 30 min and then subjected to 7x freeze-thaw cycles using a dry ice-isopropanol bath and hot tap water. Small MLVs were then subjected to centrifugation at 21,000 x *g*, 4 °C, 30 min and the supernatant was transferred to a new reaction tube, which was then overlayed with argon gas and stored at 4 °C until further use. For preparation of small unilaminar vesicles (SUVs), small MLVs were subjected to tip-sonication (Q700 Sonicator equipped with a 1.6 mm tip) on ice with the following settings: amplitude 1; 1 s on time, 4 s off time; total 5 min on time. Afterwards, SUVs were centrifuged for 20 min at 21,000 x *g*, 4°C and the supernatant was transferred to a new tube and overlayed with argon gas.

#### Supported lipid bilayer preparation

For the preparation of supported lipid bilayers (SLBs), 100 µl reactions were prepared in reaction chambers containing 89 µl HRB, 0.5 mM SUVs and vesicle rupture was induced by adding 5 mM CaCl_2_ to the SUVs. For fusion of the ruptured vesicles with the piranha-treated glass coverslips, chambers were incubated for 20 min at 37 °C followed by extensive washing with 6x 200 µl HRB to remove unfused vesicles. Chambers were again washed 3x with 200 µl HRB right before each individual experiment. SLBs were prepared fresh for each experiment day.

#### Total Internal Reflections Fluorescence microscopy

For Total Internal Reflections Fluorescence (TIRF) microscopy, 0.75 µM unlabeled CorM or ^Δ1-40^CorM, 0.25 µM Alexa488-CorM or Alexa488-^Δ1-40^CorM, 0.25 µM CorR and an oxygen scavenger mix composed of 1 µM glucose oxidase from *Aspergillus niger* (Merck, USA), 332 nM catalase from bovine liver (Merck, USA), 11 mM D-glucose, 1 mM DTT and 1 mM Trolox was used. The equal amount of buffer in the reaction chambers containing the SLBs was removed and only then all components were added to the chambers. CorM polymerization on SLBs was induced by the addition of 2.5 mM ATP and CorM polymers were observed by TIRF imaging. To test for the effect of different CorR concentrations, the same kind of reactions were prepared but with varying indicated CorR concentrations.

For dual color imaging to study the localization of individual CorM monomers within a CorM protein filament, a mixture of either 1 µM Alexa488-CorM + 265 pM Cy5-CorM + 0.25 µM CorR or 0.25 µM Alexa488-CorM + 265 pM Cy5-CorM + 0.75 µM CorM + 0.25 µM CorR was added to the reaction chambers together with the oxygen scavenger mix. Time-lapse movies were recorded between 2-5 min after addition of 2.5 mM ATP to the reaction.

To test for the effect of MinC, MinC^S50A+L55D^ and ^Δ46-67^MinC addition on CorM filaments *in vitro*, CorM filaments were polymerized on SLBs in the presence of CorR and 2.5 mM ATP for about 5 min upon which equal molar amounts of MinC or MinC mutants (i.e., 1 µM) was added to the chambers during time lapse imaging. Movies were recorded for 3 min each.

#### Fluorescence recovery after photobleaching experiments

For FRAP analysis, 3 - 6 pre-bleach frames were acquired, followed by photobleaching of a defined rectangular region of interest (ROI) using 80 % 488 nm laser power and a pixel dwell time of 100 µs/pixel. Fluorescence recovery within the ROI was subsequently recorded at a rate of 1 frame per second.

#### Pelleting assay

Two types of pelleting assays were performed in this study. One did not involve MLVs and relied on ultracentrifugation-based pelleting of protein polymers while the other employed the intrinsic binding capacity of proteins to MLVs. Pelleting assays with MLVs were either performed in a suspension with or without 0.93 mM small MLVs and proteins at indicated concentration were incubated for 30 min at 30 °C in HRB supplemented with 2.5 mM ATP. Unless otherwise stated, proteins were supplied at 5 µM. Afterwards, samples were centrifuged for 20 min at 30 °C, 21,000 x *g*. For experiments without MLVs, proteins were incubated in HRB with or without 2.5 mM of indicated nucleotide, incubated for 30 min at 30 °C and centrifuged for 45 min, 436,000 x *g* at 30 °C. In both cases, the supernatant (S) of each reaction was collected and supplemented with an appropriate amount of 6x Laemmli SDS sample buffer (Thermo Fisher Scientific, USA). The pellet (P) was resuspended in 50 µl 1x Laemmli SDS sample buffer. Between 15-25 µl of pellet and supernatant samples were applied to SDS-PAGE using SurePAGE™ Bis-Tris 4-20 % gels (GenScript, USA). Upon protein separation, gels were stained with Quick Coomassie® Stain (Serva, Germany) and protein bands were recorded in a ChemiDoc MP or ChemiDoc Go system (BioRad, USA).

#### ATPase assay

For ATPase assays, we employed the Malachite Green Phosphate Assay Kit (Sigma Aldrich, USA) according to the manufacturer’s instructions. Proteins at indicated concentrations were incubated in ATPase reaction buffer (25 mM HEPES [pH 7.4], 150 mM KCl, 1 mM MgCl_2_) and if appropriate, supplemented with 1 mM ATP or GTP. Samples were incubated at 30 °C and measurements were performed either as endpoint measurements after 2 h or in a time course manner (15, 30, 60, 120 min after ATP addition) of 1/8 dilutions and OD_620_ values were recorded in a Spectramax M2e (Molecular Devices, USA) plate reader. As controls, samples containing no nucleotide or nucleotide only (without protein) were included.

#### Electrophoretic mobility shift (EMSA) assay

For EMSA assays, test DNA sequences were amplified by PCR using Q5 polymerase (NEB, USA) with primer pairs (oBS249+oBS253 for P_corMR_ and oBS627+oBS729 for P_parMR_) that each contained one unlabeled (oBS253 and oBS729) and one 5’ Alexa488-labelled primer (oBS249 and oBS627). PCR products were purified by PCR clean-up using Monarch^®^ PCR & DNA Cleanup Kit (NEB, USA) and stored at-20 °C until further use. 150 ng DNA per sample was then incubated for 30 min at RT with indicated concentrations of CorR or ParR in EMSA reaction buffer (ERB; 25 mM HEPES [pH 7.4], 150 mM KCl, 1 mM MgCl_2_, 1 mM DTT, 0.1 mg/ml BSA, 0.1 mg/ml UltraPure™ Salmon Sperm DNA Solution (Thermo Fisher Scientific, USA), 5% glycerol). Samples were then loaded onto 5% Mini-PROTEAN® TBE Gel (BioRad, USA) without any loading dye and run in 1x TBE buffer (130 mM Tris, 45 mM boric acid, 2.5 mM EDTA) at 100 V for 1 h. Finally, Alexa488 signal was recorded in a ChemiDoc MP system (BioRad, USA).

#### in vivo CorM filament angle measurement

To measure the angle of CorM filaments in *Anabaena* cells, we used two approaches. For the first, we initially straightened *Anabaena* cells expressing mNG-CorM from obtained images using an in house ImageJ script (“straighten_2D”), and filament angles were then measured using CytoSpectre (*81*). Alternatively, angles were measured manually in ImageJ using the angle tool by following an imaginary line from one cell pole to the other and then following each individual mNG-CorM filament from the point of contact with this imaginary pole-to-pole line. Both methods resulted in comparable results.

#### Sample preparation for cryo-EM

For CorM and CorMR samples, 5 µM of CorM alone or 5 µM of both CorM and CorR proteins were incubated in a buffer containing 25 mM HEPES (pH 7.4), 150 mM KCl, 1 mM MgCl₂, and 2 mM ATP for 2 hours at 30 °C. The samples were diluted 1/10 in the same buffer and 3 µL of sample were applied onto glow-discharged 200-mesh holey carbon girds (Quantifoil Micro Tools, Au R2/2).

In the case of ^Δ1-40^CorM, 5 µM of the protein was pre-mixed in the same buffer but without ATP. ATP was only subsequently added to this mixture and 3.5 µl of the sample were applied onto glow-discharged 200-mesh holey carbon girds (Quantifoil Micro Tools, Cu R1.2/1.3) after an incubation of 1 min at RT. These two modifications were necessary as ^Δ1-40^CorM otherwise readily formed extensive, interconnected protein networks within a few minutes after ATP addition that were unsuitable for cryo-EM analysis. All the samples were vitrified in liquid ethane (-185 °C) after 3-3.5s back-side blotting using a Leica GP2 plunger (Leica Microsystems) maintained at 80% humidity and 25 °C (chamber conditions). The grids were stored in liquid nitrogen conditions until imaging.

#### Cryo-electron microscopy

Single-particle datasets were acquired under cryogenic conditions on a Glacios TEM microscope (Thermo Fisher Scientific) operating at 200 kV equipped with Falcon 4i direct electron detector. The automated data collection was set up using EPU version 3.7.0 (Thermo Fisher Scientific).

The nominal magnification was set to 150,000x, resulting in a pixel size of 0.931 Å for the CorMR dataset and 0.9551 Å for the CorM and ^Δ1-40^CorM datasets. The defocus range was set to-1.0 to-3 μm. For all three samples, single-particle datasets were acquired in 4,096×4,096px movies in EER mode, with a cumulative dose ranging from 40-80 e^-^/Å^2^.

#### Single-particle image processing

All the processing steps were performed in CryoSPARC v4.4. For all the datasets, 4,096×4,096px movies in EER mode were motion corrected using patch motion correction, and patch CTF estimation was used to estimate CTF using default patch size in cryoSPARC. Manual curation was done to remove empty micrographs and micrographs containing crystalline ice. The subsequent steps used to process each dataset are summarized below and the details are provided in the schematics presented in Fig. S6.1-6.3.

For the CorMR dataset, initial particle picking was performed using filament tracer with a diameter of 90 Å on 2,305 manually curated micrographs. Particles were initially extracted with a box size of 256 px, followed by 2D classification. Three classes were used as template to perform filament tracing with a diameter of 100 Å. Particles were extracted with a bigger box size (512px) and subjected to 2D classification. Using selected 2D classes (22 classed out of 100), ab-initio reconstruction and homogeneous refinement were performed without symmetry (C1 symmetry). The initial map obtained from homogeneous refinement was used to determine the helical twist and helical rise estimates using the online helical indexing tool (HI3D-Helical Indexing using cylindrical projection of a 3D map) (*82*). Helical refinement was performed using an estimated helical twist of-18° and a helical rise 46.5 Å. Local refinement and local CTF refinements were performed after helical refinement. Two rounds of homogeneous refinement were performed after removing duplicate particles to obtain a final map at 3.92 Å with 124,109 particles, measured at the 0.143 FSC criterion.

For the ^Δ1-40^CorM dataset, particle picking was performed using filament tracer with a diameter of 100 Å on 1,200 manually curated micrographs. Particles were extracted with a box size of 512px and subject to 2D classification. Using selected 2D classes (10 classed out of 75), ab-initio reconstruction and homogeneous refinement were performed with C1 symmetry. The initial map obtained from homogeneous refinement was used to determine the helical twist and helical rise estimates using online helical indexing tool (HI3D-Helical Indexing using cylindrical projection of a 3D map) (*82*). Helical refinement was performed using the estimated helical twist of-15.73° and helical rise 47.3 Å. The map from helical refinement had 287,700 particles, 3D classification was performed to get 3 different classes. One class with good looking map containing 44.2 % of the particles (127,289 particles) was selected and used for final helical refinement to obtain a 4.42 Å map, measured at the 0.143 FSC criterion.

For the CorM dataset, initial particle picking was performed using filament tracer with a diameter of 100 Å with 5 Å separation distance on 2,305 manually curated micrographs.

Particles were extracted with box size of 512 px and 2D classification was performed. 22 Classes out of 100 classes were used as template to perform filament tracing with a diameter of 100 Å and 1 Å separation distance. Particles were extracted with a box size of 360px and subjected to rigorous 2D classification (4 rounds). Using selected 2D classes, ab-initio reconstruction and homogeneous refinement were performed with C1 symmetry. At this stage, the particles were re-extracted with a bigger box size (512px). One more round of 2D classification was performed to remove bad particles. Using selected 2D classes, ab-initio reconstruction and homogeneous refinement were performed with C1 symmetry. The initial map obtained from homogeneous refinement was used to determine the helical twist and helical rise estimates using online helical indexing tool (HI3D-Helical Indexing using cylindrical projection of a 3D map) (*82*). Helical refinement was performed using estimated helical twist of-18.5 ° and helical rise 46.5 Å. The map from helical refinement had 197,123 particles, 3D classification was performed to get 3 different classes. Two classes with good looking map containing 78.3 % of the particles (154,726 particles) were combined and used for final helical refinement job to obtain a 7.14 Å map, measured at the 0.143 FSC criterion.

#### Model building and refinement

All model and EM-density visualizations were done in UCSF ChimeraX version 1.9 (2024-12-11) (*78*). Model building and refinement was performed using the ISOLDE plugin (v.19) (*83*) in ChimeraX.

To build a model of the CorM filament, we first predicted CorM using the AlphaFold3 (AF3) (*77*) webserver, either in its APO-form or bound with ADP or ATP. These predicted models were then rigid body-fitted into the EM density of the filament structure obtained from the CorMR sample. This revealed that the APO-model showed the best fit. However, as there was clearly a density for the nucleotide visible, we manually placed an ADP molecule into the EM-density. As we only observed densities for residues Serine 43 to Asparagine 378 we removed the remaining residues.

To generate a filament-like model, we then rigid body-fitted 6 copies of the generated CorM(43–378)-ADP monomer into the EM-density of the filament. 5 non-crystallographic symmetry copies of the monomers were restrained to a centrally positioned (working) monomer (i.e. a monomer that was surrounded by 5 copies, hence having all its contact sites occupied). Fit and geometry of the working monomer were iteratively checked and locally refined in ISOLDE. Finally, the NCS copies were replaced with the working copy monomer to obtain the final CorM-filament model.

The accuracy of the model was checked using MolProbity (*84*). The interface interactions within CorM filament structures were determined using PDBePISA (Proteins, Interfaces, Structure and Assemblies) (*52*).

#### Plunge freezing Anabaena cells

*Anabaena* WT culture (1.5 ml) was sedimented using a table-top centrifuge for 5 min at 5000 x *g* and resuspended in 1 ml of fresh BG11 medium. 4 µl of cell suspension was applied onto glow-discharged 200-mesh holey carbon grids (Cu R2/2; Quantifoil Micro Tools, Germany). The grids were blotted for 3 s on the backside and plunge frozen into liquid ethane using a Leica GP2 plunger (Leica Microsystems, Germany) maintained at 28 ^°^C and 90 % humidity. In addition, well-grown *Anabaena* Δ*cse* cells were scrapped of a BG11 agar plate and resuspended in 1ml of fresh BG11 medium. 3.5 µl of the cell suspension was applied on the same EM grids as described above and blotted for 12 s from the back using a Vitrobot plunge-freezing robot (Thermo Fisher Scientific, USA), in which one blotting paper was replaced with a Teflon sheet. The grids were stored in liquid nitrogen and subsequently used for Cryo-FIB milling.

#### Lamella preparation by Focused Ion Beam (FIB) milling

Lamellae from *Anabaena* WT were prepared using a second-generation Aquilos (Aquilos II) instrument (Thermo Fisher Scientific, USA). The instrument was controlled via the xT user interface and the MAPS software version 3.25 (Thermo Fisher Scientific, USA). FIB milling was performed at 30 kV, with milling progress monitored using the scanning electron microscopy (SEM) beam operated at 25 pA and 2-5 kV. Prior to milling, specimens were sputter-coated with platinum for 30 s at 30 mA and 10 Pa using the built-in sputter coater. This was followed by gas injection system (GIS) deposition of 1.5-2 µm of metalorganic platinum. Tile-scan overview images were acquired before and after coating using the MAPS software.

Careful lamella placement was critical due to the characteristic rounded, “pearls-on-a-string” morphology of the cyanobacteria. Lamella sites were positioned below the central axis of the cellular chain to avoid gaps between adjacent “pearls”, which could otherwise cause lamella breakage. Placing lamellas too high would create holes in the lamella due to the gaps between cells, while placement too low would prevent adequate GIS coating thickness because the spherical cell shape shields lower regions.

Each lamella was subsequently milled using rectangular patterns and progressively reduced in thickness to ∼200–250 nm. Milling proceeded in several steps, sequentially decreasing both lamella width and milling current to achieve a symmetrical, stair-like anchor. Specifically, lamellae were initially thinned to 3 µm using a 1 nA beam, then reduced to 2 µm at 0.5 nA, and further to 900 nm at 0.1 nA. Final thinning to 500 nm was performed at 50 pA, and each lamella was brought to ∼200 nm thickness using a 30 pA milling current. Due to the relatively small size of the lamellae, no over-or under-tilt milling was required to correct wedge-shaped geometry. With a depth of only 5–6 µm at most, no appreciable wedge formation occurred.

Lamellae through plunge-frozen *Anabaena* Δ*cse* filaments were prepared analogous, but using the SerialFIB software package (*85*) on a Zeiss Crossbeam 550 FIB-SEM dual-beam instrument (Carl Zeiss Microscopy, Germany) equipped with a VCT500 cryo-transfer system and a copper-band cooled mechanical cryo-stage (Leica Microsystems GmbH, Austria) (*86*).

Prior transfer to the FIB-SEM, grids were sputter-coated with a ∼8 nm thick layer of tungsten using an ACE600 cryo-sputter coater (Leica Microsystems GmbH, Austria). An organometallic platinum precursor layer was deposited onto each grid via the GIS, and the SEM (3-5 kV, 58 pA) was used to identify suitable milling targets. Milling patterns were placed and executed via the SerialFIB software package, aiming for a final lamellae thickness of ∼180 nm by applying gradually reducing currents for rough milling (700, 300, and 100 pA) and for polishing (50 pA).

All prepared samples were stored in Autogrid boxes in liquid nitrogen until cryo-ET data acquisition.

#### Cryo-ET data acquisition and image processing

Data acquisition on cryo-lamellae of *Anabaena* WT was performed using a Thermo Fisher Scientific (USA) Titan Krios G3i transmission electron microscope, operated at 300 kV in nanoProbe energy-filtered TEM (EFTEM) mode. For acquisition a Gatan K3 BioQuantum direct electron detector was used. Zero-Loss Peak (ZLP) and energy filter tuning using a 20 eV slit width were carried out with DigitalMicrograph 3.52.392.0 (Gatan). Coma-free alignment and astigmatism correction were performed using SerialEM 4.1.10 (*87*). For medium-magnification overview maps of lamella, a pixel size of 13.74 Å at a nominal magnification of 6,500× was used. For tilt series acquisition, the camera was operated in counting mode with hardware binning and dose fractionation, acquiring eight frames per tilt. The total electron dose applied was 160 to 180 e⁻/Å², distributed across 41 images following a 3° increment tilt scheme. All tilt series were collected using a dose-symmetric acquisition scheme (*88*), starting from the lamella pre-tilt angle and spanning a range of −60° to +60° at a defocus of −1.5 to −3.5 µm and the pixel size was set to 1.38 Å at a nominal magnification of 64,000×. Tilt series were acquired with SerialEM 4.1.10 (*87*) and PACE-tomo 1.4.4 (*89*) as 5,760 x 4,092 pixel movies with 8 frames.

The movies were aligned on-the-fly during data acquisition using frame alignment in SerialEM and saved as mrc stacks. Tomograms of *Anabaena* Δ*cse* lamellae were recorded on an identically equipped microscope and with the same software packages, yet with a pixel-size of 2.68 Å at the specimen level, defocus-6 µm and a total dose of ∼140 e⁻/Å², distributed across 41 images following a 3° increment, dose-symmetric tilt scheme.

The tilt series mrc image stacks from SerialEM were used to perform patch-based tilt-series alignment in AreTomo2 (version 1.3.4) (*90*) with 6×4 patches. 8x binned tomograms were reconstructed (510×720×250 pixels) using the Weighted-Back-Projection method in AreTomo2 or using the IMOD software package (*91*, *92*). In order to fill the missing-wedge information, enhance the contrast and to facilitate visualization of filaments in the z-direction, bin8 tomograms were filtered using IsoNet (*93*). Tomogram visualization and filament diameter measurements were performed using IMOD (*91*).

#### Scanning electron microscopy (SEM)

For SEM analysis, cells were grown on BG11 plates, scraped off, and re-suspended in 1 ml MilliQ H₂O. Cells were pelleted by centrifugation (5000 × *g*, 4 min, RT), washed once by centrifugation with 1 ml Milli-Q H₂O, and then re-suspended in 1 ml of 0.1 M Tris base/HCl buffer containing 2.5% glutaraldehyde. Cells were fixed o/n at 4 °C. The next day, cells were washed 2x with 1 ml MilliQ H₂O by centrifugation (5000 × *g*, 4 min, RT). After washing, cells were spotted for 30 seconds onto 12 mm round glass coverslips (Electron Microscopy Sciences, USA) previously coated overnight with 0.01% poly-L-lysine (Sigma-Aldrich, USA).

Subsequently, cells were subjected to an ethanol dehydration series by sequential incubation on the coverslips in 6-well plates with 40 %, 60 %, 80 %, and 100 % EtOH, each for 15 minutes. Samples were then critical point dried using EM CPD300 (Leica Microsystems) and sputter-coated with gold to a thickness of 5 nm using EM ACE600 coating device (Leica Microsystems) prior to imaging using a FE-SEM Merlin VP Compact (Zeiss, Germany) at 5 kV with a secondary electron detector.

#### Protein sequence analyses

We assembled a database of complete cyanobacterial genomes, aiming to represent the overall diversity of the phylum included in the NCBI Genome database (*94*) by including up to five species per genus when available, resulting in a final dataset of 128 species. To identify homologs of ParM, we used the *Nostoc* sp. PCC 7120 protein WP_044522477.1 as a query in a jackhmmer search (HMMER v3.3.2) (*95*) against our cyanobacterial database. All hits were aligned using MAFFT v7 (L-INS-i algorithm) (*96*), and the resulting alignment was manually curated to remove outliers. From this alignment, we generated a hidden Markov model (HMM), which we used for a second round of searches with hmmsearch(*95*). The same approach was applied to identify homologs of MinC (WP_190449770.1) and ParR (WP_010999216.1), both from *Nostoc* sp. PCC 7120, but in the case of ParR, additional analyses were necessary due to its low sequence conservation and variable length across homologs. To address these limitations, ParR hits were filtered based on genomic context. In all reported cases, *parR* was located adjacent to a predicted *parM* gene, and both genes were encoded in the same orientation. Based on this observation, we only retained ParR hits found immediately next to a ParM, and all others were discarded. We also examined the distribution of ParM across Bacteria. Preliminary analyses showed that ParM is unevenly distributed and can be present in some strains but absent in others within the same species. For this reason, we used a different strategy than the one applied to Cyanobacteria. We searched all complete bacterial chromosomes and plasmids deposited in RefSeq (*97*), excluding cyanobacterial entries, using hmmsearch with the ParM HMM and an E-value cutoff of 1e−6. This search retrieved 3,693 chromosomal and 2,298 plasmid hits, the majority of which belonged to the phyla Pseudomonadota and Bacillota, which together accounted for over 95% of all identified sequences. To reduce redundancy, we randomly selected one hit per genus, aligned all sequences using MAFFT v7 (L-INS-i algorithm) (*96*), and manually removed highly divergent or poorly aligned entries. The resulting curated dataset consisted of 150 chromosomal and 78 plasmid sequences. Given the poor sequence conservation of ParR described above, we did not rely on direct sequence similarity to search for its homologs across Bacteria. Instead, we extracted the two neighboring genes (upstream and downstream) of each ParM hit, provided they were in the same orientation, and predicted their three-dimensional structures using AlphaFold2 (*55*). ParR candidates were then manually annotated based on structural similarity to the ParR reference sequence. The presence of MinC homologs in other bacterial phyla was assessed as previously described, but using a database containing taxa representative of most bacterial phyla (*98*). To predict amphipathic helices in all the selected ParR sequences, we used Amphipaseek (*51*). Helical wheels were generated using NetWheels (*99*) and amphipathicity was calculated using PMIpred (*100*). All protein alignments for ParM, ParR and MinC were performed using MAFFT (*96*), and columns containing more than 20% gaps were removed using TrimAl (*101*). Alignments were visualized in JalView (*102*). Sequence logos were generated using WebLogo 3 (*103*). To analyze and display sequence conservation on protein structures, we used ChimeraX (*104*) in combination with the AL2CO algorithm (*105*).

#### Phylogenetic analyses

To reconstruct the phylogenies of ParM and MinC, we used all homologs identified in Cyanobacteria and in the Bacteria datasets. Phylogenetic inference was performed with IQ-TREE v2 (*106*), using ModelFinder (*107*) to determine the best-fitting substitution model, and ultrafast bootstrap approximation with 10,000 replicates (*108*) and a minimum bootstrap correlation coefficient of 0.9999 to assess branch support. To improve resolution within the cyanobacterial clade, we reconstructed a second phylogeny based on concatenated alignments of ParM and ParR sequences, processed using the same strategy as described above. We also built a species tree for Cyanobacteria using two conserved, assumed to be vertically inherited markers: the DNA-directed RNA polymerase subunits β (BAB77960.1) and β′ (BAB77961.1). Each sequence was aligned and trimmed separately using the same method explained above, and the two alignments were concatenated for phylogenetic analysis. The resulting tree was rooted using four species from *Candidatus* Melainabacteria, the sister clade of Cyanobacteria. All phylogenetic trees were visualized and annotated using the iTOL web server v6 (*109*).

#### In vivo CorM filament tracking

mNG-CorM or mNG-^Δ1-40^CorM filaments were tracked from time lapse movies with images recorded usually every 5 or 10 seconds. Using Fiji (*110*), we used the line tool (width 3) and followed the track of individual filaments of which we generated kymographs using the Reslice plugin. Growth or shrinkage velocities were then calculated by distance traveled over time (nm/s).

#### In vitro CorM filament tracking

Filament ends were manually tracked using the Tracking plugin in Fiji (*110*). Distance between positions were then calculated using a custom Python script.

#### Intrinsically disordered region (IDR) prediction

Unstructured domains of cyanobacterial CorM or ParM homologs was performed using ALBATROSS (*111*) on a Google Colab notebook (IDRome constructor v. 3.0).

#### AlphaFold-Multimer predictions

For prediction of protein-protein interactions, we tested CorM against the whole *Anabaena* proteome (RefSeq ID: GCF_000009705.1) as extracted from NCBI. This ‘candidates’ list was tested to predict interactions against the ‘bait’ CorM using AlphaFold-Multimer (v.2.3) (*54*, *77*). This pipeline produced five protein complexes per protein pair. Postprocessing of the results includes a filtering step, which removes protein complexes with an ipTM standard deviation larger than the 1.5xIQR outlier threshold, i.e. 0.095, indicate highly diverse structure predictions. Unless stated otherwise, all depicted complex predictions and their corresponding PAE plots are the top-scoring relaxed model generated by Alphafold-Multimer (v3) 64 and visualized in ChimeraX.

#### Prediction of protein 3D structures

All shown Protein 3D structures were predicted using AF3 (*77*) and further analyzed using UCSF ChimeraX (v. 1.9) (*78*). For AF3 predictions, the respective top ranked model is shown in cartoon form and colored by pLDDT score confidence together with the corresponding Predicted Aligned Error (PAE) plot and the ipTM/pTM scores.

#### Image and data analysis

Microscopy data analysis was performed using FIJI software (ImageJ2 for Mac; v. 2.14.0/1.54f) (*110*). Time series micrographs were routinely processed using the ImageJ gaussian blur (0.5) and walking average plugins, averaging four consecutive frames. Contrast was optimized using the Enhance Contrast command (0.05-0.35% saturated pixels; normalize, process all slices) and, where appropriate, background was subtracted using the Subtract Background command (50 pixels Rolling ball radius) from ImageJ.

Quantification of protein signals from SDS gels was done using Fiji or Image Lab (BioRad, USA) software. Protein signal intensities of the loaded pellet and supernatant samples were recorded and extrapolated to the overall sample volume. Values are given as percentage of proteins in the pellet or supernatant in respect to the whole protein amount of each reaction.

#### In vivo timelapse cell division tracking

To analyze the temporal localization of mNG-CorM, time-lapse movies of individual cells were manually stabilized using a custom Fiji macro to compensate for cell growth, division, rotation, and lateral drift. Frame intensities were normalized to ensure accurate assessment of signal localization, and each cell was cropped into a 100×100 pixel region. Initial stabilization was performed by aligning movie frames to a line region of interest (ROI) positioned along the cell’s lateral axis, guided by mNG-CorM and thylakoid membrane fluorescence signals, to correct for rotation. After stabilization, a kymograph was generated for each cell by drawing a line along the longitudinal axis, spanning from pole to pole, with a line width of 1.25 µm to average the mNG-CorM signal and compensate for gaps in fluorescence. These kymographs were temporarily aligned based on the onset of cell division, defined by the sudden separation of the mNG-CorM signal into two diverging trajectories. The aligned kymographs were then stacked and averaged to produce the composite final image.

#### Statistical analysis

For all plots throughout this work, error bars indicate the standard deviation (SD).

For TIRF microscopy of *in vitro* experiments, we define a biological replicate as a reaction performed in a separate reaction chamber. In some cases, replicates were performed on the same cover glasses, which can hold up to three reaction chambers. A normal distribution was assumed for all replicates.

## Supplementary Figures

**Fig. S1:**
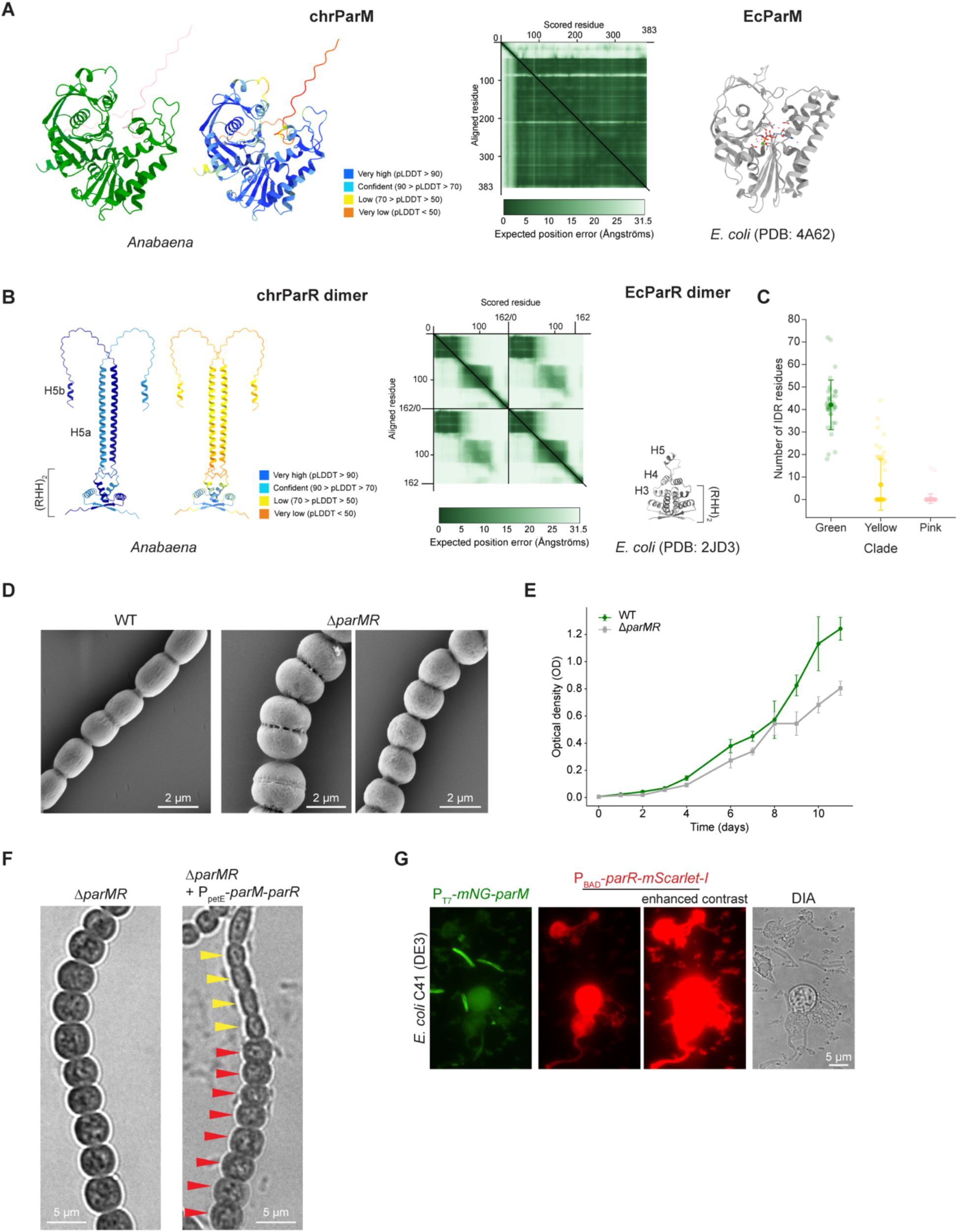
Properties of the chromosomal ParMR (**A**) AlphaFold3-predicted structure of chromosomal ParM (chrParM; green) from *Anabaena* (pTM score: 0.83) and the resolved X-ray structure of plasmid-encoded ParM from *E. coli* (PDB ID: 4A62). (**B**) AlphaFold3-predicted structure of a chromosomal ParR (chrParR; blue/purple) dimer from *Anabaena* (pTM score: 0.34; ipTM score: 0.34) and the resolved X-ray structure of the plasmid-encoded ParR dimer from *E. coli* (PDB ID: 2JD3). For A and B, predicted aligned error (PAE) maps and plDDT score-colored structure predictions for AlphaFold3 predictions are also shown. Note that we show both plDDT score-colored and individually colored AlphaFold3-predicted structures to better illustrate the individual protein chains within the dimer. (**C**) Mean amino acid residues ± SD within the N-terminal IDR from ParM homologs from the green (42.05 ± 11.03), yellow (9.66 ± 11.32) and pink (0.71 ± 2.149) clades of cyanobacterial ParM’s showing that only the green clade has any substantial N-terminal IDR. (**D**) Scanning electron microscopy images of (left) *Anabaena* WT and Δ*parMR* revealing cell division defects in 30 % of actively dividing Δ*parMR* cells (6 of 20 cells). These cells displayed irregular or incomplete septa, being partially constricted but not fully separated, a phenotype not observed in the WT (16 cells). Left Δ*parMR* image shows defects in proper constrictions, which are not observed in another subset of Δ*parMR* cells as indicated on the right Δ*parMR* image. (**E**) Representative growth curve analysis of *Anabaena* WT and Δ*parMR* revealing that the mutant lacking ParMR has a slight growth defect. OD_750_ values were recorded every day or every other day for a total of 11 days. Afterwards, cultures became too sticky to properly measure the OD_750_ value. Error bars represent the mean ± SD of three technical replicates. The experiment was performed twice with similar results. (**F**) Complementation attempts of the Δ*parMR* phenotype. DIA micrograph of Δ*parMR* carrying a replicative plasmid expressing *parM*-*parR* from the medium-to-strong *petE* promoter (P_petE_). For comparison Δ*parMR* is shown. The swollen phenotype of Δ*parMR* is complemented to WT morphology in a subset of cells (yellow arrow) within a cell filament, likely through high variability of plasmid copy numbers, and thus variability of expression levels between individual cells (*43*, *44*). Red arrows indicate cells of the same cell filament with Δ*parMR* mutant phenotype. (**G**) Micrographs of *E. coli* C41 (DE3) co-expressing mNG-ParM from P_T7_ and ParR-mScarlet-I from P_BAD_ grown over night in TB medium supplemented with 0.02 % L-arabinose and 0.25 mM IPTG at 37 °C. Cells containing ParR-mScarlet-I signal are strikingly malformed, forming huge balloon-like structures. These cells lack any discernible mNG-ParM signal, indicating that they cannot tolerate both proteins together for extended periods. Micrograph with enhanced contrast reveals red fluorescent cell debris, likely originating from burst cells expressing ParR-mScarlet-I. mNG-ParM localizes as filamentous strings in cells with WT-like morphology that lacked ParR-mScarlet-I signal, suggesting that specifically ParR is responsible for the aberrant cell phenotype.

**Fig. S2:**
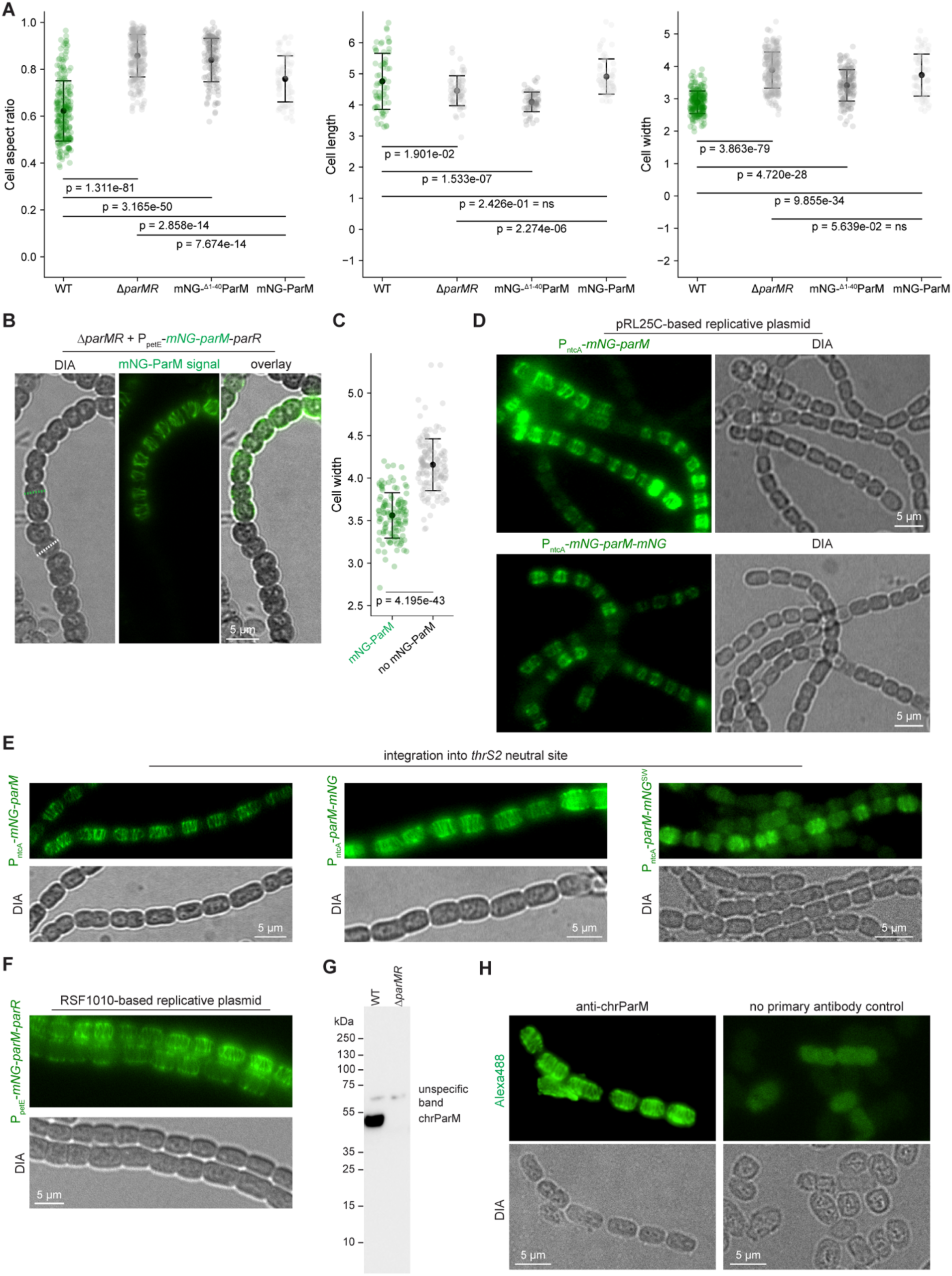
Cortical localization of chrParM filaments *in vivo* (**A**) Scatter plots showing the mean cell property values ± SD for cell width (WT: 4.8 ± 0.96 µm; Δ*parMR*: 4.53 ± 0.47 µm; mNG-ParM: 3.73 ± 0.65 µm; mNG-^Δ1-40^ParM: 3.42 ± 0.49 µm), cell length (WT: 2.89 ± 0.35 µm; Δ*parMR*: 3.88 ± 0.56 µm; mNG-ParM: 4.91 ± 0.56 µm; mNG-^Δ1-40^ParM: 4.06 ± 0.29 µm) and cell aspect ratio (WT: 0.62 ± 0.13; Δ*parMR*: 0.86 ± 0.09; mNG-ParM: 0.76 ± 0.09; mNG-^Δ1-40^ParM: 0.84 ± 0.09) of *Anabaena* WT, Δ*parMR*, mNG-ParM or mNG-^Δ1-40^ParM expressing cells. WT: n=222 cells; Δ*parMR*: n=259 cells; mNG-ParM: n=67 cells; mNG-^Δ1-40^ParM: n=144 cells. Values for WT and Δ*parMR* are the same as in Fig. 1F-H. (**B**) DIA, mNG fluorescence and merged DIA and mNG fluourescence micrographs of Δ*parMR* carrying a replicative RSF1010-based broad host-range plasmid expressing mNG-ParM-ParR from P_petE_. Only cells with discernible mNG fluourescence appear more WT-like whereas cells lacking mNG fluorescence show Δ*parMR* mutant phenotype, which we confirmed by cell width measurements (indicated by dotted green for mNG-expressing cells and white dotted lines for Δ*parMR* mutant cells). (**C**) Comparison of cell width (in µm) for cells carrying pBS417 (P_petE_-*mNG-parM-parR*) that display detectable mNG-ParM fluorescence (n=119) and for cells lacking any mNG-ParM signal (n=145). (**D**) DIA and mNG fluorescence micrographs of *Anabaena* WT carrying a replicative pRL25C-based cyanobacterial replicative plasmid expressing mNG-ParM (top) or ParM-mNG (bottom) from P_nctA_. (**E**) DIA and mNG fluorescence micrographs of *Anabaena* WT expressing mNG-ParM (left), ParM-mNG (middle) or ParM-mNG^SW^ (sandwich fusion of ParM with mNG, inserting mNG between K209 and G210 flanked by a N-terminal SGSS and a C-terminal ASAS linker) from P_nctA_ integrated into the neutral *thrS2* site (*48*). (**F**) DIA and mNG fluorescence micrographs of *Anabaena* WT carrying a replicative RSF1010-based replicative plasmid expressing mNG-ParM-ParR from P_petE_. (**G**) Anti-ParM western blot of *Anabaena* WT and Δ*parMR* cell lysates showing a distinct signal only for the WT lysates. An unspecific band with higher molecular weight is present in both, WT and Δ*parMR*, thus not reacting with ParM. (**H**) DIA and Alexa488 fluorescence micrographs showing anti-ParM immunofluorescence staining in *Anabaena* WT cells. Control cells were incubated with the secondary antibody only.

**Fig. S3:**
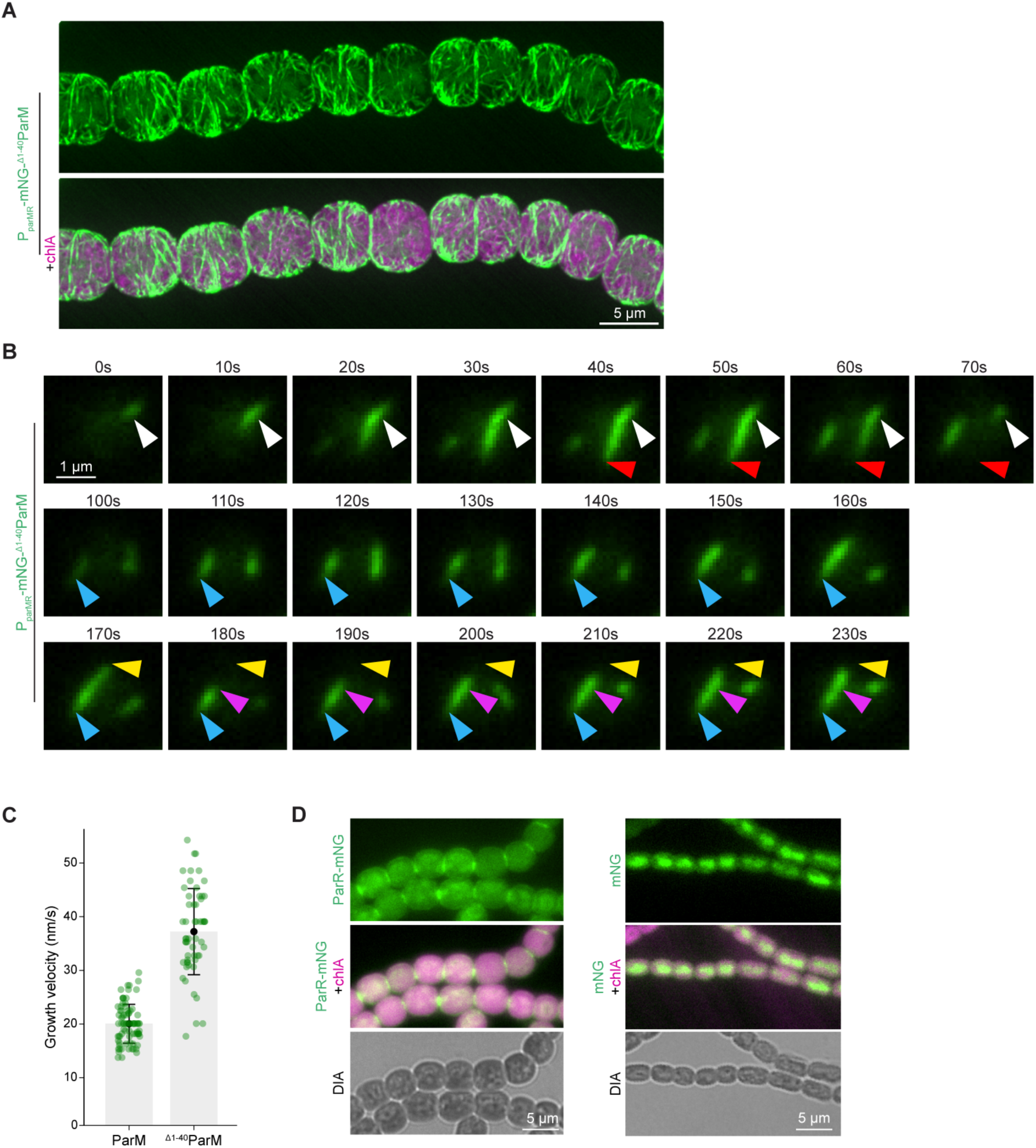
ParM’s N-terminal IDR is not involved in cellular ParM localization or dynamics (**A**) mNG-^Δ1-40^ParM and merged mNG-^Δ1-40^ParM and chlA autofluorescence micrographs of *Anabaena* cells expressing mNG-^Δ1-40^ParM as the sole copy from its native locus. The images show maximum intensity projections and were obtained with a Nikon SoRa spinning disk confocal microscope. (**B**) Time series from Supplementary Movie 3 displaying TIRF micrographs of individual mNG-^Δ1-40^ParM filaments over a period of 230 seconds. White triangles mark the initial position of a single mNG-^Δ1-40^ParM filament and red triangles mark the furthest observed position of this filament during the time series. Blue triangles mark the initial position of another single mNG-^Δ1-40^ParM filament, yellow triangles mark the furthest observed position of this filament during the time series and purple triangles indicate the position this filament retracted to upon a dynamically instable shrinkage event. (**C**) Scatter plot showing the mean ± SD of the growth velocities of mNG-ParM (18.96 ± 3.61 nm/s; n=80) and mNG-^Δ1-40^ParM (35.75 ± 8.19 nm/s; n=58) filaments, revealing the N-terminal IDR is responsible to limit the growth kinetics *in vivo*. (**D**) ParR-mNG, DIA and merged ParR-mNG and chlA autofluorescence micrographs of *Anabaena* expressing ParR-mNG as the sole copy. This strain’s cellular morphology is markedly altered, resembling the Δ*parMR* strain. This suggests that the ParR-mNG fusion is not fully functional, likely because ParR’s C-terminal peptide is essential for interaction with ParM and is sterically hindered by the C-terminal mNG fusion. Note that membrane localization of proteins in *Anabaena* looks mostly diffuse at the periphery as is the case for ParR-mNG. For comparison, *Anabaena* cells expressing soluble mNG are shown on the right side. Micrographs were obtained by a standard Nikon TI2 epi-fluorescence microscope.

**Fig. S4:**
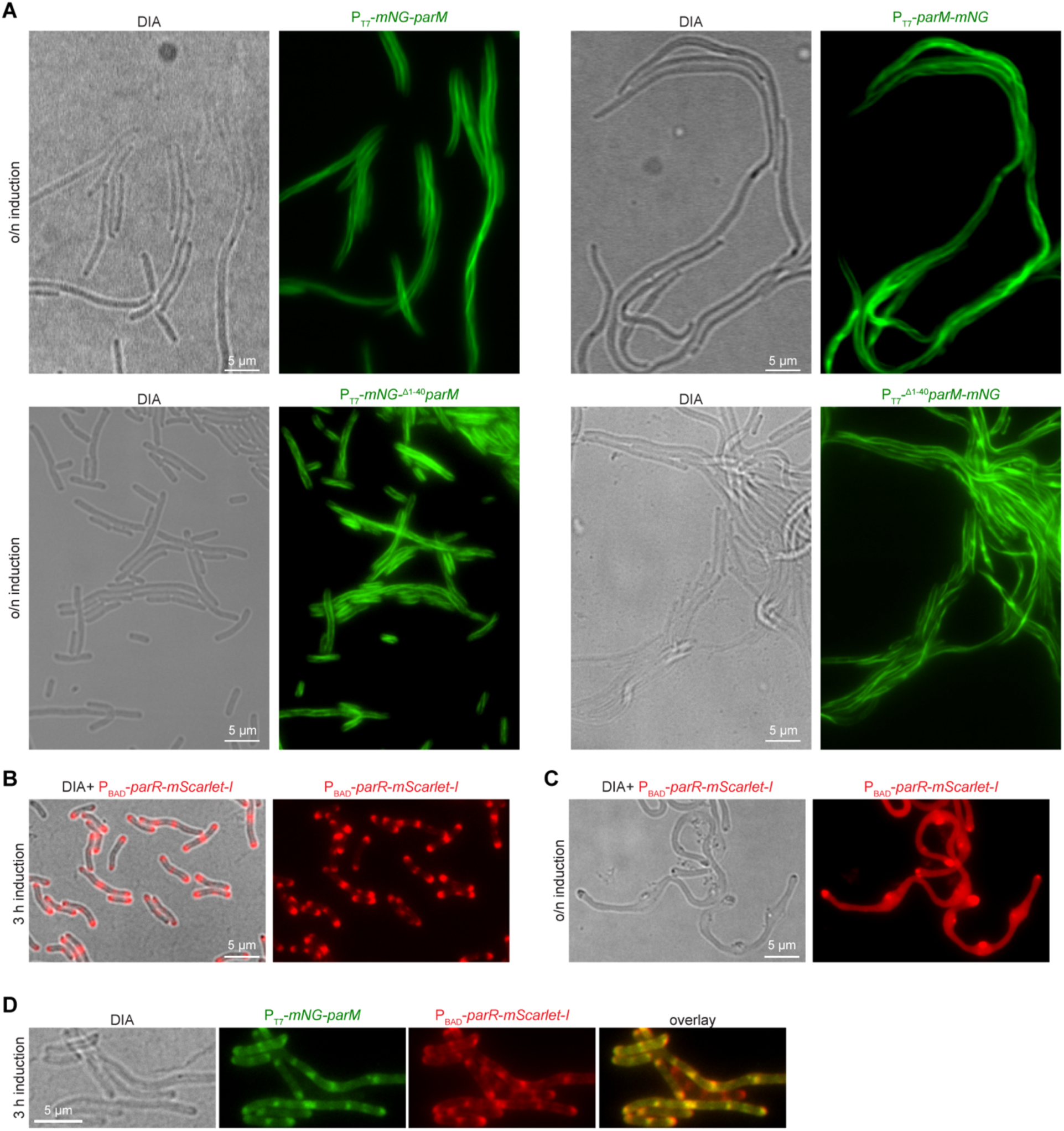
Heterologous expression of chrParMR in *E. coli* (**A**) DIA and mNG fluorescence micrographs of *E. coli* C41 (DE3) expressing mNG-ParM, ParM-mNG, mNG-^Δ1-40^chrParM or ^Δ1-40^chrParM-mNG from P_T7_ grown over night (o/n) in TB medium supplemented with 0.25 mM IPTG at 37 °C. (**B**) mScarlet-I fluorescence and merged DIA and mScarlet-I fluorescence micrographs of *E. coli* C41 (DE3) expressing ParR-mScarlet-I from P_BAD_ grown over night without induction, diluted 1:100 in fresh TB medium supplemented with 0.02% L-arabinose and grown for 3 hours at 37 °C prior imaging. (**C**) mScarlet-I fluorescence and merged DIA and mScarlet-I fluorescence micrographs of *E. coli* C41 (DE3) expressing ParR-mScarlet-I from P_BAD_ grown over night in TB medium supplemented with 0.02 % L-arabinose at 37 °C. (**D**) DIA, mNG fluorescence, mScarlet-I fluorescence and merged micrographs of *E. coli* C41 (DE3) co-expressing mNG-ParM from **P**_T7_ and ParR–mScarlet-I from **P**_BAD_, grown overnight in TB medium, diluted 1:100 into fresh TB supplemented with 0.02% L-arabinose and 0.25 mM IPTG the next day, and incubated for 3 h at 37 °C prior imaging. Note that mNG-ParM initially forms filaments similar to what is shown in (A) but then over time co-localizes with ParR-mScarlet-I.

**Fig. S5:**
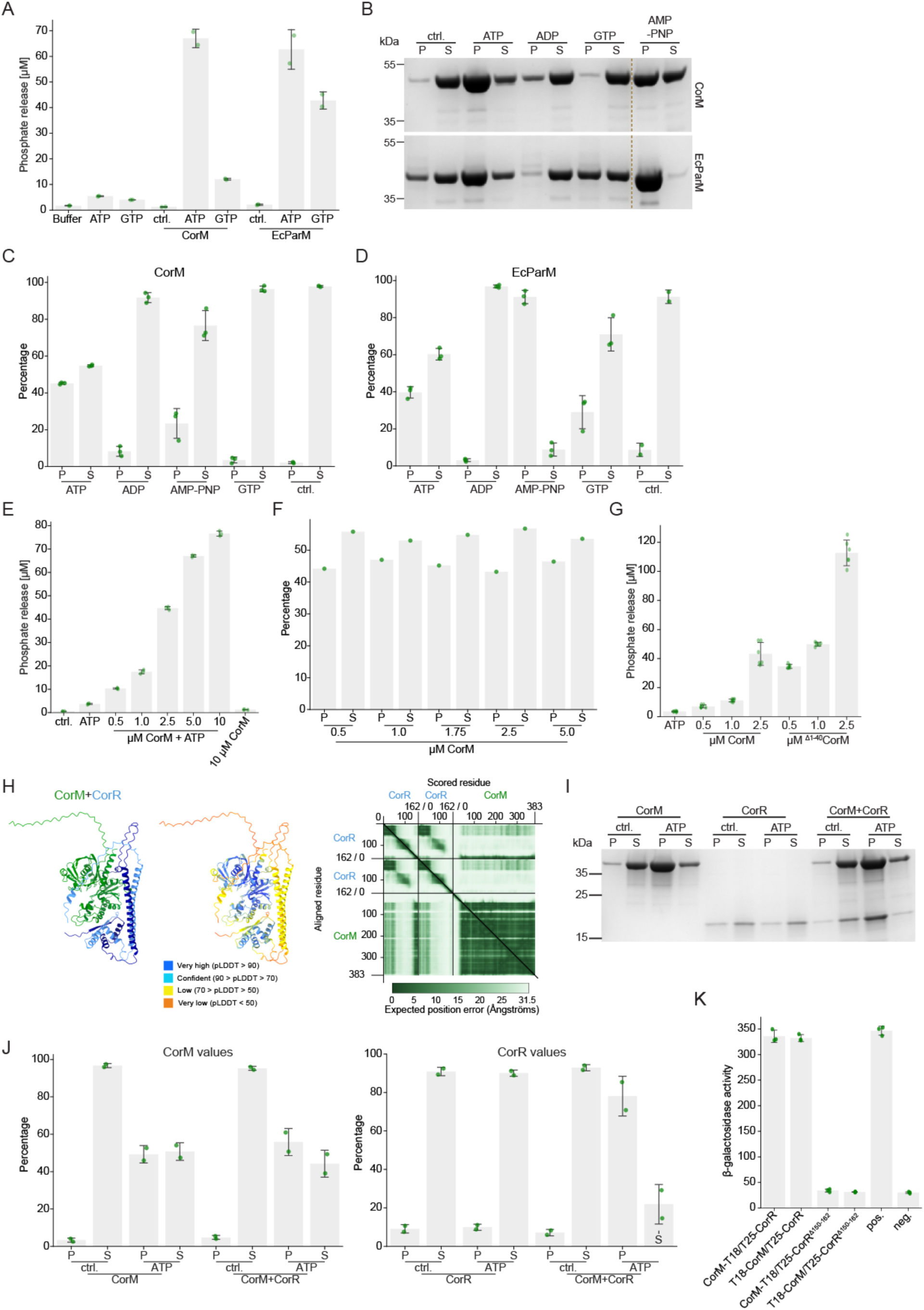
Biochemical properties of CorM reveal functional divergence from EcParM and regulation by the N-terminal IDR (**A**) NTPase activity assay using the Malachite Green Phosphate Assay Kit (Sigma Aldrich) shows that CorM readily hydrolyzes ATP but, unlike the EcParM, is not able to meaningfully hydrolyze GTP. NTPase activity is quantified by the degree of phosphate release during the conversion of NTP to NDP during NTPase activity. Error bars represent the mean ± SD of two independent replicates (n=2). (**B**) Representative Coomassie-stained SDS-PAGE gel from a high-speed ultracentrifugation-based pelleting assay comparing CorM with EcParM (both 5 µM) in the presence of buffer (ctrl.), ATP, ADP, GTP and a non-hydrolyzable ATP analog (AMP-PNP). Nucleotides were provided at 2.5 mM each. 40 % of the total pellet [P] and 11.1 % of the total supernatant [S] samples were separated on the gel. (**C, D**) Quantification of the pelleting efficiency as displayed in (B), confirming that CorM only polymerizes in the presence of ATP but not with GTP or AMP-PNP. Error bars represent the mean ± SD of three independent replicates (n=3). (**E**) ATPase activity assay showing that CorM ATPase activity is a concentration dependent process that seemingly begins to plateau at higher concentrations. Error bars represent the mean ± SD of three independent replicates (n=3). (**F**) The degree of CorM polymerization is not a concentration-dependent process and is not altered upon high-levels of CorM (n=1). (**G**) ATPase activity assay comparing CorM with a mutant lacking the first 40 aa (^Δ1-40^CorM) reveals that the N-terminal IDR limits CorM’s ATPase activity. Error bars represent the mean ± SD of two independent replicates with 3 technical replicates each. (**H**) AlphaFold3-predicted interaction between CorM (green) and a CorR dimer (blue/purple), together with the corresponding PAE map, showing interaction of CorR’s C-terminal α-helix H5b with CorM, as well as a plDDT score-colored CorM-CorR complex (pTM score: 0.59; ipTM score: 0.51). (**I**) Representative Coomassie-stained SDS-PAGE gel from a high-speed ultracentrifugation-based pelleting assay comparing CorM and CorR, alone or in combination (5 µM each), incubated with 2.5 mM ATP or without nucleotide (ctrl.) (n=2). 34 % of the total pellet [P] and 9.7 % of the total supernatant [S] samples were separated on the gel. (**J**) Quantification of the pelleting efficiency from (I), showing pellet and supernatant values for CorM (left graph) and CorR (right graph) individually. CorR is recruited to the pellet through polymerized CorM (ATP-containing sample). Data represent the mean ± SD from two independent experiments (n=2). (**K**) B2H assays testing for the interaction of CorM with CorR from *Anabaena* fused to either the T25 or T18 subunit, including a CorR variant lacking the predicted C-terminal α-helix (amino acids 150-162), which is known to facilitate the interaction of plasmid-encoded ParR with ParM in *E. coli*. CorR lacking amino acids 150-162 is no longer able to interact with CorM. Negative control: CorM-T18 with empty pKNT25 plasmid. Positive control: Zip/Zip control. Error bars represent the mean ± SD of three independent replicates (n=3).

**Fig. S6:**
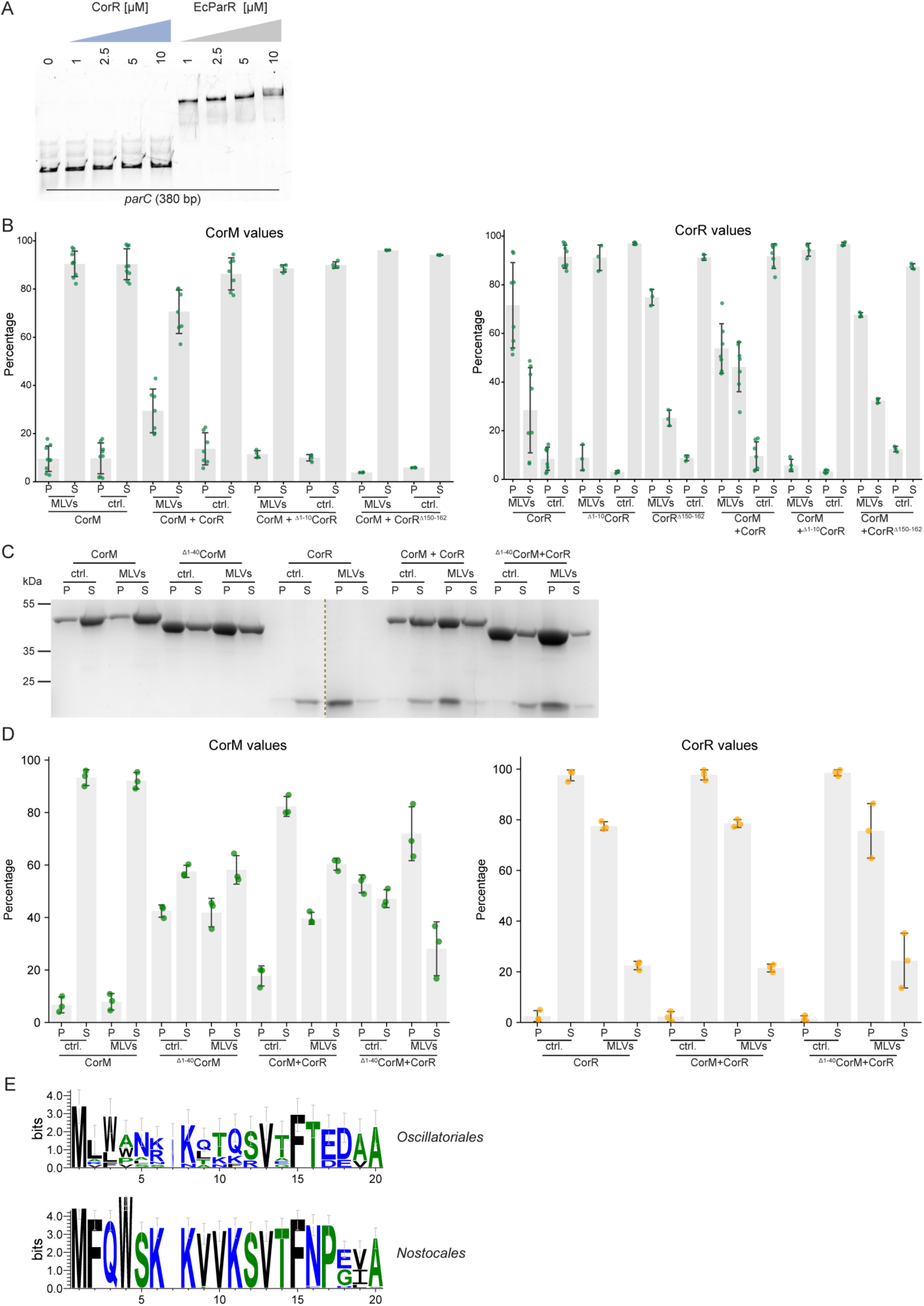
CorR binds to lipid membranes (**A**) EMSA assay with increasing concentrations of CorR or *E. coli* ParR using *parC* (380 bp) as DNA bait. Concentrations in µM are given above the individual lanes. The gel shows a representative example from two independent experiments, displaying Alexa488 fluorescence signals of PCR-amplified DNA regions obtained using fluorescently labeled primers. (**B**) Quantification analysis of the CorM with CorR pelleting assay from Fig. 3C showing the values for CorM and CorR separately. P indicates values for pellet samples and S indicates values for supernatant samples. Error bars represent the mean ± SD from at least three independent replicates. (**C**) Representative Coomassie-stained SDS-PAGE gel from a pelleting assay with indicated protein combinations with or without MLVs (proteins are provided as 5 µM each). 30 % of the total pellet [P] and 12.5 % of the total supernatant [S] samples were separated on the gel. (**D**) Quantifications from pelleting assay shown in (B), showing values for CorM and CorR separately. Error bars represent the mean ± SD from three independent replicates (n=3). (**E**) Amino acid sequence logo (using WebLogo (*103*)) from the multiple sequence alignment shown in Fig. S8, showing only the first 20 aa of CorR’s N-terminus. To enhance readability, CorR sequences from *Oscillatoriales* and *Nostocales* are shown independently, which also highlights that the N-terminal CorR region is strongly conserved among *Nostocales* species and less so among *Oscillatoriales* species.

**Fig. S7:**
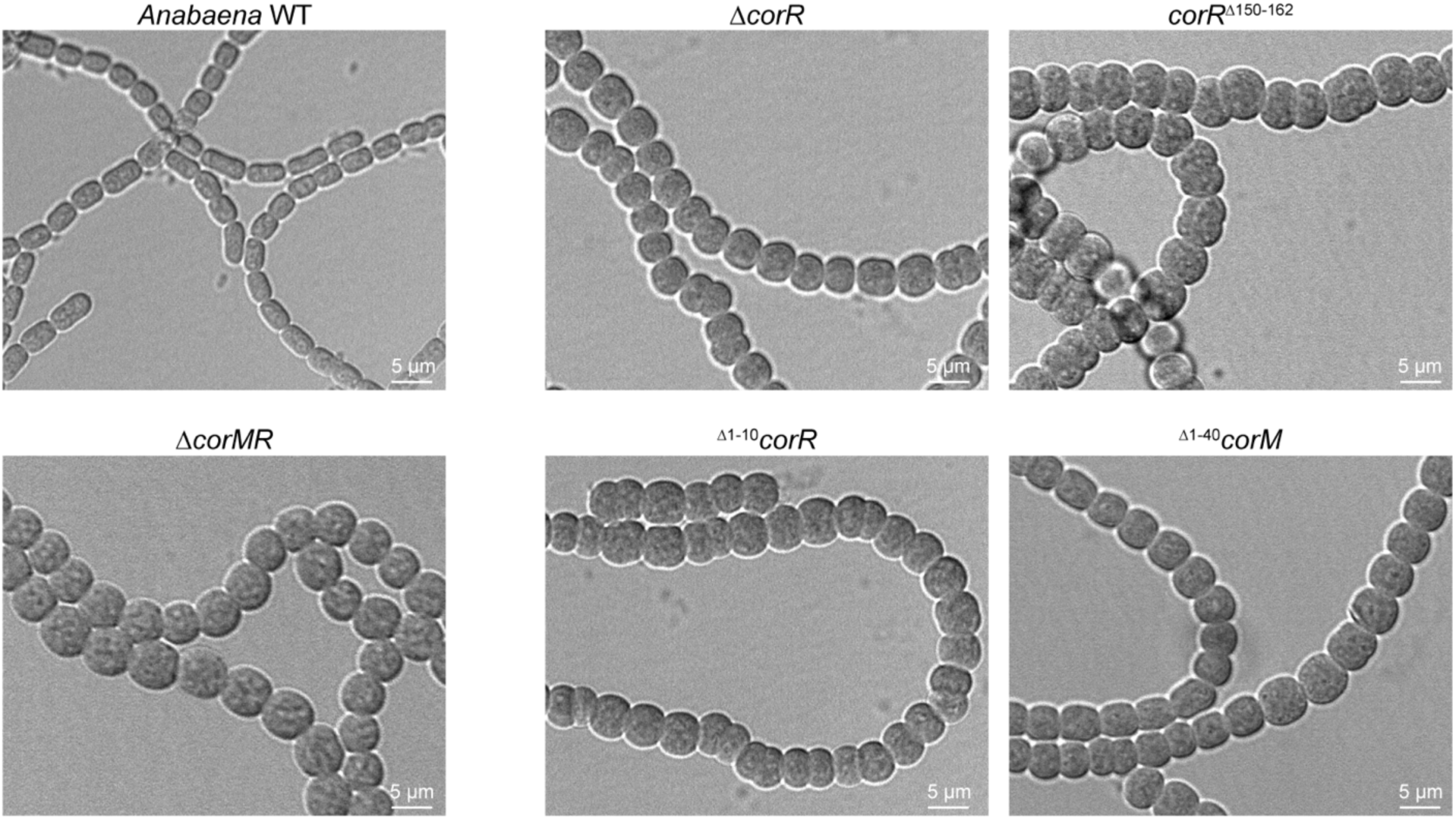
Deletion of CorMR system components and domains DIA micrographs of *Anabaena* WT, Δ*corMR*, and strains lacking *corR* (Δ*corR*), CorR’s N-terminal AH (^Δ1-10^*corR*), CorR’s C-terminal peptide essential for CorM binding (*corR*^Δ150-162^), or CorM lacking the N-terminal IDR (^Δ1-40^*corM*).

**Fig. S8.**
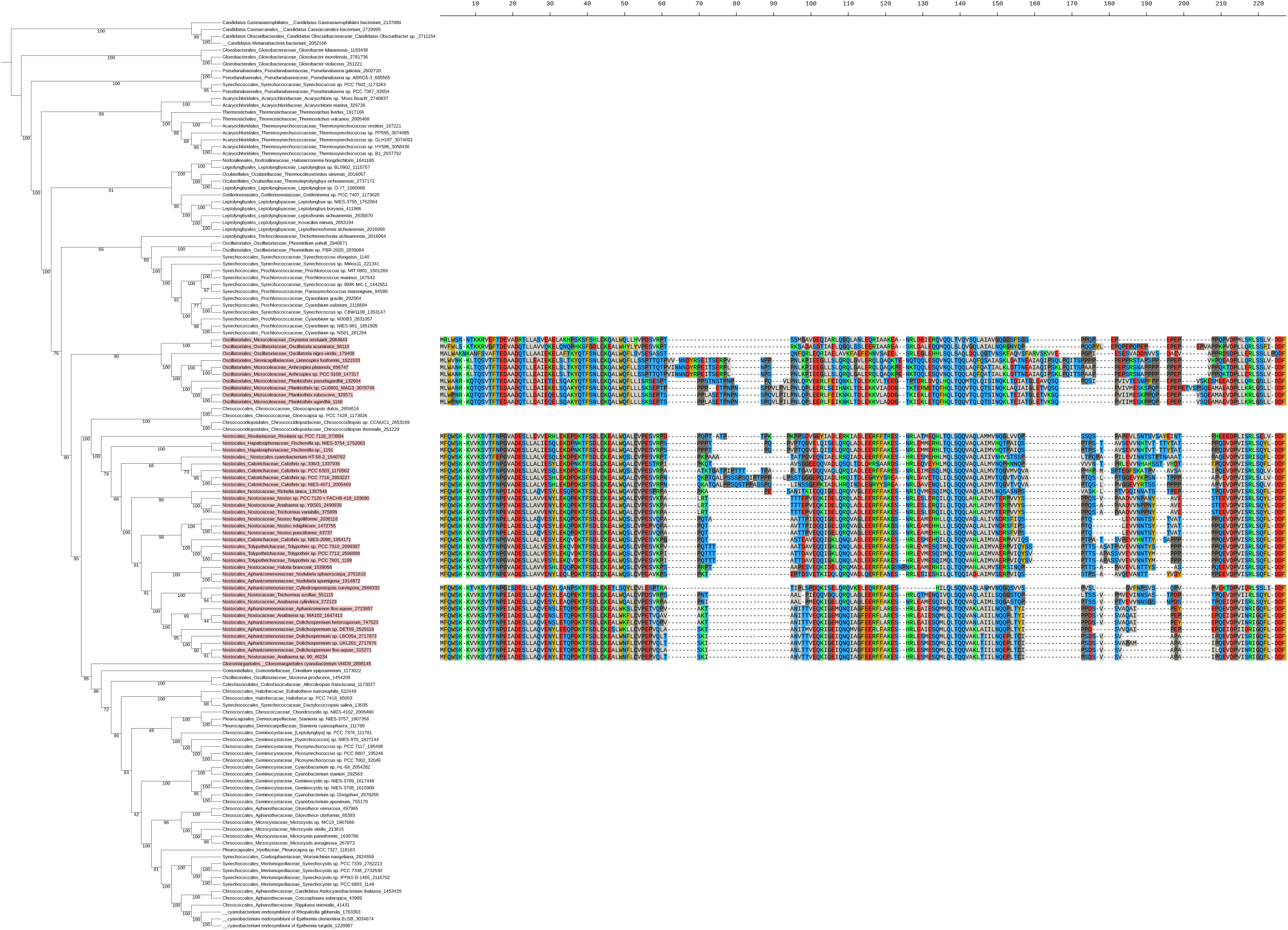
Conservation of CorR’s amphipathic helix Cyanobacterial phylogenetic species tree with multiple sequence alignment of CorR homologs containing a N-terminal AH. Since AHs were only found among chromosomal CorMR homologs in *Nostocales* and *Oscillatoriales*, only sequences from these species are shown. The N-terminal region comprising the AH (roughly 10 aa) is highly conserved among *Nostocales* species and less among *Oscillatoriales* species.

**Fig. S9.**
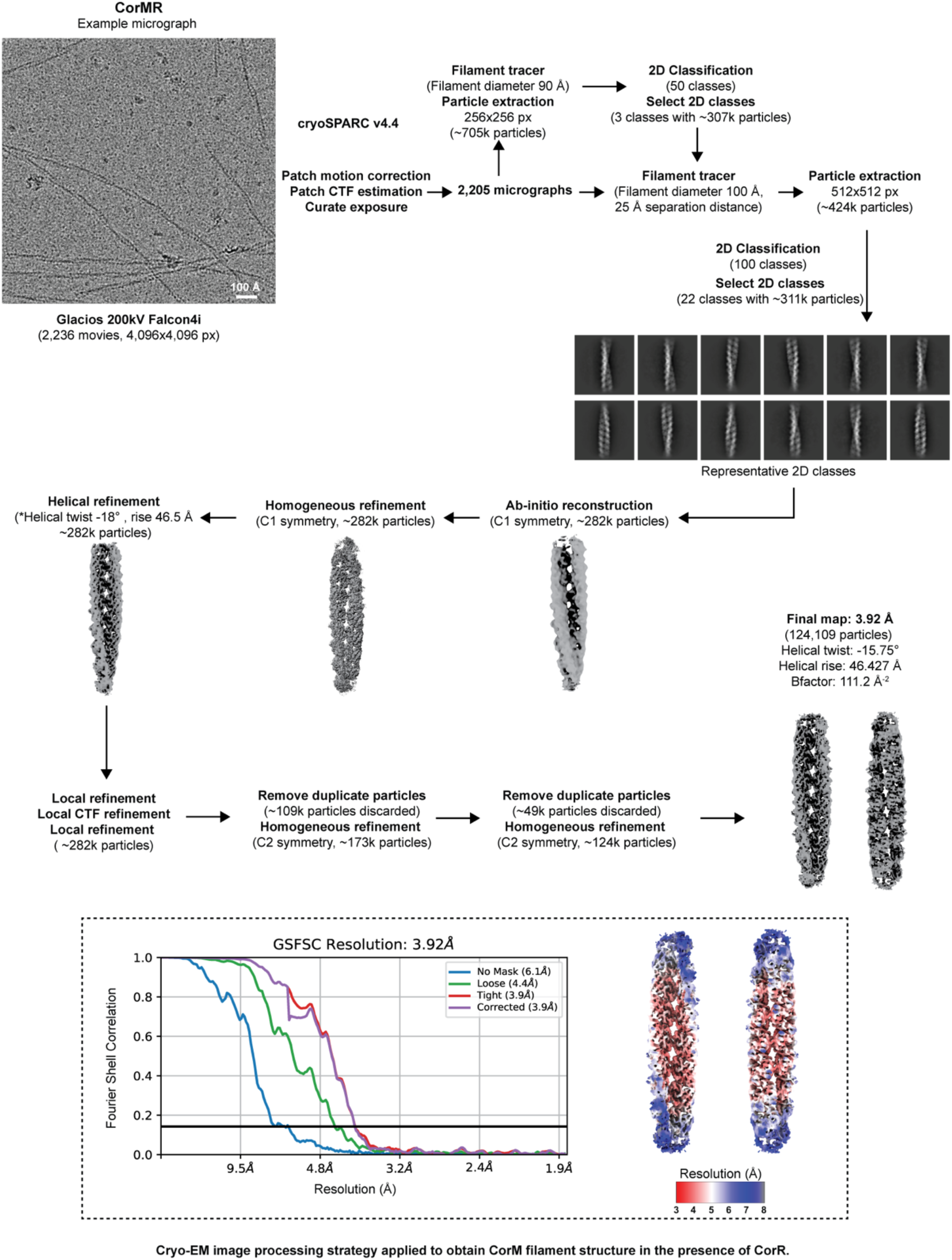
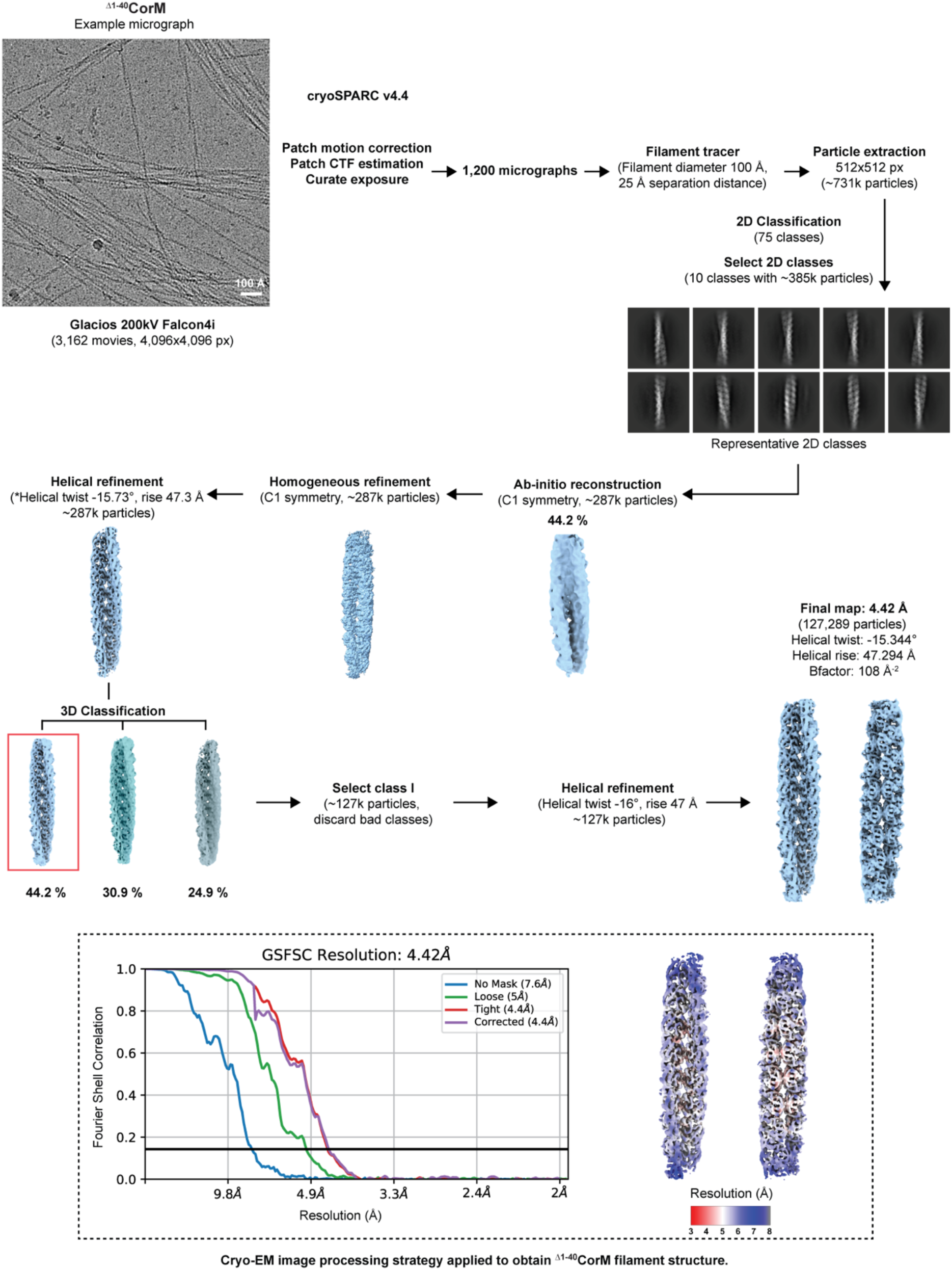
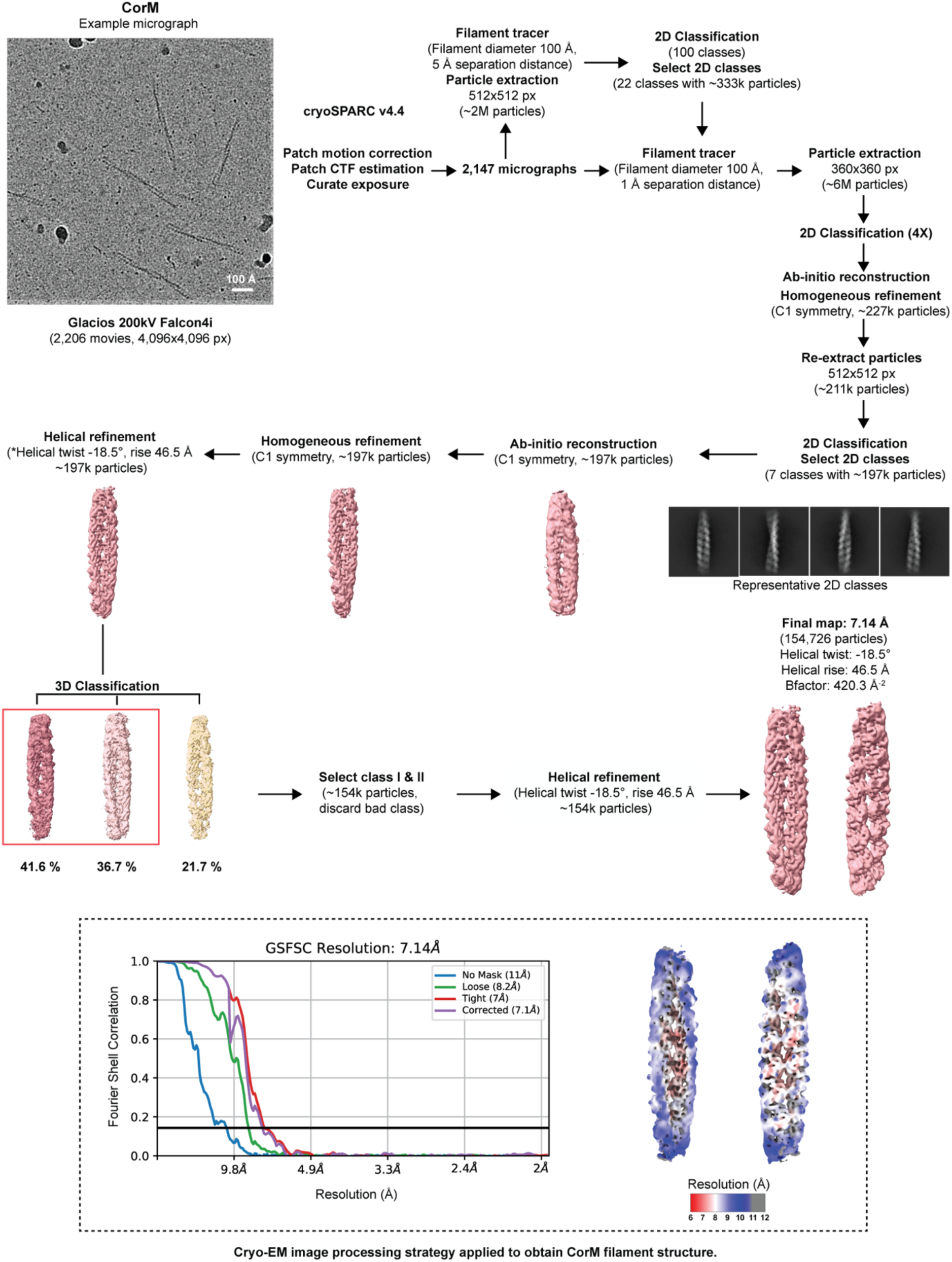
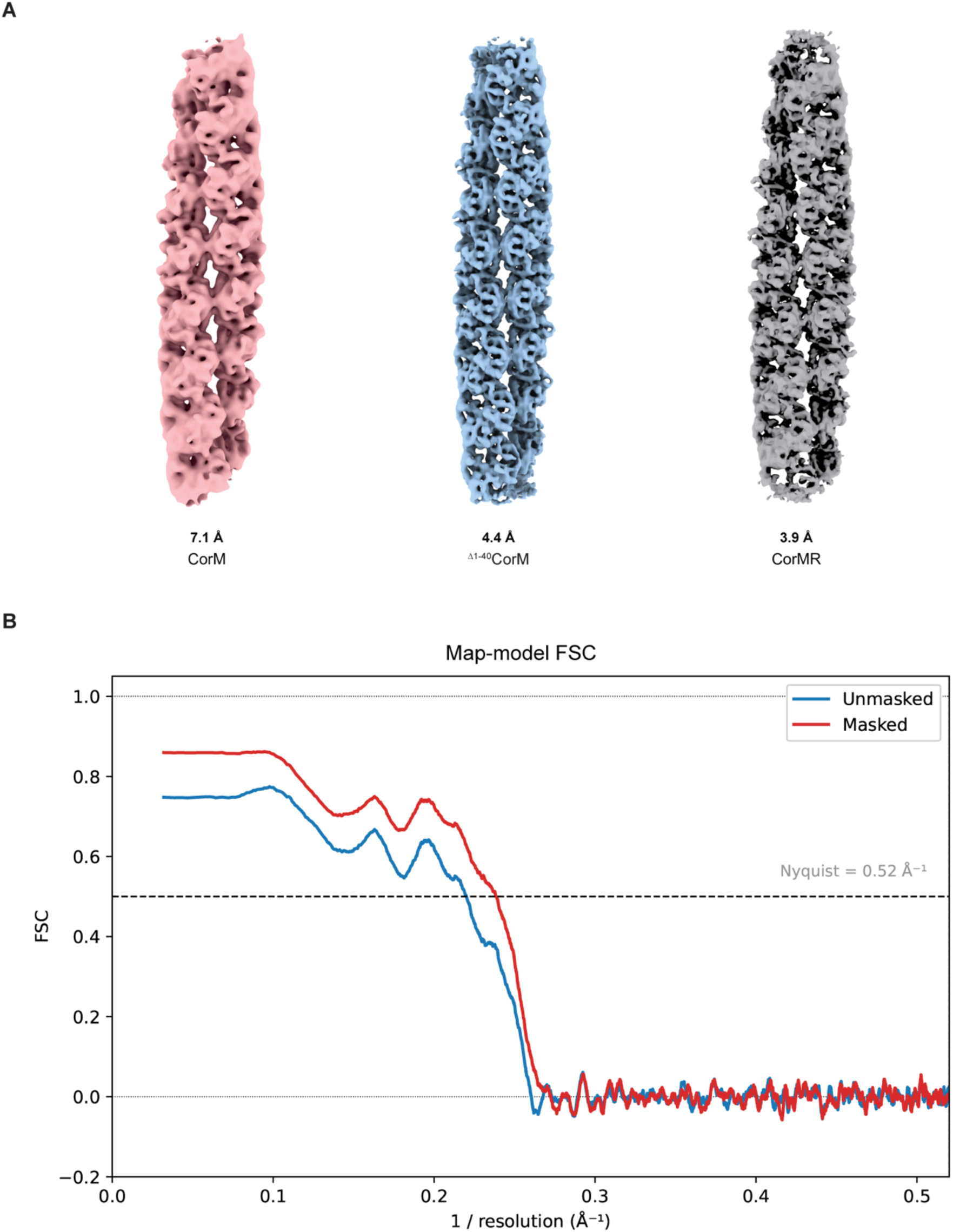
Cryo-EM image processing strategy applied to obtain structure of CorM filament (CorMR filaments (9.1), ^Δ1-40^CorM (9.2), and CorM (9.3)) and the final cryo-EM maps (9.4) Schematics of the image processing strategy used for determining all the three obtained CorM filament structures using cryoSPARC v4.4. Representative micrographs, 2D class averages, 3D class averages, and final electron microscopy density maps are shown. For helical refinement jobs, the initial estimates of helical rise and helical twist were obtained using online helical indexing tool (HI3D-Helical Indexing using cylindrical projection of a 3D map (*82*)). These values are marked with * under each helical refinement job. Two side views of local resolution-colored maps and Fourier Shell Correlation (FSC) curves are displayed. The surface representation of the final cryo-EM maps of CorM, ^Δ1-40^CorM, and CorMR and their estimated resolutions (using FSC 0.143 criteria) are shown in Fig. S9.4 (**A**). The FSC between the refined CorMR model and its cryo-EM map was calculated using Phenix. The masked (red) and unmasked (blue) curves in Fig. S9.4 (**B**) represent correlations computed with and without a soft solvent mask, respectively. The resolution estimates correspond to the FSC = 0.5 cutoff.

**Fig. S10:**
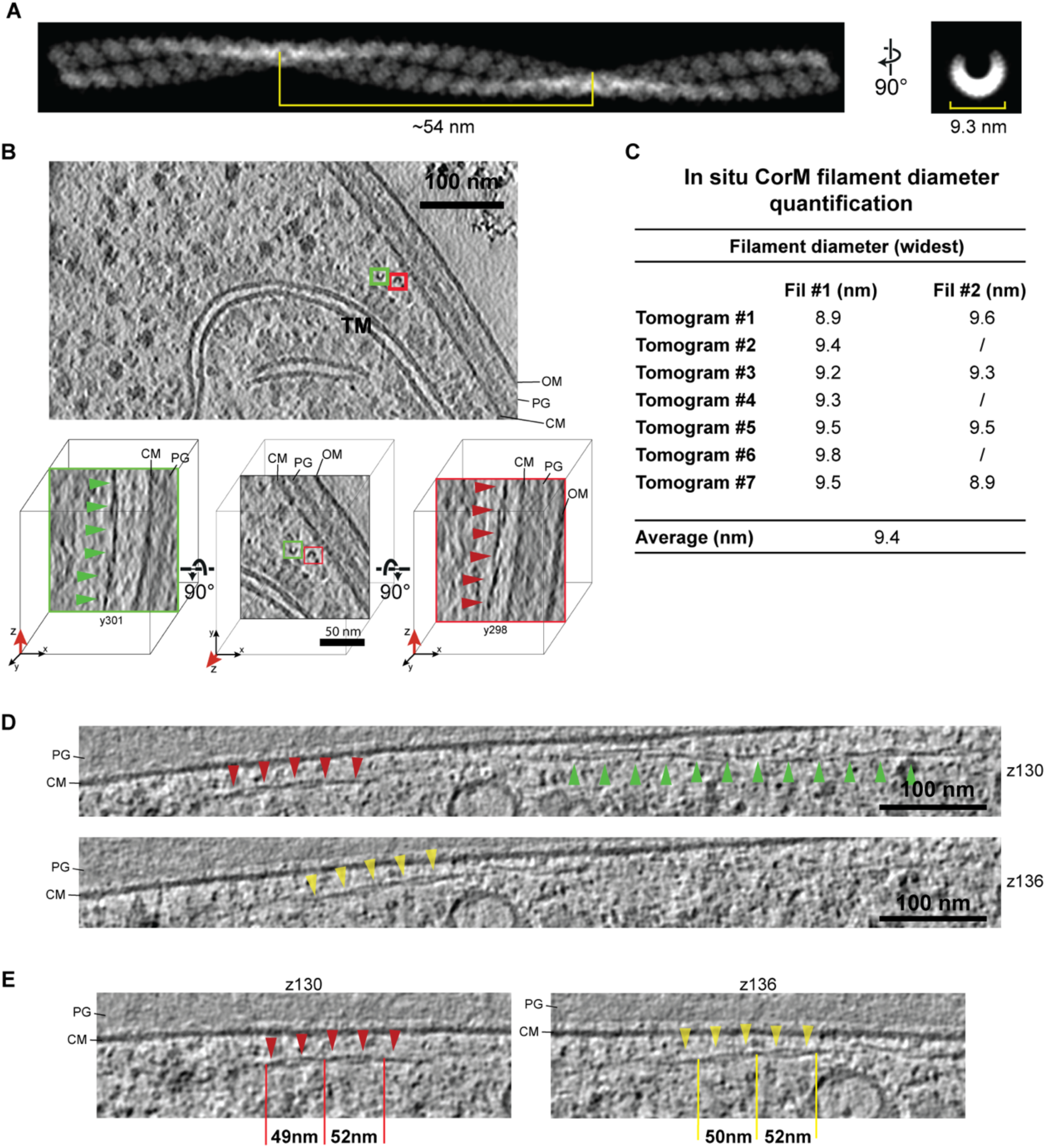
*In situ* cryo-ET of putative CorM filaments (**A)** Maximum intensity projection micrograph of an *in vitro* CorM filament shown from two orientations. This panel serves as a reference to compare the filament characteristics between the *in vitro* filament structure and filaments observed within cells (panels C, E, F). To generate the projection image, the CorM model generated in this study was expanded along the filament direction to allow better appreciation of the filament pitch. From this expanded model a 10Å density map was generated using the molmap command in UCSF ChimeraX (*78*). The density map was then opened in 3dmod (*91*) and displayed summing 100 slices in the XY and YZ direction. The filament dimension measurements were performed on the filament model using the distance measurement tool in UCSF ChimeraX. (**B**) Top image: slice through an Isonet-denoised cryo-electron tomogram (11.04 Å thickness) of *Anabaena* WT. The cross-sections of two putative CorM filaments close to the cell membrane are highlighted in green and red boxes. CM - cell membrane; PG - Peptidoglycan layer; OM - outer membrane; TM - thylakoid membrane. Bottom row: Zoom-in views of the two individual filaments, shown again in the XY slice (middle image) and XZ slices (left and right images) of the tomogram. The XZ views allow the visualization of the helical filament shape (annotated with green and red arrowheads, respectively) consisting of two protofilaments. The number below the XZ views denotes the slice in Y-direction that is displayed. (**C**) Diameter measurements from 11 CorM filaments visualized within 7 Isonet-denoised cryo-electron tomograms (all tomograms had a pixel size of 11.04 Å/px). Given the cortical location of filaments and their orientation following the cell shape, filament cross-sections were seen in XY views, and the longitudinal axis of filaments were visible in the XZ views. For each filament one single diameter measurement in the XZ view was performed at the position where filaments displayed the widest diameter using the distance measurement tool in 3dmod. Only filaments which showed helical twist and corresponded to the expected shape and dimensions were included in this quantification. (**D**) Slice through a cryo-electron tomogram of an *Anabaena Δcse* cell filament (42.84 Å thickness) (*53*) displaying a CorM filament in longitudinal orientation. The two panels show XY sections at different z-heights in the tomogram displaying the different segments of the filament, annotated by red, yellow and green arrowheads. The Z-slices of the tomogram at which the different segments are displayed are indicated on the right, respectively. Only one filament was observed in this perpendicular orientation, which can be explained with our sample preparation and milling orientation, where filaments are expected to be observed mostly in cross-section. (**E**) The two panels show enlarged views of the red (left) and yellow (right) segments from (D), where the filament twist is clearly visible. The repeat length (from smallest-to-smallest diameter section) is annotated and the distance measurements (as performed using the 3dmod measurement tool) are shown. The distance measurements are close to the value measured for the distance measured in the *in vitro* filament model (see panel A). Specifics of cryo-ET data of CorM *in situ* filaments is given in Table S3. Scale bars, where applicable, are annotated in the figure.

**Fig. S11.**
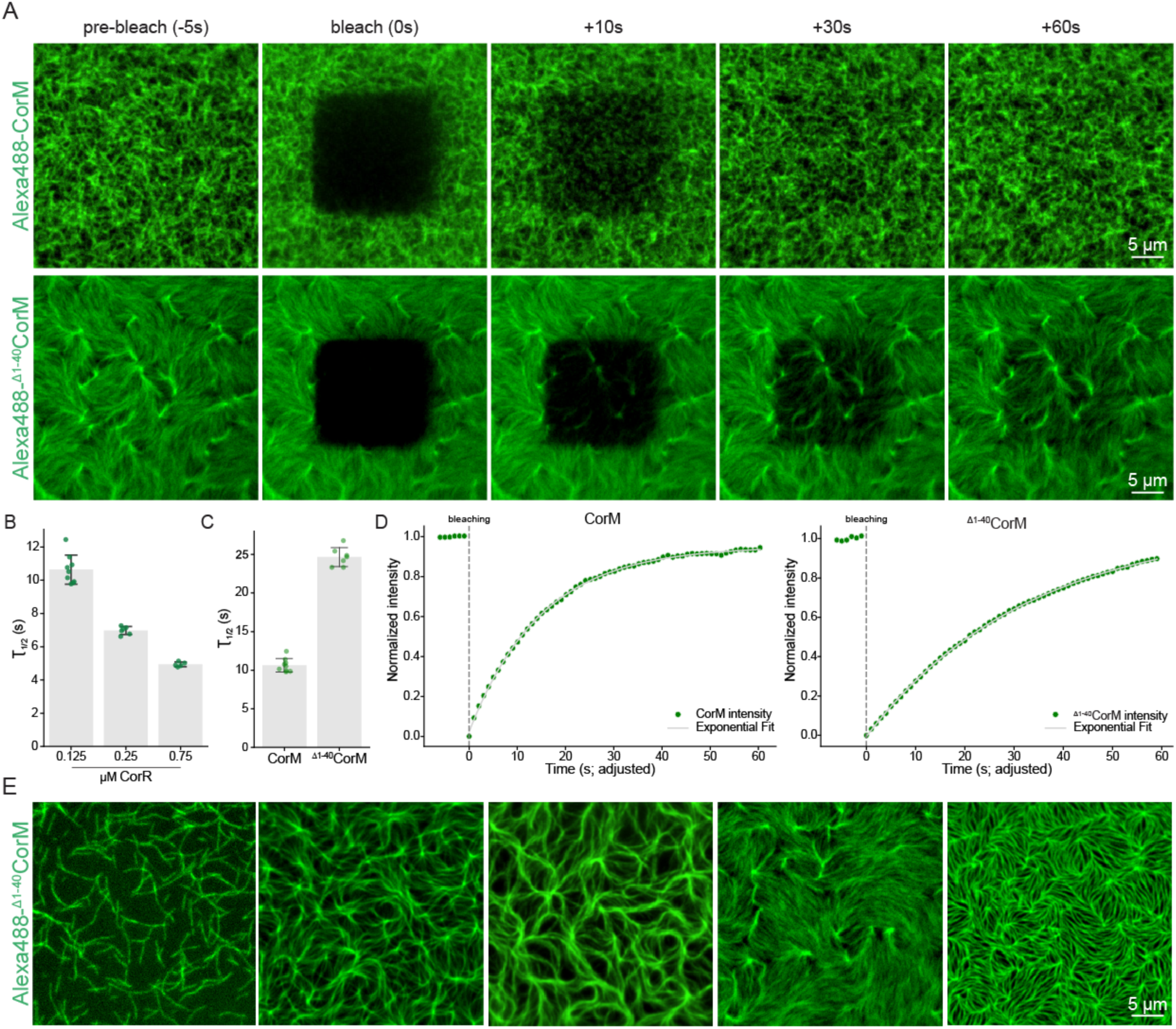
The N-terminal IDR prevents filament bundling and increases protein dynamics (**A**) Representative FRAP snapshots from TIRF microscopy time series (Supplementary Movies 4-5) of pre-formed CorMR or ^Δ1-40^CorMR filaments (0.75 µM unlabeled CorM WT or ^Δ1-40^CorM, 0.25 µM Alexa488-CorM or Alexa488-^Δ1-40^CorM and 0.125 µM unlabeled CorR) on SLBs subjected to photobleaching. (**B**) Dependency of the half-recovery time on CorR concentration. Plot showing the half-recovery time (T_1/2_) in seconds as a function of CorR concentration (0.125 µM, 0.25 µM, and 0.75 µM). Increasing concentrations of CorR lead to a dose-dependent decrease in T_1/2_, suggesting higher CorM dynamics with higher CorR levels. Error bars represent the mean ± SD from individual measurements: 0.125 µM: 10.64 ± 0.87 nm/s (n=9); 0.25 µM: 6.97 ± 0.24 nm/s (n=6); 0.75 µM: 4.94 ± 0.14 nm/s (n=5). Half-recovery times comparing CorM WT and ^Δ1-40^CorM in the presence of 0.125 µM CorR, showing that the N-terminal IDR in CorM modulates dynamics and reaction kinetics. Error bars represent the mean ± SD from individual measurements: CorM values are the same as in (B); ^Δ1-40^CorM: 24.65 ± 1.22 nm/s (n=7). (**D**) Representative recovery curves from FRAP experiments comparing CorM WT and ^Δ1-40^CorM in the presence of 0.125 µM CorR. (**E**) Representative snapshots of ^Δ1-40^CorM (protein concentrations same as in A) filaments in the course of an imaging session. Like CorM WT, ^Δ1-40^CorM initially forms individual filaments. These filaments then form thick bundles but eventually assemble into a well-defined network of filaments, which appears much more ordered compared to CorM WT (see Fig. 5B for comparison).

**Fig. S12.**
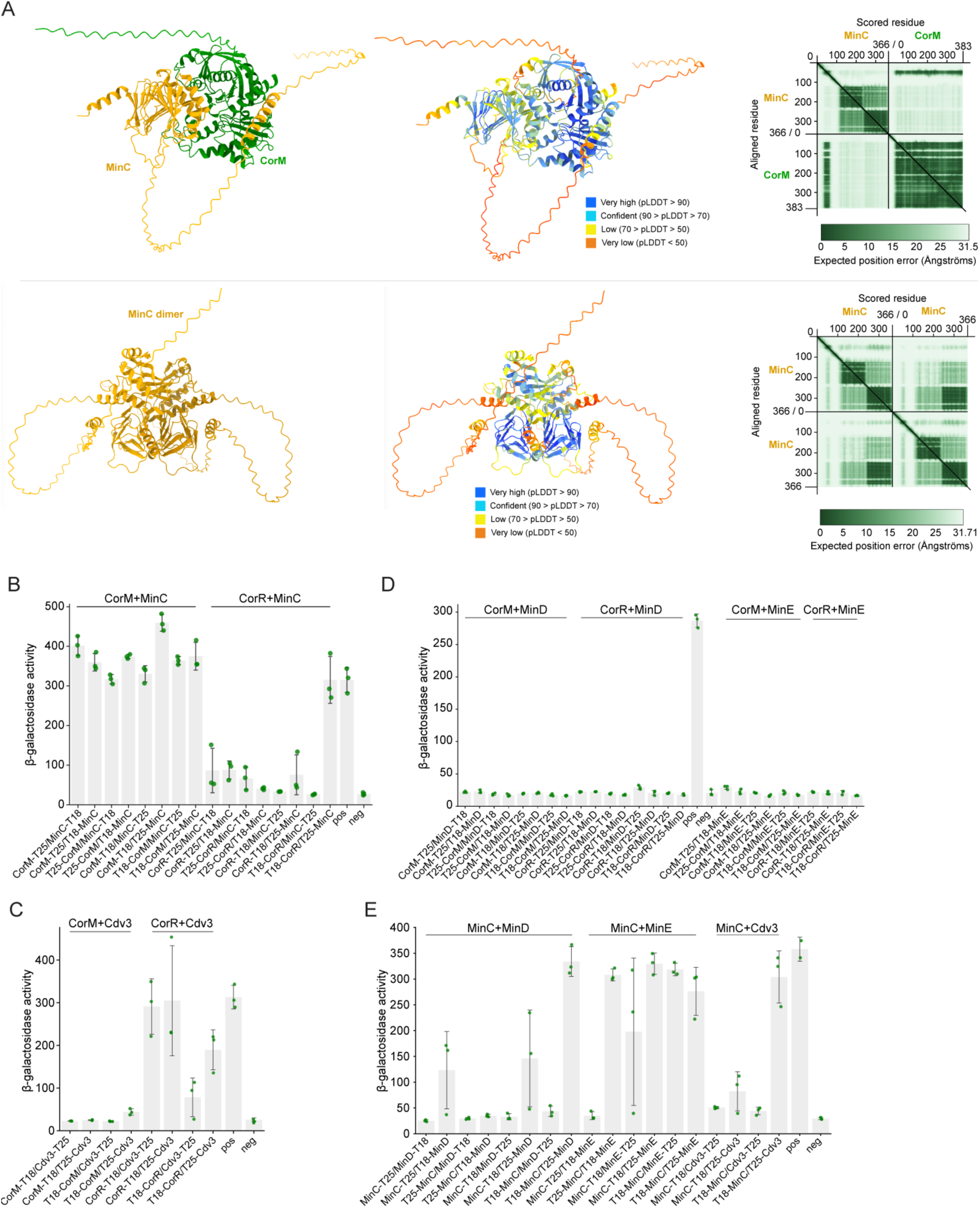
Properties of the *Anabaena* Min system and it’s modulating function on CorM (A) (top) AlphaFold3-predicted interaction between CorM (green) and MinC (orange), together with the corresponding PAE map, showing interaction of MinC’s N-terminal α-helix with CorM, as well as a plDDT score-colored CorM-MinC complex (pTM score: 0.51; ipTM score: 0.76). (bottom) AlphaFold3-predicted MinC (orange) dimer (pTM score: 0.46; ipTM score: 0.47), including the corresponding PAE map and a plDDT score-colored dimer. (**B-D**) B2H assays testing for the interaction of CorMR with (**B**) MinC, (**C**) Cdv3 or (**D**) MinD and MinE fused to either the T25 or T18 subunit. Both, CorM and CorR interact with MinC but only CorR interacts with Cdv3. Neither CorM nor CorR interacts with MinD or MinE. Negative control: (B, D) CorM-T25 with empty pUT18C plasmid or (C) CorM-T18 with empty pKNT25 plasmid. Positive control: Zip/Zip control Error bars represent the mean ± SD of three independent replicates (n=3). (**E**) B2H assays testing for the interaction of MinC with MinD, MinE and Cdv3 fused to either the T25 or T18 subunit. Results show that MinC interacts with all three proteins. Negative control: MinC-T25 with empty pUT18C plasmid. Positive control: Zip/Zip control. Error bars represent the mean ± SD of three independent replicates (n=3).

**Fig. S13.**
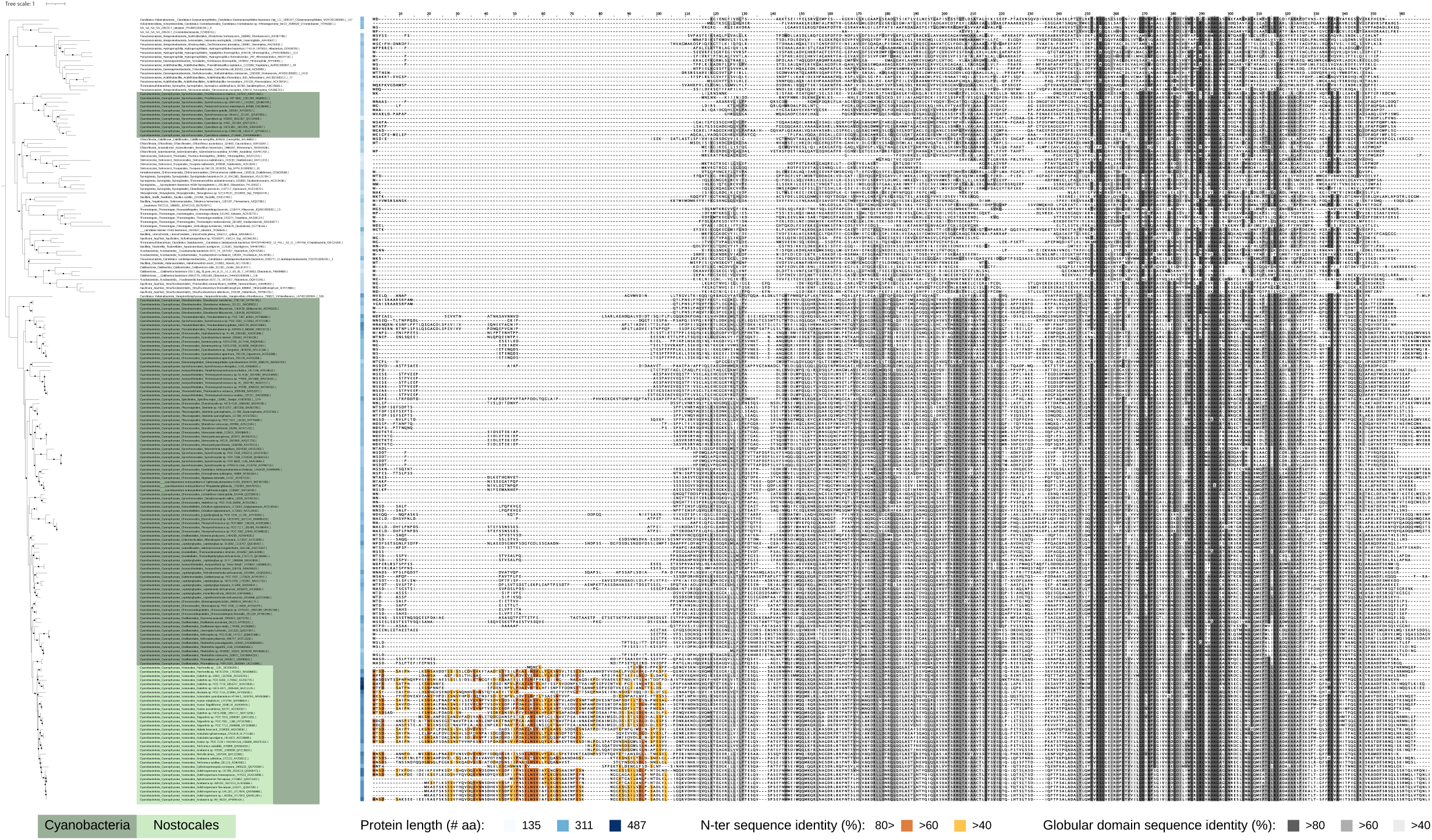
In *Nostocales* cyanobacteria MinC exhibits an extended N-terminal domain MinC phylogenetic tree, including cyanobacteria and other bacterial species. Blue-shaded cubes next to the tree indicates MinC sequence lengths. Note that specifically *Nostocales* cyanobacteria contain long MinC sequences (>300 aa), which is represented by an N-terminal IDR with an a-helix in between. On the right, a multiple sequence alignment of all listed MinCs is shown with two different conservation patterns. Orange shades indicate conservation exclusively among the N-terminal IDR of *Nostocales* MinCs whereas grey shades indicate conservation exclusively among the globular domain of MinC. Black dots indicate Ultrafast Bootstrap supports (UFB) >= 85. The scale bar represents the average number of substitutions per site.

**Fig. S14.**
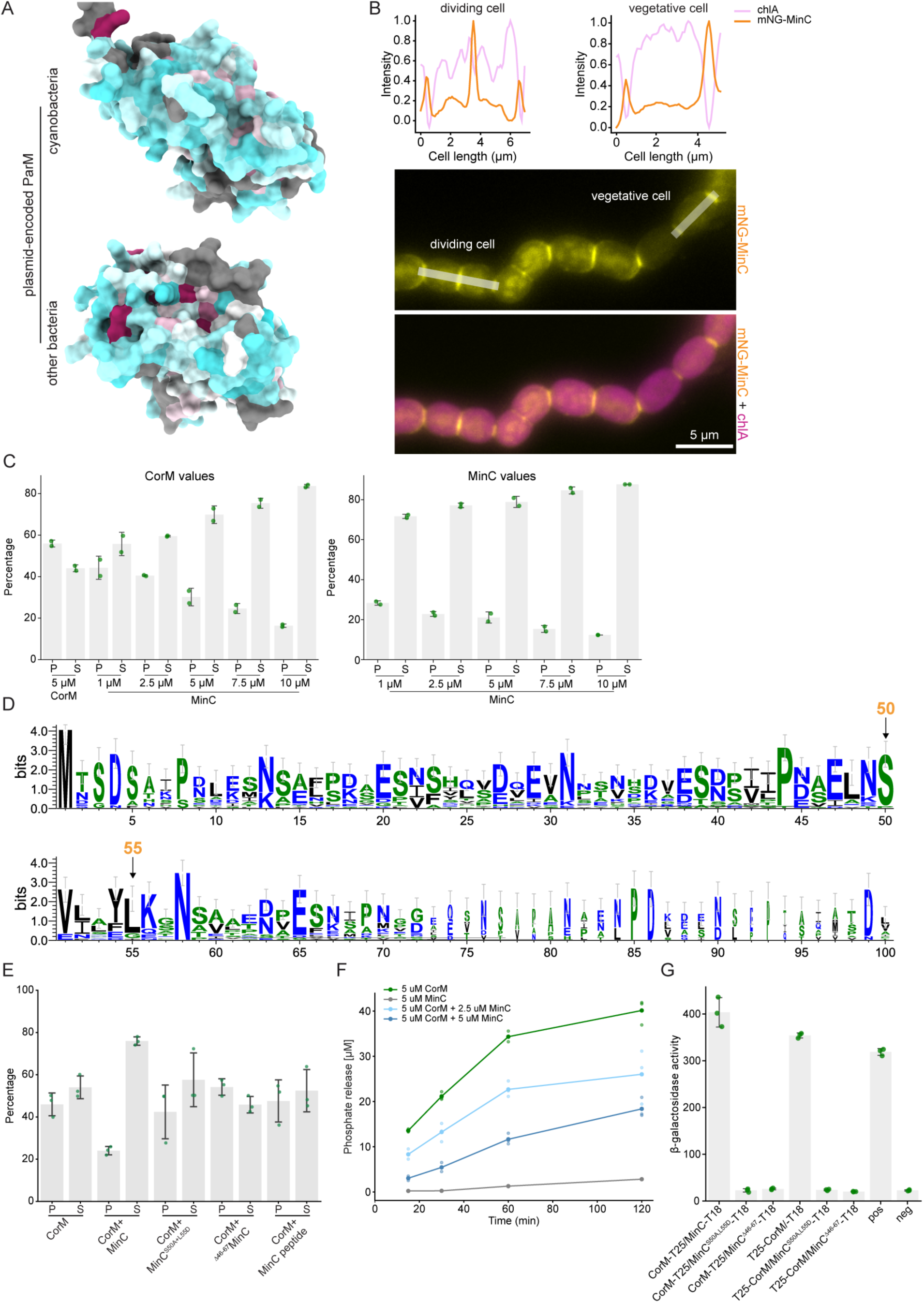
MinC’s N-terminal a-helix mediates CorM depolymerization (**A**) AlphaFold3-predicted structures of (top) plasmid-encoded ParM (#BAB78165.1; All7081) from the large low copy number Alpha plasmid from *Anabaena* or (bottom) plasmid-encoded EcParM (WP_000959884.1) from the R1 plasmid. Conservation is based on multiple sequence alignments from all identified cyanobacterial plasmid-encoded ParMs or all identified other bacterial plasmid-encoded ParMs. Coloring by sequence conservation is based on the entropy-based measure AL2CO from ChimeraX (*78*) using the default coloring scheme (cyan: poor conservation; white: intermediate conservation; red: highly conserved). (**B**) mNG-MinC fluorescence, chlA autofluorescence and merged micrographs of *Anabaena* expressing mNG-MinC from P_ntcA_ integrated into the neutral chromosomal *thrS2* sites. Fluorescence intensity profiles show high levels of MinC at the cell poles and midcell in diving cells and only at the cell poles in vegetative cells. (**C**) Quantifications from pelleting assay shown in Fig. 6F showing values for CorM and MinC separately. Different MinC protein concentrations are indicated below. Error bars represent the mean ± SD of two independent replicates (n=2). (**D**) Sequence logo showing the conservation of aa residues within the N-terminal a-helix of *Nostocales* cyanobacteria. A multiple sequence alignment of all tested MinC sequences from cyanobacteria and other bacteria is shown in Fig. S13. (**E**) Quantifications from pelleting assay shown in Fig. 6G showing values for CorM. Both MinC helix mutants, MinC^S50A+L55D^ and ^Δ46-67^MinC lost the ability to depolymerize CorM filaments. A synthesized peptide mimicking MinC’s N-terminal a-helix had no influence on CorM polymerization, indicating that other aspects of the interaction of CorM with MinC are essential for active depolymerization. Error bars represent the mean ± SD of three independent replicates (n=3). (**F**) CorM ATPase activity assay with or without indicated MinC concentrations resolved over time, showing that MinC acts on CorM by decreasing its enzymatic activity. Data represents the mean values for each data point from three independent replicates (n=3). (**G**) B2H assays testing for the interaction of CorM with MinC helix mutants (MinC^S50A+L55D^ and ^Δ46-67^MinC) fused to either the T25 or T18 subunit. Results show that CorM interaction with MinC is lost in both MinC helix mutants. Negative control: CorM-T25 with empty pUT18C plasmid. Positive control: Zip/Zip control. Error bars represent the mean ± SD of three independent replicates (n=3).

---

**Table S1.** Strains and plasmids.

| Strain | Details | Reference |
| --- | --- | --- |
| DH5α | *fhuA2 lac(del)U169 phoA glnV44 Φ80' lacZ(del)M15 gyrA96 recA1 relA1 endA1 thi-1 hsdR17* | LSF |
| BL21 (DE3) | *fhuA2 [lon] ompT gal ($\lambda$ DE3) [dcm] ΔhsdS $\lambda$ DE3 = $\lambda$ sBamHlo ΔEcoRI-B int::(*lacI*::PlacUV5::T7 gene1) i21 Δnin5* | LSF |
| OverExpress<sup>TM</sup> C41 (DE3) | *F- ompT hsdSB (rB- mB-) gal dcm (DE3)* | Sigma Aldrich |
| BTH101 | *F- cya-99 araD139 galE15 galK16 rpsL1 (Str<sup>R</sup>) hsdR2 mcrA1 mcrB1* | Euromedex (A gift from Tal Dagan; Kiel University) |
| *Anabaena* sp. PCC 7120 | wild type (WT) | A gift from Tal Dagan (Kiel University) |
| BS108 | ΔcorMR; Marker-less deletion of *corMR* open reading frame (*corM*: WP_044522477.1; *corR*: WP_010999216.1) | This study |
| BS191 | *Anabaena*; P<sub>corMR</sub>-*mNG-corM-corR* (in frame translational fusion of *mNG* with *corM*, including an ASAS-linker) | This study |
| BS196 | *Anabaena*; P<sub>corMR</sub>-*corM-corR-mNG* (in frame translational fusion of *corR* with *mNG*, including an ASASA-linker) | This study |
| BS326 | *Anabaena*; *thrS2::P<sub>ntcA</sub>-mNG-minC* | This study |
| BS396 | *Anabaena*; P<sub>corMR</sub>-*mNG-Δ<sup>1-40</sup>corM-corR* (in frame translational fusion of *mNG* with Δ<sup>1-40</sup>*corM*, including an ASAS-linker) | This study |
| Plasmid | Details | Reference |
| pRL25C | Shuttle plasmid for *E. coli* and cyanobacteria, Km<sup>R</sup> | (112) |
| pRL623 | Methylation plasmid/helper plasmid, Cm<sup>R</sup> | (112) |
| pRL443 | Conjugational plasmid to mobilize plasmids into *Anabaena*, Amp<sup>R</sup> | (112) |
| pCSL135 | P<sub>ntcA</sub>-SepF |  |
| pKNT25 | P<sub>lac</sub>-T25, Km<sup>R</sup> | Euromedex |
| pKT25 | P<sub>lac</sub>-T25, Km<sup>R</sup> | Euromedex |
| pUT18 | P<sub>lac</sub>-T18, Amp<sup>R</sup> | Euromedex |
| pUT18C | P<sub>lac</sub>-T18, Amp<sup>R</sup> | Euromedex |
| pKT25-*zip* | pKT25; P<sub>lac</sub>-T25-*zip*; Km<sup>R</sup> | Euromedex |
| pUT18C-*zip* | pUT18C; P<sub>lac</sub>-T18-*zip*, Amp<sup>R</sup> | Euromedex |
| pAM5411 | RSF1010-based broad host-range plasmid with RK2 bom site and pUC origin expressing YFP, Sp<sup>R</sup>/Sm<sup>R</sup> | (44) |
| pCpf1b | Cpf1-based CRISPR plasmid with *sacB* counter selection, Km<sup>R</sup> | (46) |
| pTB146 | P<sub>T7</sub>-His6-SUMO-X; *E. coli* expression vector for translational fusions of His6-SUMO with your gene of interest X, Amp<sup>R</sup> | (113) |
| pBAD33.1-Km | pBAD33.1 vector with Cm<sup>R</sup> replaced by Km<sup>R</sup> | This study |
| pCSAV266 | pKNT25; P<sub>lac</sub>-*minC*-T25, (MinC: #WP_190449770.1), Km<sup>R</sup> | A gift from Antonia Herrero (Sevilla University) |
| pCSAV268 | pKT25; P<sub>lac</sub>-T25-*minC*, Km<sup>R</sup> |  |
| pCSAV269 | pUT18; P<sub>lac</sub>-*minC*-T18, Amp<sup>R</sup> |  |
| pCSAV267 | pUT18C; P<sub>lac</sub>-T18-*minC*, Amp<sup>R</sup> |  |
| pBS1 | pKNT25; P<sub>lac</sub>-*corM*-T25, (CorM: #WP_044522477.1), Km<sup>R</sup> | This study |
| pBS2 | pKT25; P<sub>lac</sub>-T25-*corM*, Km<sup>R</sup> | This study |
| pBS3 | pUT18; P<sub>lac</sub>-*corM*-T18, Amp<sup>R</sup> | This study |
| pBS4 | pUT18C; P<sub>lac</sub>-T18-*corM*, Amp<sup>R</sup> | This study |
| pBS9 | pKNT25; P<sub>lac</sub>-*corR*-T25, (CorR: #WP_010999216.1), Km<sup>R</sup> | This study |
| pBS10 | pKT25; P<sub>lac</sub>-T25-*corR*, Km<sup>R</sup> | This study |
| pBS11 | pUT18; P<sub>lac</sub>-*corR*-T18, Amp<sup>R</sup> | This study |
| pBS12 | pUT18C; P<sub>lac</sub>-T18-*corR*, Amp<sup>R</sup> | This study |
| pBS37 | pTB146; P<sub>T7</sub>-His6-SUMO-*corM*, Amp<sup>R</sup> | This study |
| pBS41 | pTB146; P<sub>T7</sub>-His6-SUMO-*corR*, Amp<sup>R</sup> | This study |
| pBS47 | pTB146; P<sub>T7</sub>-His6-SUMO-*parR* (ParR from *E. coli* R1 plasmid - #P11906.1), Amp<sup>R</sup> | This study |
| pBS71 | pTB146; P<sub>T7</sub>-His6-SUMO-3xGly-*corM*, Amp<sup>R</sup> | This study |
| pBS75 | pTB146; P<sub>T7</sub>-His6-SUMO-3xGly-*parM* (ParM from *E. coli* R1 plasmid - #WP_000959884.1), Amp<sup>R</sup> | This study |
| pBS80 | pTB146; P<sub>T7</sub>-*mNG*-*corM*, Amp<sup>R</sup> | This study |
| pBS81 | pTB146; P<sub>T7</sub>-*corM*-*mNG*, Amp<sup>R</sup> | This study |
| pBS103 | pCpf1b with spacer targeting 5' region of *corM* | This study |
| pBS105 | pCpf1b with spacer targeting *corMR* | This study |
| pBS108 | pBS105 with ~1000 bp homologous flanks for Δ*corMR* | This study |
| pBS112 | pRL25C-P<sub>ntcA</sub>-*mNG*-*corM* | This study |
| pBS113 | pRL25C-P<sub>ntcA</sub>-*corM*-*mNG* | This study |
| pBS134 | pTB146; P<sub>T7</sub>-*mNG*-*parM* (ParM from *E. coli* R1 plasmid), Amp<sup>R</sup> | This study |
| pBS135 | pTB146; P<sub>T7</sub>-*parM*-*mNG* (ParM from *E. coli* R1 plasmid), Amp<sup>R</sup> | This study |
| pBS136 | pCpf1b with spacer targeting *thrS2* neutral site, Km<sup>R</sup> | This study |
| pBS138 | pBS136 with ~1000 bp homologous flanks, sandwiching a BamHI site for cloning purposes, for integration into *thrS2* neutral site, Km<sup>R</sup> | This study |
| pBS140 | pBS138; *thrS2::P<sub>ntcA</sub>-mNG-corM* (in frame translational fusion of *mNG*-*corM*, including an ASAS-linker); plasmid for double homologous recombination to insert *P<sub>ntcA</sub>-mNG-corM* into the *thrS2* neutral site using ~1000 bp homologous flanks, Km<sup>R</sup> | This study |
| pBS142 | pBS138; *thrS2::P<sub>ntcA</sub>-corM-mNG* (in frame translational fusion of *corM-mNG*, including an ASAS-linker); plasmid for double homologous recombination to insert *P<sub>ntcA</sub>-corM-mNG* into the *thrS2* neutral site using ~1000 bp homologous flanks, Km<sup>R</sup> | This study |
| pBS154 | pCpf1b with spacer targeting 3' region of *corR*, Km<sup>R</sup> | This study |
| pBS159 | pBAD33.1-Km with P<sub>BAD</sub>-*corR-mScarlet-I*, Km<sup>R</sup> | This study |
| pBS166 | pBS138; *thrS2::P<sub>ntcA</sub>-corR-SCFP3* (in frame translational fusion of *corR-SCFP3*, including an ASASA-linker); plasmid for double homologous recombination to insert *P<sub>ntcA</sub>-corR-SCFP3* into the *thrS2* neutral site using ~1000 bp homologous flanks, Km<sup>R</sup> | This study |
| pBS184 | pBS138; *thrS2::P<sub>ntcA</sub>-corR-mNG* (in frame translational fusion of *corR-mNG*, including an ASASA-linker); plasmid for double homologous recombination to insert *P<sub>ntcA</sub>-corR-mNG* into the *thrS2* neutral site using ~1000 bp homologous flanks, Km<sup>R</sup> | This study |
| pBS190 | pCpf1b with spacer targeting 5' region of *corM*, Km<sup>R</sup> | This study |
| pBS191 | pBS190; plasmid for double homologous recombination for in frame translational fusion of *mNG-corM*, including an ASAS-linker and ~1000 bp homologous flanks, Km<sup>R</sup> | This study |
| pBS192 | pBAD33.1-Km with P<sub>BAD</sub>-$\Delta 1-10$*corR-mScarlet-I*, Km<sup>R</sup> | This study |
| pBS196 | pBS154; plasmid for double homologous recombination for in frame translational fusion of *corR-mNG*, including an ASASA-linker and ~1000 bp homologous flanks, Km<sup>R</sup> | This study |
| pBS201 | pKNT25; P<sub>lac</sub>-*minD*-T25, Km<sup>R</sup> (MinD: #WP_010997606.1) | This study |
| pBS202 | pKT25; P<sub>lac</sub>-T25-*minD*, Km<sup>R</sup> | This study |
| pBS203 | pUT18; P<sub>lac</sub>-*minD*-T18, Amp<sup>R</sup> | This study |
| pBS204 | pUT18C; P<sub>lac</sub>-T18-*minD*, Amp<sup>R</sup> | This study |
| pBS205 | pKNT25; P<sub>lac</sub>-*minE*-T25, Km<sup>R</sup> (MinE: #WP_010997607.1) | This study |
| pBS206 | pKT25; P<sub>lac</sub>-T25-*minE*, Km<sup>R</sup> | This study |
| pBS208 | pUT18C; P<sub>lac</sub>-T18-*minE*, Amp<sup>R</sup> | This study |
| pBS213 | pKNT25; P<sub>lac</sub>-*cdv3*-T25, Km<sup>R</sup> (Cdv3: #BAB76400.1) | This study |
| pBS214 | pUT18; P<sub>lac</sub>-T25-*cdv3*, Km<sup>R</sup> | This study |
| pBS229 | pTB146; P<sub>T7</sub>-*mNG*-$\Delta 1-40$*corM*, Amp<sup>R</sup> | This study |
| pBS230 | pTB146; P<sub>T7</sub>-$\Delta 1-40$*corM-mNG*, Amp<sup>R</sup> | This study |
| pBS231 | pTB146; P<sub>T7</sub>-His6-SUMO-$\Delta 1-10$*corR*, Amp<sup>R</sup> | This study |
| pBS233 | pTB146; P<sub>T7</sub>-His6-SUMO-3xGly-$\Delta 1-40$*corM*, Amp<sup>R</sup> | This study |
| pBS313 | pBS138; *thrS2::P<sub>ntcA</sub>-corM-mNG<sup>SW</sup>* (in frame translational sandwich fusion of *corM* with *mNG*, including a 5' SGSS and a 3' ASAS-linker between *corM* and *mNG*); plasmid for double homologous recombination to insert *P<sub>ntcA</sub>-corM-mNG<sup>SW</sup>* into the | This study |
|  | *thrS2* neutral site using ~1000 bp homologous flanks, Km<sup>R</sup> |  |
| pBS316 | pTB146; P<sub>T7</sub>-His6-SUMO-4xGly-*minC*, Amp<sup>R</sup> | This study |
| pBS320 | pCpf1b with spacer targeting 5' region of *corR*, Km<sup>R</sup> | This study |
| pBS321 | pBS320 with ~1000 bp homologous flanks for $\Delta^{1-10}$*corR* | This study |
| pBS324 | pBS190 with ~1000 bp homologous flanks for $\Delta^{1-40}$*corM* | This study |
| pBS326 | pBS138; *thrS2::P<sub>ntcA</sub>-mNG-minC* (in frame translational fusion of *mNG-minC*, including an ASASGS-linker); plasmid for double homologous recombination to insert *P<sub>ntcA</sub>-mNG-minC* into the *thrS2* neutral site using ~1000 bp homologous flanks, Km<sup>R</sup> | This study |
| pBS338 | pKT25; P<sub>lac</sub>-T25-$\Delta^{150-162}$*corR*, Km<sup>R</sup> | This study |
| pBS339 | pUT18C; P<sub>lac</sub>-T18-$\Delta^{150-162}$*corR*, Amp<sup>R</sup> | This study |
| pBS346 | pTB146; P<sub>T7</sub>-His6-SUMO-$\Delta^{150-162}$*corR*, Amp<sup>R</sup> | This study |
| pBS348 | pBS320 with ~1000 bp homologous flanks for Δ*corR* | This study |
| pBS353 | pTB146; P<sub>T7</sub>-His6-SUMO-4xGly-*minC*<sup>S50A,L55D</sup>, Amp<sup>R</sup> | This study |
| pBS354 | pTB146; P<sub>T7</sub>-His6-SUMO-4xGly-$\Delta^{46-67}$*minC* (deletion of N-terminal helix), Amp<sup>R</sup> | This study |
| pBS356 | pBS154 with ~1000 bp homologous flanks for *corR*$\Delta^{15-162}$ | This study |
| pBS386 | pUT18; P<sub>lac</sub>-*minC*<sup>S50A,L55D</sup>-T18, Amp<sup>R</sup> | This study |
| pBS387 | pUT18; P<sub>lac</sub>-$\Delta^{46-67}$*minC*-T18, Amp<sup>R</sup> | This study |
| pBS396 | pBS103; plasmid for double homologous recombination for in frame translational fusion of *mNG-$\Delta^{1-40}$corM*, including an ASAS-linker and ~1000 bp homologous flanks, Km<sup>R</sup> | This study |
| pBS415 | pCpf1b lacking *sacB-cpf1*, Km<sup>R</sup> | This study |
| pBS416 | pBS415; P<sub>petE</sub>-*corMR*, Km<sup>R</sup> | This study |
| pBS417 | pBS415; P<sub>petE</sub>-*mNG-corMR* (translational fusion of *mNG* with *corM*, including a GSAGSAAGSGEFGS linker) , Km<sup>R</sup> | This study |

LSF: Lab Support Facility at ISTA; mNG: mNeonGreen; Km<sup>R</sup>: kanamycin resistance; Amp<sup>R</sup>: ampicillin resistance; Cm<sup>R</sup>: chloramphenicol resistance; Str<sup>R</sup>: streptomycin resistance

**Table S2.**
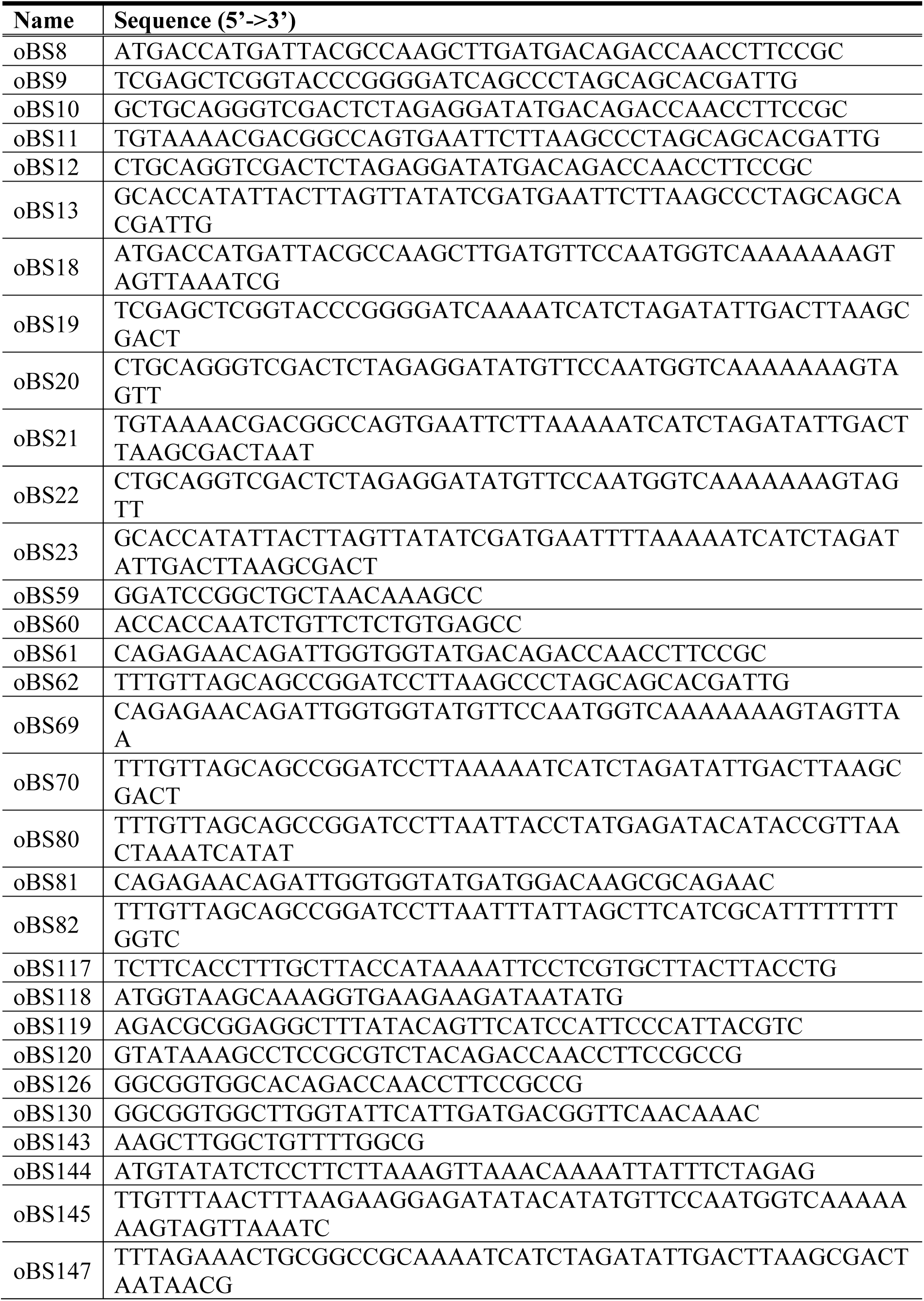
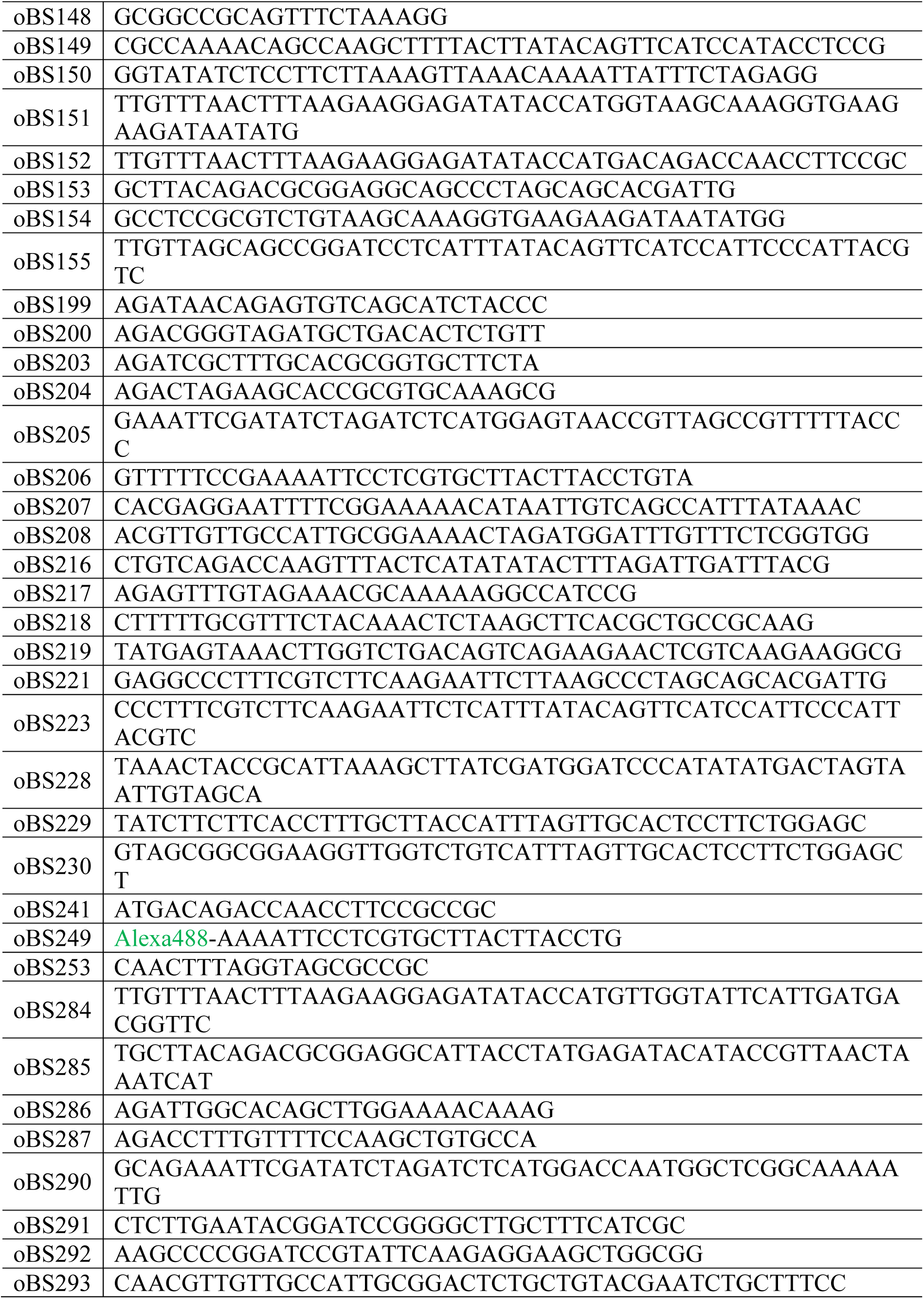
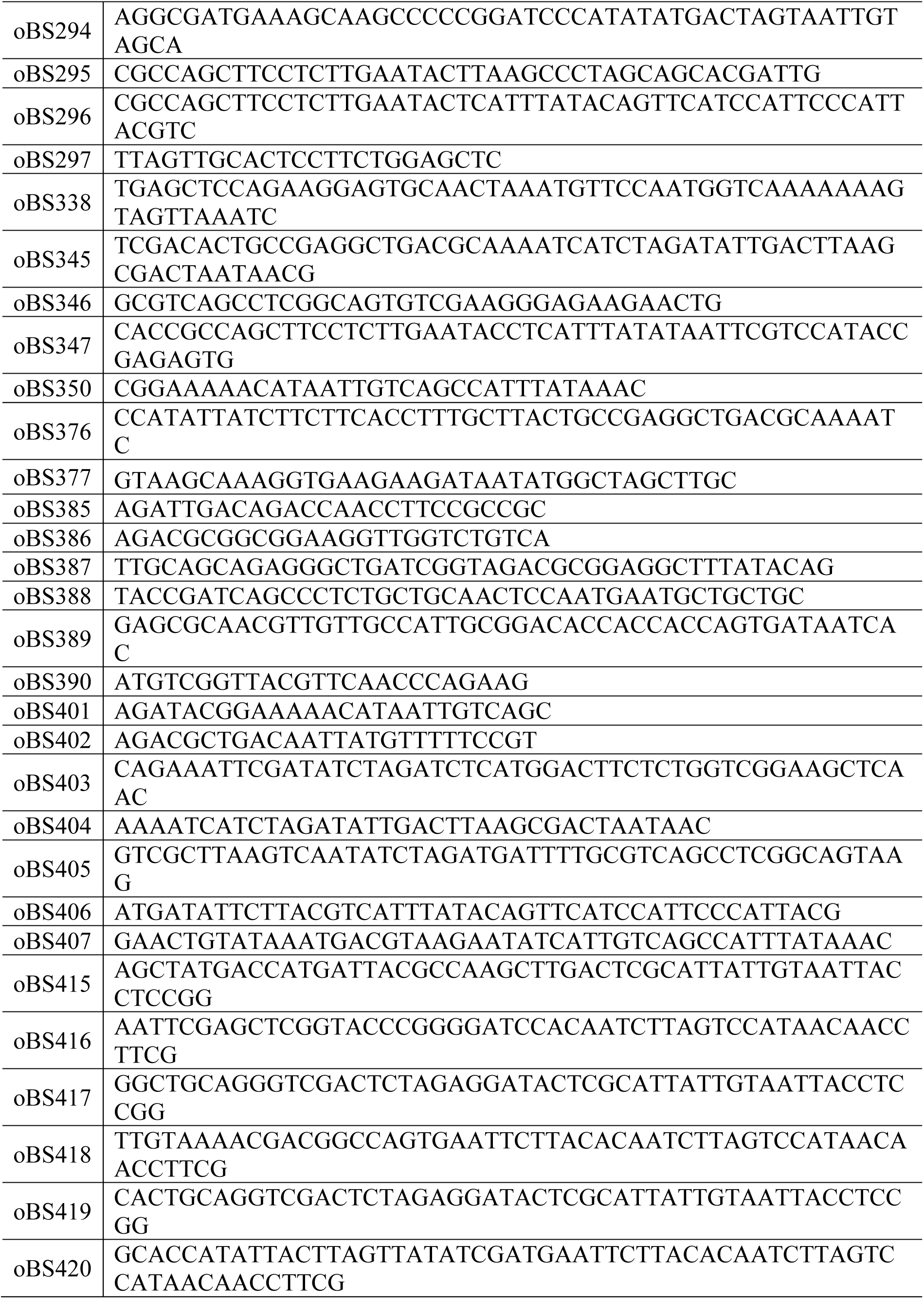
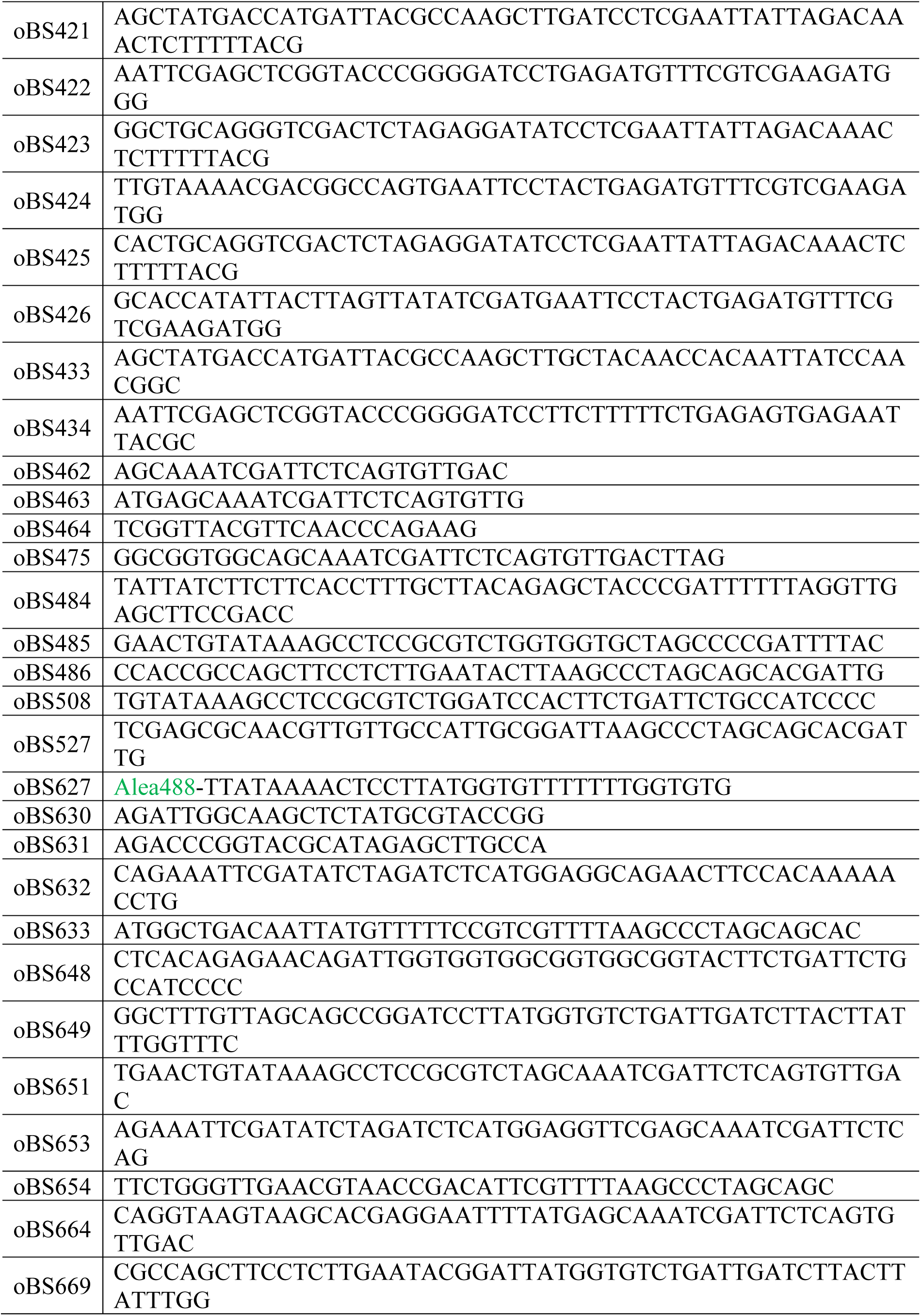
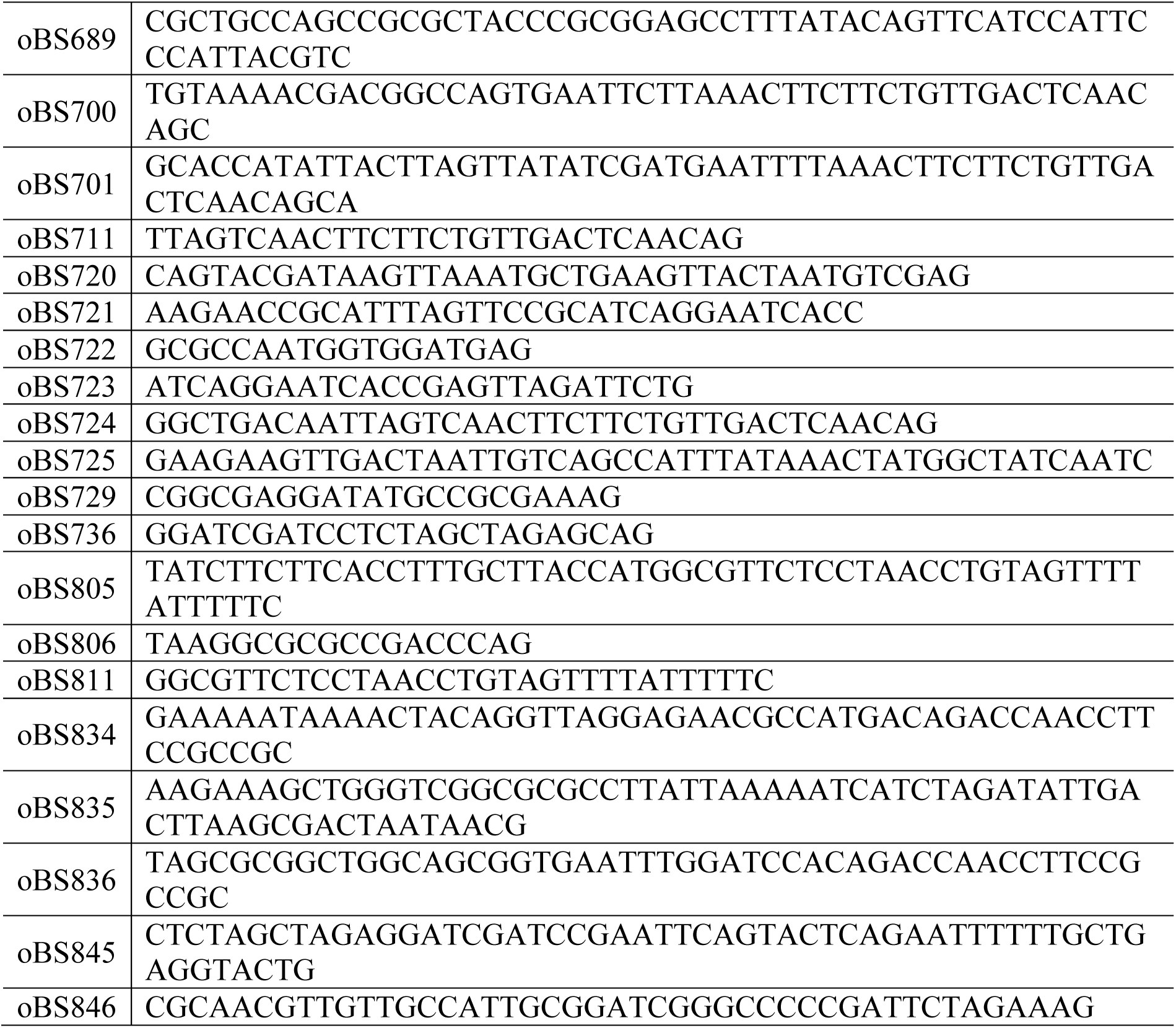
Oligonucleotides.

**Table S3.** Cryo-ET data of CorM *in situ* filaments.

| | *Anabaena* WT | *Anabaena* $\Delta cse$ |
| --- | --- | --- |
| Microscope | FEI Titan Krios 3Gi | FEI Titan Krios 3Gi |
| Voltage [kV] | 300 | 300 |
| Detector | Gatan BioQuantum K3 | Gatan BioQuantum K3 |
| Energy-filter | Yes (slit width 20eV) | Yes (slit width 20eV) |
| Nominal magnification | 64,000x | 33,000x |
| Pixel size [$\text{\AA}/\text{pix}$] | 1.38 | 2.68 |
| Defocus range [$\mu\text{m}$] | -1.5 to -3.5 | -5 to -6 |
| Acquisition scheme | Dose-symmetric | Dose-symmetric |
| Tilt range [$^{\circ}$] | -60 to +60 | -50 to +70 |
| Tilt increment [$^{\circ}$] | 3 | 3 |
| Acquisition software | SerialEM 4.1.10, PACE-tomo<br>1.4.4 | SerialEM 4.2.1, PACE-tomo<br>1.9.2 |
| Total dose [electrons/$\text{\AA}^2$] | 160-180 | 140 |
| Number of frames | 8 | 10 |
| Reconstruction software | AreTomo2 (version 1.3.4) | IMOD |
| Filter | IsoNet | Deconvolution |

## Supplementary Movies

**Movie S1. Live cell time lapse microscopy of mNG-CorM**

*Anabaena* cells expressing mNG-CorM were grown on 1.5% agar pads and illuminated with an in-house plant growth light. Images were acquired at 15-minute intervals. Scale bar: 5 µm.

**Movie S2. Dynamic instability of mNG-CorM *in vivo***

Polymerization and sudden shrinkage of mNG-CorM filaments in *Anabaena* cells recorded by TIRF microscopy. Images were acquired at 10-second intervals. Scale bar: 1 µm.

**Movie S3. Dynamic instability of mNG-^Δ1-40^CorM *in vivo***

Polymerization and sudden shrinkage of mNG-^Δ1-40^CorM filaments in *Anabaena* cells recorded by TIRF microscopy. Images were acquired at 5-second intervals. Scale bar: 1 µm.

**Movie S4. *In situ* cryo-ET of putative CorM filaments in WT *Anabaena***

Movie of the Isonet-corrected cryo-electron tomogram displayed in Fig. S10B showing the two CorM filaments in cross-section in XY view and after rotation in XZ view.

**Movie S5: *In situ* cryo-ET of putative CorM filaments in Δ*cse Anabaena***

Movie of the cryo-electron tomogram showing the *in situ* filaments displayed in Fig. S10D-E. To facilitate filament identification in the crowded tomogram environment, a model of the CorM filament is shown at the positions of the *in situ* filaments. This superimposition shows the good correspondence of filament dimensions and appearance.

The shown CorM filament model was derived identically as described in Fig. S10A.

**Movie S6. FRAP of CorM *in vitro***

FRAP experiment of Alexa488-CorM filaments *in vitro* on SLBs. Images were acquired at 1-second intervals. Scale bar: 10 µm.

**Movie S7. FRAP of ^Δ1-40^CorM *in vitro***

FRAP experiment of Alexa488-^Δ1-40^CorM filaments *in vitro* on SLBs. Images were acquired at 1-second intervals. Scale bar: 10 µm.

**Movie S8. CorM polymerization dynamics *in vitro***

TIRF microscopy of Alexa488-CorM filaments on supported lipid bilayers. Individual filaments can only be tracked during the early stages of polymerization. Images were acquired at 0.15-second intervals. Scale bar: 2 µm.

**Movie S9. CorM dual color filament tracking *in vitro***

TIRF microscopy of Alexa488-CorM filaments with single Cy5-CorM monomers on supported lipid bilayers. Images were acquired at 1-second intervals. Scale bar: 1 µm.

**Movie S10. Live cell time lapse microscopy of mNG-MinC**

*Anabaena* cells expressing mNG-MinC were grown on 1.5% agar pads and images were acquired at 1-minute intervals.

**Movie S11. CorM + MinC *in vitro***

TIRF microscopy of pre-assembled Alexa488-CorM filaments. MinC was added at equimolar concentration after 30 seconds. Images were acquired at 1-second intervals. Scale bar: 5 µm.

**Movie S12. CorM + MinC^S50A+L55D^ *in vitro***

TIRF microscopy of pre-assembled Alexa488-CorM filaments. MinC^S50A+L55D^ was added at equimolar concentration after 30 seconds. Images were acquired at 1-second intervals. Scale bar: 5 µm.

**Movie S13. CorM + ^Δ46-67^MinC *in vitro***

TIRF microscopy of pre-assembled Alexa488-CorM filaments. ^Δ46-67^MinC was added at equimolar concentration after 30 seconds. Images were acquired at 1-second intervals. Scale bar: 5 µm.

**Movie S14. CorM + buffer ctrl. *in vitro***

TIRF microscopy of pre-assembled Alexa488-CorM filaments. Buffer was added at a volume equivalent to that of MinC after 30 seconds. Images were acquired at 1-second intervals. Scale bar: 5 µm.

## Notes

### Competing Interest Statement

The authors have declared no competing interest.

### Summary of Updates

Our revised manuscript includes: 1.Clear complementation of the ΔcorMR mutant phenotype, using new constructs expressing corMR under alternative promoters that restore wild-type-like morphology. Together with additional single-gene deletion mutants that recapitulate the ΔcorMR phenotype, these data demonstrate the essential roles of CorM and CorR in maintaining cell shape. 2.New immunofluorescence microscopy of wild-type Anabaena using a newly generated anti-CorM antibody. These experiments independently confirm the cortical localization of mNG-CorM observed during live-cell imaging, demonstrating that the fluorescent fusions faithfully reflect the native protein distribution. 3.New cryo-electron tomography of FIB-milled Anabaena cells, revealing cortical filaments with dimensions matching those of CorM filaments assembled in vitro, thereby providing direct structural evidence for CorM filaments in vivo. Together, these new data provide independent and convergent evidence that the CorMR system forms a cortical cytoskeletal network critical for cell shape maintenance in multicellular cyanobacteria.

## References

1. J. Wagstaff, J. Löwe, Prokaryotic cytoskeletons: protein filaments organizing small cells. Nat. Rev. Microbiol. 16, 187–201 (2018).

2. J. Møller-Jensen, J. Borch, M. Dam, R. B. Jensen, P. Roepstorff, K. Gerdes, Bacterial Mitosis: ParM of Plasmid R1 Moves Plasmid DNA by an Actin-like Insertional Polymerization Mechanism. Mol. Cell 12, 1477–1487 (2003).

3. E. Bi, J. Lutkenhaus, FtsZ ring structure associated with division in Escherichia coli. Nature 354, 161–164 (1991).

4. M. Loose, T. J. Mitchison, The bacterial cell division proteins ftsA and ftsZ self-organize into dynamic cytoskeletal patterns. Nat. Cell Biol. 16, 38–46 (2014).

5. D. Shiomi, W. Margolin, Dimerization or oligomerization of the actin-like FtsA protein enhances the integrity of the cytokinetic Z ring. Mol. Microbiol. 66, 1396–1415 (2007).

6. E. C. Garner, R. Bernard, W. Wang, X. Zhuang, D. Z. Rudner, T. Mitchison, Coupled, circumferential motions of the cell wall synthesis machinery and MreB filaments in B. subtilis. Science (80-.). 333, 222–225 (2011).

7. B. Ramm, T. Heermann, P. Schwille, The E. coli MinCDE system in the regulation of protein patterns and gradients. Cell. Mol. Life Sci. 76, 4245–4273 (2019).

8. B. Wickstead, K. Gull, The evolution of the cytoskeleton. J. Cell Biol. 194, 513–525 (2011).

9. E. Flores, A. Herrero, The cyanobacteria: morphological diversity in a photoautotrophic lifestyle. (2014).

10. R. Rippka, R. Y. Stanier, J. Deruelles, M. Herdman, J. B. Waterbury, Generic Assignments, Strain Histories and Properties of Pure Cultures of Cyanobacteria. Microbiology 111, 1–61 (1979).

11. A. Herrero, J. Stavans, E. Flores, The multicellular nature of filamentous heterocyst-forming cyanobacteria. FEMS Microbiol. Rev. 40, 831–854 (2016).

12. A. K. Kieninger, I. Maldener, Cell–cell communication through septal junctions in filamentous cyanobacteria. Curr. Opin. Microbiol. 61, 35–41 (2021).

13. G. L. Weiss, A.-K. Kieninger, I. Maldener, K. Forchhammer, M. Pilhofer, Structure and Function of a Bacterial Gap Junction Analog. Cell 178, 374–384.e15 (2019).

14. S. Y. Miyagishima, P. P. Wolk, K. W. Osteryoung, Identification of cyanobacterial cell division genes by comparative and mutational analyses. Mol. Microbiol. 56, 126–143 (2005).

15. B. L. Springstein, J. Weissenbach, R. Koch, F. Stücker, K. Stucken, The role of the cytoskeletal proteins MreB and FtsZ in multicellular cyanobacteria. FEBS Open Bio 10, 2510–2531 (2020).

16. A. Naha, D. P. Haeusser, W. Margolin, Anchors: A way for FtsZ filaments to stay membrane bound. Mol. Microbiol. 120, 525–538 (2023).

17. S. Camargo, S. Picossi, L. Corrales-Guerrero, A. Valladares, S. Arévalo, A. Herrero, ZipN is an essential FtsZ membrane tether and contributes to the septal localization of SepJ in the filamentous cyanobacterium Anabaena. Sci. Rep. 9, 1–15 (2019).

18. J. S. MacCready, J. Schossau, K. W. Osteryoung, D. C. Ducat, Robust Min-system oscillation in the presence of internal photosynthetic membranes in cyanobacteria. Mol. Microbiol. 103, 483–503 (2017).

19. B. L. Springstein, D. J. Nürnberg, G. L. Weiss, M. Pilhofer, K. Stucken, Structural determinants and their role in cyanobacterial morphogenesis. Life 10, 355 (2020).

20. C. Velázquez-Suárez, A. Valladares, I. Luque, A. Herrero, The Role of Mre Factors and Cell Division in Peptidoglycan Growth in the Multicellular Cyanobacterium Anabaena. MBio 13 (2022).

21. B. Hu, G. Yang, W. Zhao, Y. Zhang, J. Zhao, MreB is important for cell shape but not for chromosome segregation of the filamentous cyanobacterium Anabaena sp. PCC 7120. Mol. Microbiol. 63, 1640–1652 (2007).

22. C. Velázquez-Suárez, I. Luque, A. Herrero, The Inorganic Nutrient Regime and the mre Genes Regulate Cell and Filament Size and Morphology in the Phototrophic Multicellular Bacterium Anabaena. mSphere 5, e00747–20 (2020).

23. M. Griese, C. Lange, J. Soppa, Ploidy in cyanobacteria. FEMS Microbiol. Lett. 323, 124– 131 (2011).

24. J. Salje, P. Gayathri, J. Löwe, The ParMRC system: Molecular mechanisms of plasmid segregation by actin-like filaments. Nat. Rev. Microbiol. 8, 683–692 (2010).

25. P. Gayathri, T. Fujii, J. Møller-Jensen, F. Van Den Ent, K. Namba, J. Löwe, A bipolar spindle of antiparallel ParM filaments drives bacterial plasmid segregation. Science (80-.). 338, 1334–1337 (2012).

26. F. Van den Ent, J. Møller-Jensen, L. A. Amos, K. Gerdes, J. Löwe, F-actin-like filaments formed by plasmid segregation protein ParM. EMBO J. 21, 6935–6943 (2002).

27. T. A. M. Bharat, G. N. Murshudov, C. Sachse, J. Löwe, Structures of actin-like ParM filaments show architecture of plasmid-segregating spindles. Nature 523, 106–110 (2015).

28. J. Møller-Jensen, R. B. Jensen, J. Löwe, K. Gerdes, Prokaryotic DNA segregation by an actin-like filament. EMBO J. 21, 3119–3127 (2002).

29. S. Ali, A. Koh, D. Popp, K. Tanaka, Y. Kitaoku, N. Miyazaki, K. Iwasaki, K. Mitsuoka, R. C. Robinson, A. Narita, Bacterial genome encoded ParMs. J. Biol. Chem., 110351 (2025).

30. K. Gerdes, J. Moller-Jensen, R. B. Jensen, Plasmid and chromosome partitioning: Surprises from phylogeny. Mol. Microbiol. 37, 455–466 (2002).

31. J. Møller-Jensen, S. Ringgaard, C. P. Mercogliano, K. Gerdes, J. Löwe, Structural analysis of the ParR/parC plasmid partition complex. EMBO J. 26, 4413–4422 (2007).

32. A. Breüner, R. B. Jensen, M. Dam, S. Pedersen, K. Gerdes, The centromere-like parC locus of plasmid R1. Mol. Microbiol. 20, 581–592 (1996).

33. M. Dam, K. Gerdes, Partitioning of plasmid R1 Ten direct repeats flanking the parA promoter constitute a centromere-like partition site parC, that expresses incompatibility. J. Mol. Biol. 236, 1289–1298 (1994).

34. S. Jiang, A. Narita, D. Popp, U. Ghoshdastider, L. J. Lee, R. Srinivasan, M. K. Balasubramanian, T. Oda, F. Koh, M. Larsson, R. C. Robinson, Novel actin filaments from Bacillus thuringiensis form nanotubules for plasmid DNA segregation. Proc. Natl. Acad. Sci. U. S. A. 113, E1200–E1205 (2016).

35. A. J. Brzoska, S. O. Jensen, D. A. Barton, D. S. Davies, R. L. Overall, R. A. Skurray, N. Firth, Dynamic filament formation by a divergent bacterial actin-like ParM protein. PLoS One 11, 1–14 (2016).

36. E. R. Schreiter, C. L. Drennan, Ribbon-helix-helix transcription factors: Variations on a theme. Nat. Rev. Microbiol. 5, 710–720 (2007).

37. J. Salje, J. Löwe, Bacterial actin: Architecture of the ParMRC plasmid DNA partitioning complex. EMBO J. 27, 2230–2238 (2008).

38. A. Orlova, E. C. Garner, V. E. Galkin, J. Heuser, R. D. Mullins, E. H. Egelman, The structure of bacterial ParM filaments. Nat. Struct. Mol. Biol. 14, 921–926 (2007).

39. P. S. Lee, A. D. Grossman, The chromosome partitioning proteins Soj (ParA) and Spo0J (ParB) contribute to accurate chromosome partitioning, separation of replicated sister origins, and regulation of replication initiation in Bacillus subtilis. Mol. Microbiol. 60, 853–869 (2006).

40. D. A. Mohl, J. W. Gober, Cell Cycle–Dependent Polar Localization of Chromosome Partitioning Proteins in Caulobacter crescentus. Cell 88, 675–684 (1997).

41. H. Niki, T. Ogura, C. Ichinose, H. Mori, B. Ezaki, A. Jaffe, Chromosome partitioning in Escherichia coli: novel mutants producing anucleate cells. J. Bacteriol. 171, 1496–1505 (1989).

42. J. Svoboda, B. Cisneros, B. Philmus, Evaluation of inducible promoter-riboswitch constructs for heterologous protein expression in the cyanobacterial species Anabaena sp. PCC 7120. Synth. Biol. 6, 1–8 (2021).

43. Y. Yang, X.-Z. Huang, L. Wang, V. Risoul, C.-C. Zhang, W.-L. Chen, Phenotypic variation caused by variation in the relative copy number of pDU1-based plasmids expressing the GAF domain of Pkn41 or Pkn42 in Anabaena sp. PCC 7120. Res. Microbiol. 164, 127–135 (2013).

44. B. Bishé, A. Taton, J. W. Golden, Modification of RSF1010-Based Broad-Host-Range Plasmids for Improved Conjugation and Cyanobacterial Bioprospecting. iScience 20, 216– 228 (2019).

45. J. Ungerer, H. B. Pakrasi, Cpf1 Is A Versatile Tool for CRISPR Genome Editing Across Diverse Species of Cyanobacteria. Sci. Rep. 6, 1–9 (2016).

46. T. C. Niu, G. M. Lin, L. R. Xie, Z. Q. Wang, W. Y. Xing, J. Y. Zhang, C. C. Zhang, Expanding the Potential of CRISPR-Cpf1-Based Genome Editing Technology in the Cyanobacterium Anabaena PCC 7120. ACS Synth. Biol. 8, 170–180 (2019).

47. C. S. Campbell, R. D. Mullins, In vivo visualization of type II plasmid segregation: Bacterial actin filaments pushing plasmids. J. Cell Biol. 179, 1059–1066 (2007).

48. M. Á. Rubio, M. Napolitano, J. A. G. Ochoa De Alda, J. Santamaría-Gómez, C. J. Patterson, A. W. Foster, R. Bru-Martínez, N. J. Robinson, I. Luque, Trans-oligomerization of duplicated aminoacyl-tRNA synthetases maintains genetic code fidelity under stress. Nucleic Acids Res. 43, 9905–9917 (2015).

49. E. C. Garner, C. S. Campbell, R. Dyce Mullins, Dynamic Instability in a DNA-Segregating Prokaryotic Actin Homolog. Science (80-.). 306, 1021–1025 (2004).

50. E. C. Garner, C. S. Campbell, D. B. Weibel, R. D. Mullins, Reconstitution of DNA Segregation Driven by Assembly of a Prokaryotic Actin Homolog. Science (80-.). 315, 1270–1274 (2007).

51. N. Sapay, Y. Guermeur, G. Deléage, Prediction of amphipathic in-plane membrane anchors in monotopic proteins using a SVM classifier. BMC Bioinformatics 7, 1–11 (2006).

52. E. Krissinel, K. Henrick, Inference of Macromolecular Assemblies from Crystalline State. J. Mol. Biol. 372, 774–797 (2007).

53. T. Müller, F. M. Kleusberg, K. Roganowicz, G. L. Weiss, M. Coles, K. A. Selim, The Ca2+-binding protein CSE links Ca2+-signaling with cell-cell communication in multicellular cyanobacteria. bioRxiv, 2025.03.26.645587 (2025).

54. R. Evans, M. O’Neill, A. Pritzel, N. Antropova, A. Senior, T. Green, A. Žídek, R. Bates, S. Blackwell, J. Yim, O. Ronneberger, S. Bodenstein, M. Zielinski, A. Bridgland, A. Potapenko, A. Cowie, K. Tunyasuvunakool, R. Jain, E. Clancy, P. Kohli, J. Jumper, D. Hassabis, Protein complex prediction with AlphaFold-Multimer. bioRxiv, 2021.10.04.463034 (2022).

55. J. Jumper, R. Evans, A. Pritzel, T. Green, M. Figurnov, O. Ronneberger, K. Tunyasuvunakool, R. Bates, A. Žídek, A. Potapenko, A. Bridgland, C. Meyer, S. A. A. Kohl, A. J. Ballard, A. Cowie, B. Romera-Paredes, S. Nikolov, R. Jain, J. Adler, T. Back, S. Petersen, D. Reiman, E. Clancy, M. Zielinski, M. Steinegger, M. Pacholska, T. Berghammer, S. Bodenstein, D. Silver, O. Vinyals, A. W. Senior, K. Kavukcuoglu, P. Kohli, D. Hassabis, Highly accurate protein structure prediction with AlphaFold. Nature 596, 583–589 (2021).

56. P. A. J. de Boer, R. E. Crossley, L. I. Rothfield, A division inhibitor and a topological specificity factor coded for by the minicell locus determine proper placement of the division septum in E. coli. Cell 56, 641–649 (1989).

57. Z. Hu, J. Lutkenhaus, Topological regulation of cell division in Escherichia coli involves rapid pole to pole oscillation of the division inhibitor MinC under the control of MinD and MinE. Mol. Microbiol. 34, 82–90 (1999).

58. D. M. Raskin, P. A. J. De Boer, Rapid pole-to-pole oscillation of a protein required for directing division to the middle of Escherichia coli. Proc. Natl. Acad. Sci. 96, 4971–4976 (1999).

59. T. D. Pollard, Actin and actin-binding proteins. Cold Spring Harb. Perspect. Biol. 8, 1–17 (2016).

60. B. Xue, C. Leyrat, J. M. Grimes, R. C. Robinson, Structural basis of thymosin-β4/profilin exchange leading to actin filament polymerization. Proc. Natl. Acad. Sci. U. S. A. 111, E4596–E4605 (2014).

61. S. Palmgren, M. Vartiainen, P. Lappalainen, Twinfilin, a molecular mailman for actin monomers. J. Cell Sci. 115, 881–886 (2002).

62. P. A. Curmi, S. S. L. Andersen, S. Lachkar, O. Gavet, E. Karsenti, M. Knossow, A. Sobel, The stathmin/tubulin interaction in vitro. J. Biol. Chem. 272, 25029–25036 (1997).

63. Z. Hu, A. Mukherjee, S. Pichoff, J. Lutkenhaus, The MinC component of the division site selection system in Escherichia coli interacts with FtsZ to prevent polymerization. Proc. Natl. Acad. Sci. U. S. A. 96, 14819–14824 (1999).

64. A. R. Paredez, C. R. Somerville, D. W. Ehrhardt, Visualization of Cellulose Synthase Demonstrates Functional Association with Microtubules. Science (80-.). 312, 1491–1495 (2006).

65. K. Sugimoto, R. E. Williamson, G. O. Wasteneys, New techniques enable comparative analysis of microtubule orientation, wall texture, and growth rate in intact roots of Arabidopsis. Plant Physiol. 124, 1493–1506 (2000).

66. A. R. Paredez, S. Persson, D. W. Ehrhardt, C. R. Somerville, Genetic evidence that cellulose synthase activity influences microtubule cortical array organization. Plant Physiol. 147, 1723–1734 (2008).

67. D. F. Savage, B. Afonso, A. H. Chen, P. A. Silver, Spatially Ordered Dynamics of the Bacterial Carbon Fixation Machinery. Science (80-.). 327, 1258–1261 (2010).

68. I. H. Jain, V. Vijayan, E. K. O’Shea, Spatial ordering of chromosomes enhances the fidelity of chromosome partitioning in cyanobacteria. Proc. Natl. Acad. Sci. U. S. A. 109, 13638–43 (2012).

69. M. Wachi, M. Doi, S. Tamaki, W. Park, S. Nakajima-Iijima, M. Matsuhashi, Mutant isolation and molecular cloning of mre genes, which determine cell shape, sensitivity to mecillinam, and amount of penicillin-binding proteins in Escherichia coli. J. Bacteriol. 169, 4935–4940 (1987).

70. H. Shi, B. P. Bratton, Z. Gitai, K. C. Huang, How to Build a Bacterial Cell: MreB as the Foreman of E. coli Construction. Cell 172, 1294–1305 (2018).

71. F. van den Ent, T. Izoré, T. A. M. Bharat, C. M. Johnson, J. Löwe, Bacterial actin MreB forms antiparallel double filaments. Elife 2014, 1–22 (2014).

72. S. van Teeffelen, S. Wang, L. Furchtgott, K. C. Huang, N. S. Wingreen, J. W. Shaevitz, Z. Gitai, The bacterial actin MreB rotates, and rotation depends on cell-wall assembly. Proc. Natl. Acad. Sci. 108, 15822 LP – 15827 (2011).

73. A. Mukherjee, J. Lutkenhaus, Dynamic assembly of FtsZ regulated by GTP hydrolysis. EMBO J. 17, 462–469 (1998).

74. M. F. Carlier, D. Pantaloni, E. D. Korn, Evidence for an ATP cap at the ends of actin filaments and its regulation of the F-actin steady state. J. Biol. Chem. 259, 9983–9986 (1984).

75. T. Mitchison, M. Kirschner, Dynamic instability of microtubule growth. Nature 312, 237– 242 (1984).

76. T. D. Pollard, R. D. Goldman, Overview of the cytoskeleton from an evolutionary perspective. Cold Spring Harb. Perspect. Biol. 10, a030288 (2018).

77. J. Abramson, J. Adler, J. Dunger, R. Evans, T. Green, A. Pritzel, O. Ronneberger, L. Willmore, A. J. Ballard, J. Bambrick, S. W. Bodenstein, D. A. Evans, C. C. Hung, M. O’Neill, D. Reiman, K. Tunyasuvunakool, Z. Wu, A. Žemgulytė, E. Arvaniti, C. Beattie, O. Bertolli, A. Bridgland, A. Cherepanov, M. Congreve, A. I. Cowen-Rivers, A. Cowie, M. Figurnov, F. B. Fuchs, H. Gladman, R. Jain, Y. A. Khan, C. M. R. Low, K. Perlin, A. Potapenko, P. Savy, S. Singh, A. Stecula, A. Thillaisundaram, C. Tong, S. Yakneen, E. D. Zhong, M. Zielinski, A. Žídek, V. Bapst, P. Kohli, M. Jaderberg, D. Hassabis, J. M. Jumper, Accurate structure prediction of biomolecular interactions with AlphaFold 3. Nature 630, 493–500 (2024).

78. E. C. Meng, T. D. Goddard, E. F. Pettersen, G. S. Couch, Z. J. Pearson, J. H. Morris, T. E. Ferrin, UCSF ChimeraX: Tools for structure building and analysis. Protein Sci. 32, e4792 (2023).

79. A. Valladares, S. Picossi, L. Corrales-Guerrero, A. Herrero, The role of SepF in cell division and diazotrophic growth in the multicellular cyanobacterium Anabaena sp. strain PCC 7120. Microbiol. Res. 277, 127489 (2023).

80. G. Karimova, M. Davi, D. Ladant, The β-lactam resistance protein Blr, a small membrane polypeptide, is a component of the Escherichia coli cell division machinery. J. Bacteriol. 194, 5576–5588 (2012).

81. K. Kartasalo, R. P. Pölönen, M. Ojala, J. Rasku, J. Lekkala, K. Aalto-Setälä, P. Kallio, CytoSpectre: A tool for spectral analysis of oriented structures on cellular and subcellular levels. BMC Bioinformatics 16, 1–23 (2015).

82. C. Sun, B. Gonzalez, W. Jiang, Helical indexing in real space. Sci. Rep. 12, 8162 (2022).

83. T. I. Croll, ISOLDE: a physically realistic environment for model building into low-resolution electron-density maps. Biol. Crystallogr. 74, 519–530 (2018).

84. C. J. Williams, J. J. Headd, N. W. Moriarty, M. G. Prisant, L. L. Videau, L. N. Deis, V. Verma, D. A. Keedy, B. J. Hintze, V. B. Chen, MolProbity: more and better reference data for improved all-atom structure validation. Protein Sci. 27, 293–315 (2018).

85. S. Klumpe, H. K. H. Fung, S. K. Goetz, I. Zagoriy, B. Hampoelz, X. Zhang, P. S. Erdmann, J. Baumbach, C. W. Müller, M. Beck, A modular platform for automated cryo-FIB workflows. Elife 10, e70506 (2021).

86. J. M. Medeiros, D. Böck, G. L. Weiss, R. Kooger, R. A. Wepf, M. Pilhofer, Robust workflow and instrumentation for cryo-focused ion beam milling of samples for electron cryotomography. Ultramicroscopy 190, 1–11 (2018).

87. D. N. Mastronarde, SerialEM: a program for automated tilt series acquisition on Tecnai microscopes using prediction of specimen position. Microsc. Microanal. 9, 1182–1183 (2003).

88. W. J. H. Hagen, W. Wan, J. A. G. Briggs, Implementation of a cryo-electron tomography tilt-scheme optimized for high resolution subtomogram averaging. J. Struct. Biol. 197, 191–198 (2017).

89. F. Eisenstein, H. Yanagisawa, H. Kashihara, M. Kikkawa, S. Tsukita, R. Danev, Parallel cryo electron tomography on in situ lamellae. Nat. Methods 20, 131–138 (2023).

90. S. Zheng, G. Wolff, G. Greenan, Z. Chen, F. G. A. Faas, M. Bárcena, A. J. Koster, Y. Cheng, D. A. Agard, AreTomo: An integrated software package for automated marker-free, motion-corrected cryo-electron tomographic alignment and reconstruction. J. Struct. Biol. X 6, 100068 (2022).

91. J. R. Kremer, D. N. Mastronarde, J. R. McIntosh, Computer visualization of three-dimensional image data using IMOD. J. Struct. Biol. 116, 71–76 (1996).

92. D. N. Mastronarde, Correction for non-perpendicularity of beam and tilt axis in tomographic reconstructions with the IMOD package. J. Microsc. 230, 212–217 (2008).

93. Y.-T. Liu, H. Zhang, H. Wang, C.-L. Tao, G.-Q. Bi, Z. H. Zhou, Isotropic reconstruction for electron tomography with deep learning. Nat. Commun. 13, 6482 (2022).

94. E. W. Sayers, E. E. Bolton, J. R. Brister, K. Canese, J. Chan, D. C. Comeau, C. M. Farrell, M. Feldgarden, A. M. Fine, K. Funk, E. Hatcher, S. Kannan, C. Kelly, S. Kim, W. Klimke, M. J. Landrum, S. Lathrop, Z. Lu, T. L. Madden, A. Malheiro, A. Marchler-Bauer, T. D. Murphy, L. Phan, S. Pujar, S. H. Rangwala, V. A. Schneider, T. Tse, J. Wang, J. Ye, B. W. Trawick, K. D. Pruitt, S. T. Sherry, Database resources of the National Center for Biotechnology Information in 2023. Nucleic Acids Res. 51, D29–D38 (2023).

95. S. R. Eddy, Accelerated Profile HMM Searches. PLOS Comput. Biol. 7, e1002195 (2011).

96. K. Katoh, D. M. Standley, MAFFT multiple sequence alignment software version 7: Improvements in performance and usability. Mol. Biol. Evol. 30, 772–780 (2013).

97. T. Goldfarb, V. K. Kodali, S. Pujar, V. Brover, B. Robbertse, C. M. Farrell, D.-H. Oh, A. Astashyn, O. Ermolaeva, D. Haddad, W. Hlavina, J. Hoffman, J. D. Jackson, V. S. Joardar, D. Kristensen, P. Masterson, K. M. McGarvey, R. McVeigh, E. Mozes, M. R. Murphy, S. S. Schafer, A. Souvorov, B. Spurrier, P. K. Strope, H. Sun, A. R. Vatsan, C. Wallin, D. Webb, J. R. Brister, E. Hatcher, A. Kimchi, W. Klimke, A. Marchler-Bauer, K. D. Pruitt, F. Thibaud-Nissen, T. D. Murphy, NCBI RefSeq: reference sequence standards through 25 years of curation and annotation. Nucleic Acids Res. 53, D243–D257 (2025).

98. D. Megrian, N. Taib, A. L. Jaffe, J. F. Banfield, S. Gribaldo, Ancient origin and constrained evolution of the division and cell wall gene cluster in Bacteria. Nat. Microbiol. 7, 2114–2127 (2022).

99. M. AR, C. MS, F. W, NetWheels: A Web Application to Create High Quality Peptide Helical Wheel and Net Projections. J. Bioinforma. Syst. Biol. 7, 98–100 (2024).

100. N. Van Hilten, N. Verwei, J. Methorst, C. Nase, A. Bernatavicius, H. J. Risselada, PMIpred: a physics-informed web server for quantitative protein-membrane interaction prediction. Bioinformatics 40, 1–10 (2024).

101. S. Capella-Gutiérrez, J. M. Silla-Martínez, T. Gabaldón, trimAl: a tool for automated alignment trimming in large-scale phylogenetic analyses. Bioinformatics 25, 1972–1973 (2009).

102. A. M. Waterhouse, J. B. Procter, D. M. A. Martin, M. Clamp, G. J. Barton, Jalview Version 2—a multiple sequence alignment editor and analysis workbench. Bioinformatics 25, 1189–1191 (2009).

103. G. E. Crooks, G. Hon, J.-M. Chandonia, S. E. Brenner, WebLogo: a sequence logo generator. Genome Res. 14, 1188–1190 (2004).

104. E. F. Pettersen, T. D. Goddard, C. C. Huang, E. C. Meng, G. S. Couch, T. I. Croll, J. H. Morris, T. E. Ferrin, UCSF ChimeraX: Structure visualization for researchers, educators, and developers. Protein Sci. 30, 70–82 (2021).

105. J. Pei, N. V Grishin, AL2CO: calculation of positional conservation in a protein sequence alignment. Bioinformatics 17, 700–712 (2001).

106. B. Q. Minh, H. A. Schmidt, O. Chernomor, D. Schrempf, M. D. Woodhams, A. von Haeseler, R. Lanfear, IQ-TREE 2: New Models and Efficient Methods for Phylogenetic Inference in the Genomic Era. Mol. Biol. Evol. 37, 1530–1534 (2020).

107. S. Kalyaanamoorthy, B. Q. Minh, T. K. F. Wong, A. von Haeseler, L. S. Jermiin, ModelFinder: fast model selection for accurate phylogenetic estimates. Nat. Methods 14, 587–589 (2017).

108. D. T. Hoang, O. Chernomor, A. von Haeseler, B. Q. Minh, L. S. Vinh, UFBoot2: Improving the Ultrafast Bootstrap Approximation. Mol. Biol. Evol. 35, 518–522 (2018).

109. I. Letunic, P. Bork, Interactive Tree Of Life (iTOL) v5: an online tool for phylogenetic tree display and annotation. Nucleic Acids Res. 49, W293–W296 (2021).

110. J. Schindelin, I. Arganda-Carreras, E. Frise, V. Kaynig, M. Longair, T. Pietzsch, S. Preibisch, C. Rueden, S. Saalfeld, B. Schmid, J. Y. Tinevez, D. J. White, V. Hartenstein, K. Eliceiri, P. Tomancak, A. Cardona, Fiji: An open-source platform for biological-image analysis. (2012). 10.1038/nmeth.2019.

111. J. M. Lotthammer, G. M. Ginell, D. Griffith, R. J. Emenecker, A. S. Holehouse, Direct prediction of intrinsically disordered protein conformational properties from sequence. Nat. Methods 21, 465–476 (2024).

112. C. P. Wolk, Y. Cai, L. Cardemil, E. Flores, B. Hohn, M. Murry, G. Schmetterer, B. Schrautemeier, R. Wilson, Isolation and complementation of mutants of Anabaena sp. strain PCC 7120 unable to grow aerobically on dinitrogen. J. Bacteriol. 170, 1239–1244 (1988).

113. F. O. Bendezú, C. A. Hale, T. G. Bernhardt, P. A. J. De Boer, RodZ (YfgA) is required for proper assembly of the MreB actin cytoskeleton and cell shape in E. coli. EMBO J. 28, 193–204 (2009).

